# Properties and unbiased estimation of *F*- and *D*-statistics in samples containing related and inbred individuals

**DOI:** 10.1101/2020.11.20.391367

**Authors:** Mehreen R. Mughal, Michael DeGiorgio

**Affiliations:** Bioinformatics and Genomics at the Huck Institutes of the Life Sciences, Pennsylvania State University, University Park, PA 16802, USA; Department of Computer and Electrical Engineering and Computer Science, Florida Atlantic University, Boca Raton, FL 33431, USA

## Abstract

The Patterson *F*- and *D*-statistics are commonly-used measures for quantifying population relationships and for testing hypotheses about demographic history. These statistics make use of allele frequency information across populations to infer different aspects of population history, such as population structure and introgression events. Inclusion of related or inbred individuals can bias such statistics, which may often lead to the filtering of such individuals. Here we derive statistical properties of the *F*- and *D*-statistics, including their biases due to finite sample size or the inclusion of related or inbred individuals, their variances, and their corresponding mean squared errors. Moreover, for those statistics that are biased, we develop unbiased estimators and evaluate the variances of these new quantities. Comparisons of the new unbiased statistics to the originals demonstrates that our newly-derived statistics often have lower error across a wide population parameter space. Furthermore, we apply these unbiased estimators using several global human populations with the inclusion of related individuals to highlight their application on an empirical dataset. Finally, we implement these unbiased estimators in open-source software package `funbiased` for easy application by the scientific community.

## Introduction

The recently introduced *F*- and *D*-statistics (Huson et al., 2005; Kulathinal et al., 2009; Reich et al., 2009; Green et al., 2010; Patterson et al., 2012) have transformed the way geneticists measure population differentiation. These statistics have been instrumental in many major recent discoveries, including testing which Neanderthal populations are closest to the populations that admixed with modern humans (Hajdinjak et al., 2018), and detecting which population is likely the admixing source for European admixture in modern Ethiopian populations (Molinaro et al., 2019). Iterating through different combinations of populations using the *F*_4_- and *D*-statistics has allowed reconstruction of population histories in diverse groups such as Native Americans and South Asians (Reich et al., 2012; Moorjani et al., 2013). In addition, The *D*-statistics have been used extensively to provide evidence of introgression and hybridization among species of *Drosophila* fruit flies and Heliconius butterflies (Martin et al., 2014; Turissini and Matute, 2017).

In many cases, however, the populations tested by these statistics are small, and proper random sampling may include data from related individuals. It is common to remove one or more of the relatives from a group of related individuals because including them might provide a bias in the value of a particular statistic being measured (Rosenberg, 2006; DeGiorgio and Rosenberg, 2009; DeGiorgio et al., 2010; Waples and Anderson, 2017; Harris and DeGiorgio, 2017a). For this reason we explore whether the current estimators for these statistics are biased with the inclusion of related or inbred individuals and if so, then develop unbiased estimators under such scenarios.

These statistics are flexible and relatively simple to compute, as they measure genetic drift along branches of a population tree by contrasting allele frequencies between different combinations of populations. Using allele frequency data from two, three, or four populations, these statistics measure shared variation along specific branches of the tree relating the populations. We begin by providing intuitive descriptions and formal definitions of each of the *F*- and *D*-statistics that we evaluate. Specifically, consider that we have allele frequency data at *J* biallelic loci from each of four populations, denoted *A*, *B*, *C*, and *D*. We denote the frequencies of the reference allele at locus *j* as *a*_*j*_, *b*_*j*_, *c*_*j*_, and *d*_*j*_ in populations *A*, *B*, *C*, and *D*, respectively. We define each of the *F*- and *D*-statistics as they are in Reich et al. (2009) and Patterson et al. (2012).

We first examine the *F*_2_ statistic, which measures the amount of genetic drift separating a pair of populations, and is thus a test for differentiation between them, and is akin to the widely-used fixation index *F*_ST_ (Weir and Cockerham, 1984). For a pair of populations *A* and *B*, we define the *F*_2_ statistic as

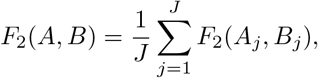

where for locus *j*

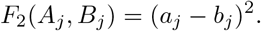

It is clear from this definition that *F*_2_ takes values between zero, when the populations have identical allele frequencies, and one, when the populations are fixed for different alleles (Figure 1).

**Figure 1:**
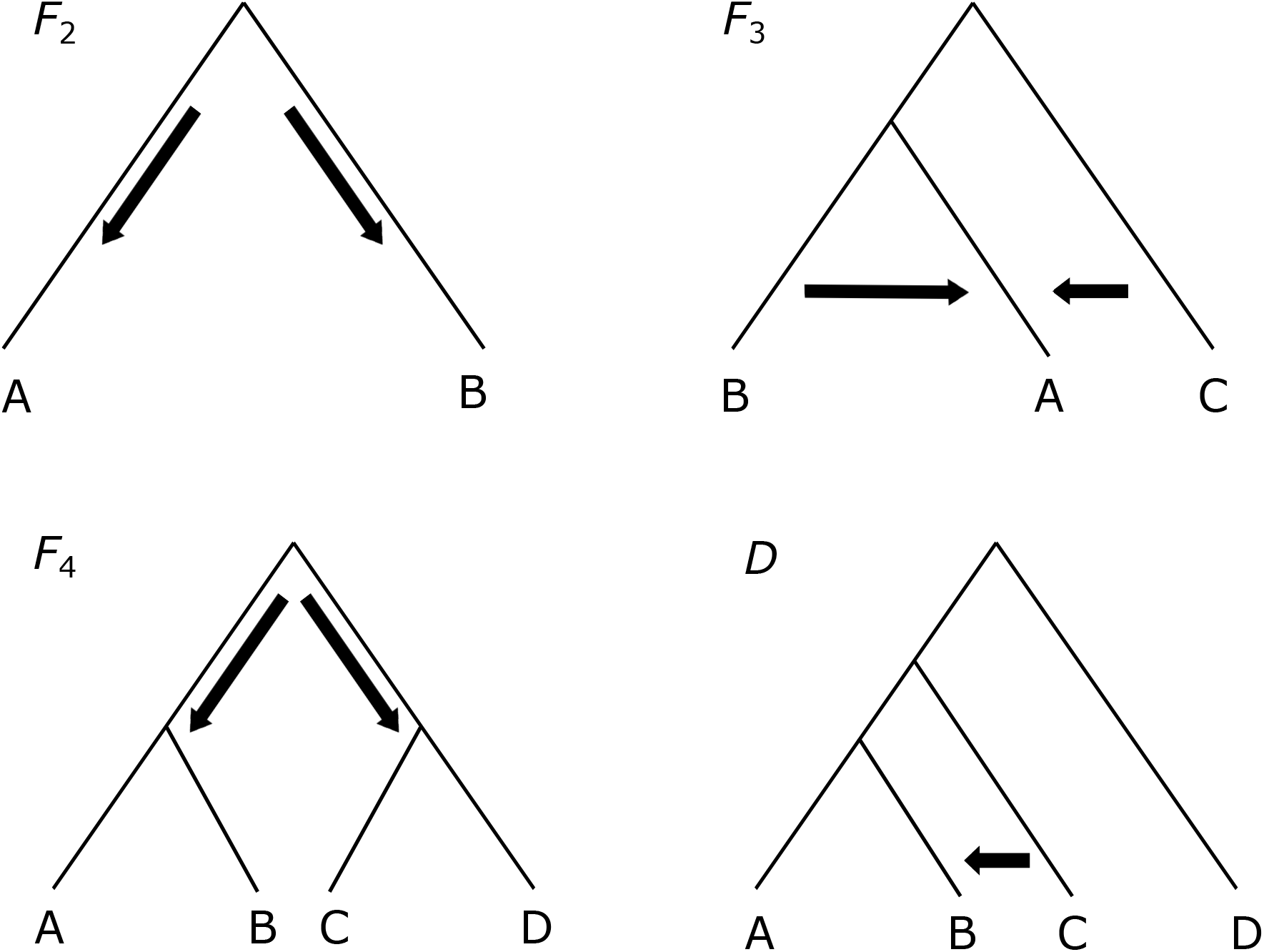
Trees showing the different relationships the *F*- and *D*-statistics are designed to test. *F*_2_(*A, B*) can test the differentiation of two populations *A* and *B*. *F*_3_(*A; B, C*) can test for introgression or relatedness between populations A and B or populations *A* and *C*. *F*_4_(*A, B*; *C, D*) can test the hypothesis of whether two populations are closer to each other than they are to two other populations, in this case are *A* and *B* closer to each other than they are to *C* and *D*. The *D*(*A, B, C, D*) statistic can test whether there has been admixture between population *C* and either populations *A* or *B*.

The *F*_3_ statistic employs allele frequencies from three populations, and measures the amount of genetic drift along the branch leading to a target population, given allele frequency data from two reference populations. For a target population *A* and two reference populations *B* and *C*, we define the *F*_3_ statistic as

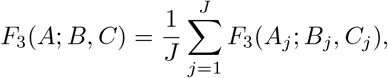

where for locus *j*

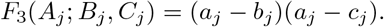

Because it measures genetic drift along a branch leading to a target population, its value is expected to be non-negative. However, an interesting property of the *F*_3_ statistic is that it can be negative if the target population experienced admixture, and therefore a negative value directly indicates admixture in the history of the target population (Reich et al., 2009; Patterson et al., 2012). However, though *F*_3_ can detect admixture if its value is negative, admixture is not guaranteed to lead to negative values (Reich et al., 2009; Patterson et al., 2012), and it is therefore an inconclusive test for admixture if *F*_3_ is non-negative. Moreover, because loci with higher minor allele frequencies may affect *F*_3_ more than loci with lower minor allele frequencies, the *F*_3_ statistic is sometimes normalized (Reich et al., 2009; Patterson et al., 2012) based on levels of diversity of the target population. Formally, this normalized *F*_3_ statistic has definition

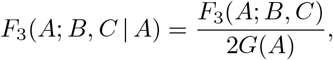

where we define for population *P* (here *P* = *A*)

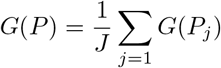

such that for locus *j*

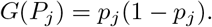

The *F*_4_ statistic, on the other hand, is a test of “treeness” among a set of four populations, examining whether the unrooted tree relating four populations is supported by the allele frequencies within the set of populations. For a pair of sister populations *A* and *B* and a pair of sister populations *C* and *D*, we define the *F*_4_ statistic as

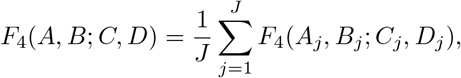

where for locus *j*

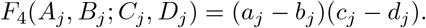

If the unrooted relationship is true, then *F*_4_ takes the value of zero, and is non-zero otherwise. If it is known *a priori* that the unrooted relationship should be true, then a non-zero *F*_4_ statistic can be indicative of admixture, and the sign of the statistic will suggest which set of populations may be violating the assumed unrooted tree topology (Figure 1). As with the *F*_3_ statistic, a normalized version (Reich et al., 2009; Patterson et al., 2012) of the *F*_4_ statistic is sometimes used, with normalization based on the diversity of one of the four populations. Formally, this normalized *F*_4_ statistic has definition

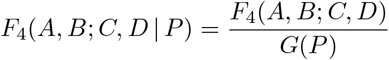

where we normalize by diversity in population *P* ∈ {*A, B, C, D*}.

Finally, the *D*-statistic is a special version of the *F*_4_ statistic that is a test of treeness for a particular asymmetric rooted tree relating four populations, with the tree topology containing a pair of sister populations, together with a close and a distant outgroup population (Figure 1). For sister populations *A* and *B*, close outgroup population *C*, and distant outgroup population *D*, we define the *D* statistic as

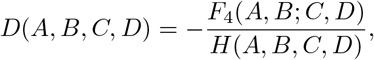

where

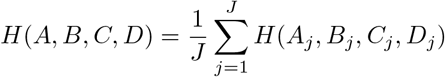

is a normalizing factor to constrain the *D* statistic to take values between negative one and one, such that for locus *j*

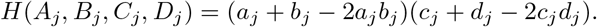

If the rooted relationship is true, then *D* takes the value of zero, and is non-zero otherwise. A non-zero *D* value can be used to detect admixture between the close outgroup population and one of the two sister populations based on its sign (Figure 1).

## Theory

The *F*- and *D*-statistic equations presented in the *Introduction* employ population allele frequencies, and are thus parameters of the set of populations. To estimate them, we first need to build an estimator of allele frequencies based on samples. We denote estimates of the reference allele frequencies at locus *j*, *j* = 1, 2, …, *J*, in populations *A*, *B*, *C*, and *D* by 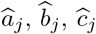, and 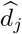, respectively.

As used previously (*e.g.*, McPeek et al., 2004; DeGiorgio and Rosenberg, 2009; DeGiorgio et al., 2010; Harris and DeGiorgio, 2017a), a linear unbiased estimator of population reference allele frequency *p* at a biallelic locus can be defined as

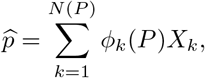

where *N* (*P*) is the number of individuals sampled at the locus, *X_k_* is the frequency of the reference allele in individual *k* at the locus, and *ϕ*_*k*_(*P*) is the weight of individual *k* in population *P* at the locus. McPeek et al. (2004) discussed the impact of various weighting schemes on allele frequency estimation, and Harris and DeGiorgio (2017a) examined the effects of weighting scheme on estimation of expected heterozygosity.

Typical estimators of the *F*- and *D*-statistics are computed as

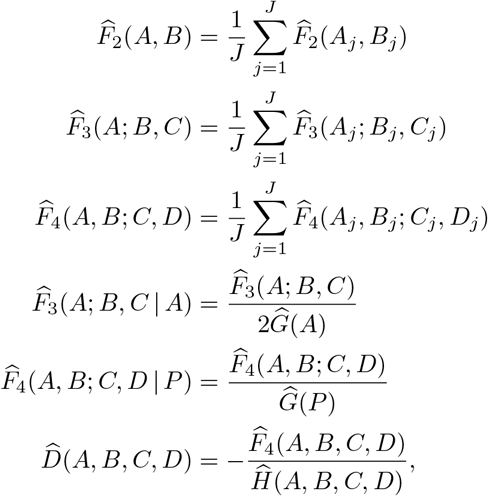

where

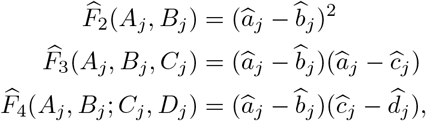

and where

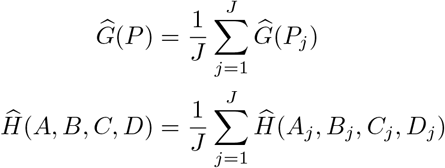

with

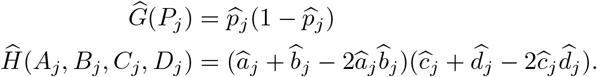

In the following, we discuss properties of these estimators, and where appropriate, develop unbiased estimators for the statistics that are biased either due to finite sample size or due to the inclusion of related or inbred individuals in the sample.

To begin, we define the kinship coefficient Φ_*xy*_ between individuals *x* and *y*, as the probability that a pair of sampled alleles, one from *x* and one from *y* are identical by descent if *x* ≠ *y*, and as the probability that a pair of alleles sampled with replacement from individual *x* are identical by descent if *x* = *y* (Lange, 2002). A pair of unrelated individuals *x* and *y* have kinship coefficient Φ_*xy*_ = 0 (Lange, 2002). Moreover, an individual *x* with ploidy *m_x_* has kinship coefficient Φ_*xx*_ = 1/*m*_*x*_ + (1 − 1/*m*_*x*_)*f*_*x*_ = (1/*m*_*x*_)[1 + (*m*_*x*_ − 1)*f*_*x*_], where *f*_*x*_ is the inbreeding coefficient of individual *x*, and is defined as the probability that a pair of alleles sampled without replacement in individual *x* are identical by descent (DeGiorgio et al., 2010). A non-inbred individual *x* has inbreeding coefficient *f*_*x*_ = 0, and so if *x* is non-inbred, then their kinship coefficient is Φ_*xx*_ = 1/*m*_*x*_. As in DeGiorgio et al. (2010) and Harris and DeGiorgio (2017a), we define the weighted mean kinship coefficients across sets of individuals sampled in population *P* ∈ {*A, B, C, D*} at locus *j* as

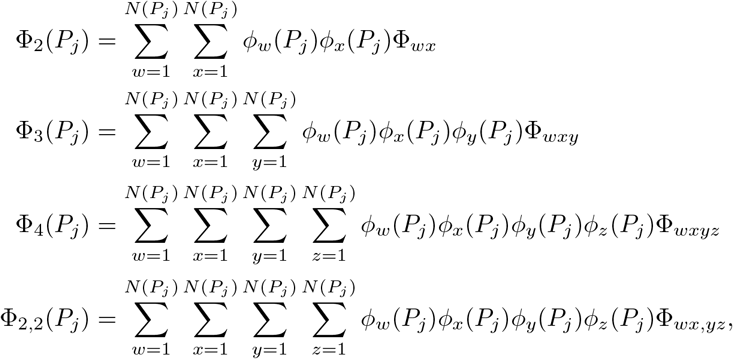

which are the weighted mean kinship coefficients for the *N* (*P_j_*) individuals sampled at locus *j* in population *P* for pairs, triples, quadruples, and pairs of pairs of individuals, respectively. From our definitions of kinship, we know that unrelated individuals have kinship coefficients of zero, but non-inbred individuals still have positive values of their self kinship coefficient, thereby causing the mean kinship coefficients to necessarily be positive quantities. It is for this reason that some *F*-statistic estimators will be biased even without related or inbred individuals, and this bias would be due to finite sample size. For accurate estimates of the drift quantities, it is therefore important to obtain unbiased estimators.

A number of quantities (particularly variances and covariances involving the *F*- and *D*-statistics) will be mathematically complex, as they will involve linear combinations of higher order mean kinship coefficients. For this reason, we follow prior studies (DeGiorgio et al., 2010; Harris and DeGiorgio, 2017a) and make the simplifying assumption that no individual in a sample from population *P* is related to more than one other individual in the sample, such that terms Φ_3_(*P*_*j*_), Φ_4_(*P*_*j*_), Φ_2,2_(*P*_*j*_), and Φ_2_(*P*_*j*_)^2^ negligible to Φ_2_(*P*_*j*_). Moreover, we assume that individuals sampled in different populations are unrelated to each other. Under these assumptions, we approximate a few key results from prior studies (DeGiorgio et al., 2010; Harris and DeGiorgio, 2017a) that will ultimately make derivations easier. Given that 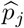 is an estimate of the frequency of a reference allele at locus *j* in population *P*, we have the following expectations (approximate notation when not exact)

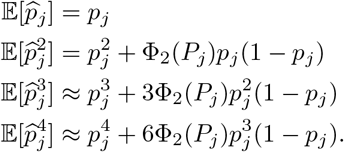

From prior studies (Nei and Roychoudhury, 1974; Weir, 1989; DeGiorgio and Rosenberg, 2009; DeGiorgio et al., 2010; Harris and DeGiorgio, 2017a), we know that 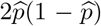 is a downwardly biased estimator of expected heterozygosity at a locus, with the bias due to finite sample size (Nei and Roychoudhury, 1974) and exacerbated by the inclusion of inbred (Weir, 1989) and related (DeGiorgio and Rosenberg, 2009; DeGiorgio et al., 2010; Harris and DeGiorgio, 2017a) individuals in the sample. Based on this definition, 2*G*(*P*) = 2*p*(1 − *p*) is expected heterozygosity, and its estimator 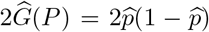 therefore biased. We begin by developing an unbiased estimator for *G*(*P*), as it is a key normalization quantity in the *F*_3_ and *F*_4_ statistics.

### Lemma 1.

Consider *J* polymorphic loci in a population *P* with parametric reference allele frequencies *P*_*j*_ ∈ (0, 1), and suppose we take a random sample of *N* (*P*_*j*_) individuals at locus *j*, some of which may be related or inbred. The estimator 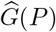 has downward bias

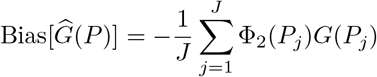

and an unbiased estimator of *G*(*P*) is

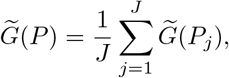

where

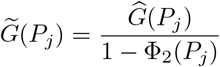

is an unbiased estimator of *G*(*P*_*j*_).

The proof of Lemma 1 is in the *Appendix*. Intuitively though, because 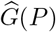 involves the product of frequencies for two alleles drawn from population *P*, there is a chance of having the two alleles being identical by descent by sampling the same allele twice, and is therefore a biased estimator with and without related or inbred individuals

We next consider examining the bias of the estimator 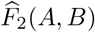. As with 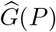, because 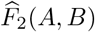 requires sampling two alleles from population *A* and two alleles from population *B*, we find it is biased due to not only finite sample size but also the inclusion of related or inbred individuals. We present the formal result for *F*_2_ next (Proposition 2), and prove the result in the *Appendix*.

### Proposition 2.

Consider *J* polymorphic loci in populations *A* and *B* with respective parametric reference allele frequencies *a*_*j*_, *b*_*j*_ ∈ (0, 1), and suppose we take a random sample of *N* (*P*_*j*_) individuals at locus *j* in population *P* ∈ {*A, B*}, some of which may be related or inbred. The estimate 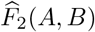 has upward bias

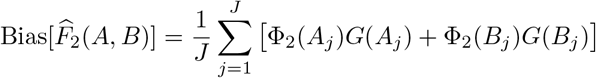

and an unbiased estimator of *F*_2_(*A, B*) is

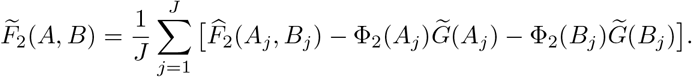

As one can see, the original estimator 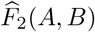 is upwardly biased due to finite sample size and relatedness, and that sampling within both populations *A* and *B* contributes proportionally to this bias. The new unbiased estimator 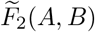 corrects this bias by adjust ng the computation to account for the kinship coefficients and diversity within each population, with the adjustment of diversity using the unbiased estimator 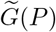 presented in Lemma 1.

Similarly to 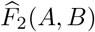, the original estimator 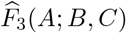 is also upwardly biased because it requires the sampling of two alleles from the target population *A*. We show the formal results for *F*_3_ next (Proposition 3), and prove the result in the *Appendix*.

### Proposition 3.

Consider *J* polymorphic loci in populations *A*, *B*, and *C* with respective parametric reference allele frequencies *a*_*j*_, *b*_*j*_, *c*_*j*_ ∈ (0, 1), and suppose we take a random sample of *N* (*P*_*j*_) individuals at locus *j* in population *P* ∈ {*A, B, C*}, some o which may be related or inbred. The estimate 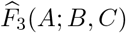 has upward bias

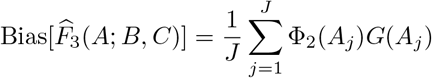

and an unbiased estimator of *F*_3_(*A*; *B, C*) is

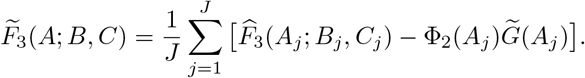

The bias of the original estimator is proportional to the relatedness and diversity within the target population *A*. The new unbiased estimator 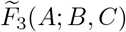 corrects the bias by adjusting the computation to account for the kinship and diversity within the target population, with the adjustment of diversity using the unbiased estimator 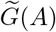. Moreover, it is important to note that the reference populations *B* and *C* do not contribute to bias, as only a single allele is sampled from each of these populations.

Given that 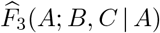 uses the biased estimators 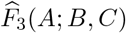 and 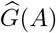 in its definition, we can expect that it would be biased as its component estimators are biased, and these components have different biases that are also in different directions. However, 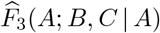 is a ratio estimator, and we can therefore not directly take its expectation to evaluate bias. Instead, we will make some simplifying assumptions and compute the approximate bias of 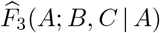. We show the formal results next (Proposition 4), and prove the result in the *Appendix*.

### Proposition 4.

Consider *J* polymorphic loci in populations *A*, *B*, and *C* with respective parametric reference allele frequencies *a*_*j*_, *b*_*j*_, *c*_*j*_ ∈ (0, 1), and suppose we take a random sample of *N* (*P*_*j*_) individuals at locus *j* in population *P* ∈ {*A, B, C*}, some of which may be related or inbred. The ratio estimator 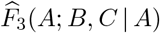 is approximately upwardly biased, assuming that its mean is well-approximated by the ratio of means of 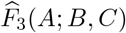 and 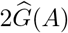 that it uses in its definition, with its upward approximate bias

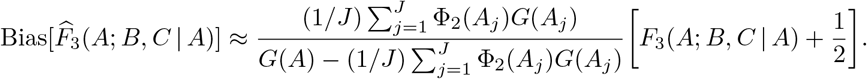

Moreover, an approximately unbiased estimator of *F*_3_(*A*; *B, C* | *A*) is

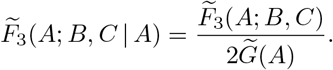

There is an upward approximate bias of the original normalized *F*_3_ estimator, and the bias is, as with the standard estimator of *F*_3_, due partially to the diversity and sampling in the target population. The new approximately unbiased estimator 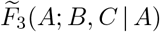 is based simply on the ratio of unbiased estimators of its components 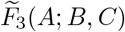 and 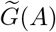.

Finally, we move to the four population statistics *F*_4_ and *D*. Note that the *F*_4_ statistic by definition only samples a single allele per population, and therefore the original estimator 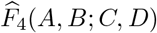 is intuitively unbiased. We show the formal results next (Proposition 5), and prove the result in the *Appendix*.

### Proposition 5.

Consider *J* polymorphic loci in populations *A*, *B*, *C*, and *D* with respective parametric reference allele frequencies *a_j_, b_j_, c_j_, d_j_* ∈ (0, 1), and suppose we take a random sample of *N* (*P*_*j*_) individuals at locus *j* in population *P* ∈ {*A, B, C, D*}, some of which may be related or inbred. The estimator 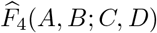 is unbiased.

Though the original *F*_4_ estimator is unbiased, the normalized *F*_4_ and *D* statistics are more complicated as they are ratio estimators, meaning their biases cannot be directly assessed. However, intuitively, because both estimators have 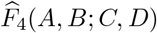 as their numerator, bias would seemingly derive from their denominator component. Next, we show formally in Proposition 6 that the normalized 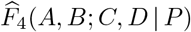 estimator is approximately upwardly biased, and prove the result in the *Appendix*.

### Proposition 6.

Consider *J* polymorphic loci in populations *A*, *B*, *C* and *D* with respective parametric reference allele frequencies *a_j_, b_j_, c_j_, d_j_* ∈ (0, 1), and suppose we take a random sample of *N* (*P*_*j*_) individuals at locus *j* in population *P* ∈ {*A, B, C, D*}, some of which may be related or inbred. The ratio estimator 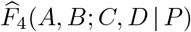 approximately upwardly biased, assuming that its mean is well-approximated by the ratio of means of 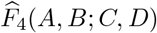 and 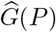 for any population *P* ∈ {*A, B, C, D*} that it uses in its definition, with its upward approximate bias

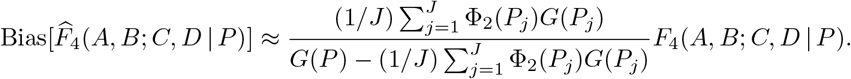

Moreover, an approximately unbiased estimator of *F*_4_(*A, B*; *C, D* | *P*) is

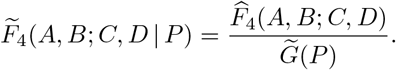

The reasoning that the original 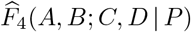 estimator has upward approximate bias is that its estimator 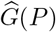 used in its denominator is downwardly biased. By using the unbiased estimator 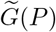 in its place within the denominator, we find a new estimator 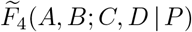 is approximately unbiased.

The bias property of the *D* statistic is different than the normalized *F*_4_ statistic, as the estimator 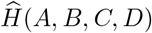 of its denominator is unbiased (Lemma 8 of the *Appendix*). Intuitively, this result is due to the denominator not having a product of frequencies for two alleles sampled from the same population. Because both its numerator and denominator are unbiased, we next show that the ratio estimator 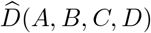 is approximately unbiased in Proposition 7, and prove the result in the *Appendix*.

### Proposition 7.

Consider *J* polymorphic loci in populations *A*, *B*, *C*, and *D* with respective parametric reference allele frequencies *a_j_, b_j_, c_j_, d_j_* ∈ (0, 1), and suppose we take a random sample of *N* (*P*_*j*_) individuals at locus *j* in population *P* ∈ {*A, B, C, D*}, some of which may be related or inbred. The ratio estimator 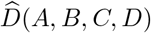 is approximately unbiased, assuming that its mean is well-approximated by the ratio of means of – 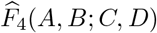 and 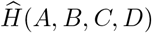 that it uses in its definition.

In addition to bias, variance is an important property of an estimator, as both bias and variance are components of mean squared error. Because the formulas and derivations for the variances of the *F*- and *D*-statistics are not particularly insightful, we relegate these results to the *Appendix*. Specifically, we provide the variances for 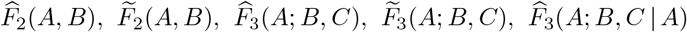,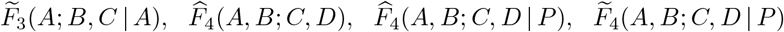, and 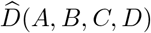 in Propositions 11, 12, 13, 15, 17, 19, 16, 21, 23, and 26 of the Appendix, respectively.

## Results

In the *Theory* and *Appendix*, we introduced new unbiased estimators of *F*_2_ and *F*_3_ statistics, and derived biases and variances (and hence mean squared errors) for the original and new estimators of *F*- and *D*-statistics. In this section, we theoretically evaluate the relative performances of the old biased estimators and the new unbiased estimators under an array of settings, including different mixtures of relatedness, inbreeding, sample sizes, and population parameters.

For all of our results we require the kinship coefficients for each pair of individuals. To acquire these values, we need to know if each individual is related to any other in the population and also whether they are inbred, and if so, how these values are quantified through the use of kinship coefficients (Φ_*xy*_). To summarize how an entire sample from a population *P* is related to each other at a locus, we use

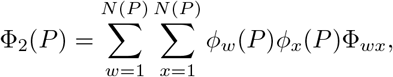

where *ϕ_w_*(*P*) and *ϕ_x_*(*P*) are weights of individuals *w* and *x* in population *P*, and in this study we use weights corresponding to the proportion of alleles contributed by individual *x* to the sample from population *P*, which is computed as

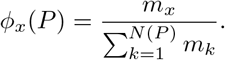

Here *m*_*x*_ is the ploidy of individual *x*. Moreover, using this weighting scheme, we also estimate the frequency of the reference allele at a biallelic locus as the sample proportion (McPeek et al., 2004; DeGiorgio et al., 2010; Harris and DeGiorgio, 2017a)

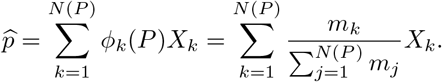

### Effect of population *F*-statistic value on mean squared error

The relationship between the population parameter for a statistic and the estimate based on a sample from the population is important to evaluate. We compare the difference in the mean squared error (MSE) between the biased 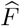 estimators and our unbiased 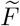 estimators to the true value of each statistic in the cases for which both estimators exist. The *F*_2_, *F*_3_, and *F*_4_ statistics require allele frequency information from either two, three, or four populations, respectively.

For our *F*_2_ comparisons, we use the sample allele frequencies from the YRI (sub-Saharan African) and CEU (central Europeans) from the 1000 Genomes Project (The 1000 Genomes Project Consortium, 2015) as the true population allele frequencies to obtain the true *F*_2_(*A, B*) statistic by using the population definition from the *Introduction*, with populations *A* = CEU and *B* = YRI. To evaluate the relative performances of *F*_2_ estimators over a range of true *F*_2_ values, we randomly sample 20 independent loci from both populations for 1000 independent replicates of *J* = 20 loci, yielding 1000 independent draws of the true *F*_2_ statistic, which ranged across the set of values *F*_2_ ∈ [0.02, 0.12]. Using these allele frequencies, along with the sample size and relatedness information, we also calculate the difference in MSE between the 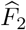 and 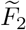 estimators by using Propositions 2, 11 and 12. We then calculate the MSE by summing the variance and squared bias. We note that the MSEs of the unbiased estimators are equal to their variances. We repeat this process for *F*_3_(*A*; *B, C*), *F*_3_(*A*; *B, C* | *A*), and *F*_4_(*A, B*; *C, D* | *A*) as these are the estimators that are biased in their 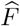 forms.

We use Propositions 3, 13, and 15 to determine the MSE for both biased and unbiased *F*_3_ estimators by including allele frequency information from the JPT (Japanese) population where *A* = JPT, *B* = CEU, and *C* = YRI, with true range for *F*_3_ ∈ [0.00, 0.08]. For the normalized *F*_3_(*A*; *B, C* | *A*) estimators, we compare MSE between the biased and unbiased versions by using bias and variances derived in Propositions 4, 17, and 19, with true range for normalized *F*_3_ ∈ [0.00, 0.30]. Finally, we estimate the MSE for the normalized *F*_4_(*A, B*; *C, D* | *A*) estimators by including GIH (Gujurati Indian) allele frequency data and using the derivations in Propositions 6, 21, and 23. In this case we set *A* = YRI, *B* = CEU, *C* = JPT, and *D* = GIH for a true range of normalized *F*_4_ ∈ [−0.3, 0.2].

For each analysis we estimate MSE for instances when samples of 60 diploid individuals from each population include 30 relative pairs, including ten avuncular relationships, ten inbred full siblings, and ten outbred full siblings. We also assumed every individual was related to exactly one other individual. In these estimates, all populations contain the same composition of related individuals.

The difference in log_10_(MSE) between 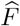 and 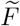 estimators for *F*_2_(*A, B*), *F*_3_(*A*; *B, C*), *F*_3_(*A*; *B, C* | *A*), and *F*_4_(*A, B*; *C, D* | *A*) show similar trends with respect to the true *F*-statistic values. Specifically, the difference in log_10_(MSE) decreases as the true *F*-statistic value approaches zero (Figure 2). In our evaluation of *F*_4_(*A, B*; *C, D* | *A*), we considered both positive and negative values for its true value, which shows that the difference in log_10_(MSE) of 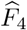 and 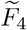 exhibits a quadratic shaped trend as a function of true *F*_4_. Overall, we notice that the difference in MSE between biased and unbiased estimators is dependent on the true value of the *F*-statistic, with the least difference occurring when the true *F*-statistic is closest to zero.

**Figure 2:**
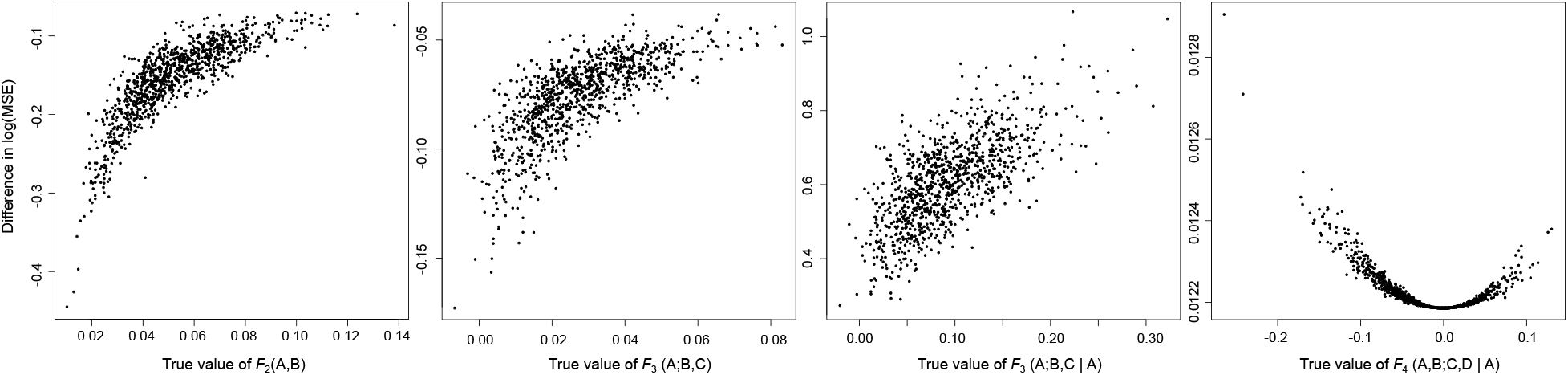
Difference in theoretically calculated log_10_(MSE) of 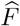 and 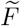 estimators when including relatives or inbred individuals. The MSE is estimated for instances when samples of 60 individuals include individuals related to exactly one other in the sample, with 10 pairs of avuncular relationships, 10 pairs of inbred full siblings and 10 pairs of outbred full siblings. Each point represents calculations from *J* = 20 randomly sampled loci from the 1000 Genomes Project dataset for CEU, European, YRI African, JPT Japanese, and GIH Indian populations. For *F*_2_(*A, B*) we use *A* = CEU and *B* = YRI, while for *F*_3_(*A*; *B, C*) and *F*_3_(*A*; *B, C* | *A*) we use *A* = JPT, *B* = CEU, and *C* = YRI and for *F*_4_(*A, B*; *C, D* | *A*) we assign *A* = YRI, *B* = CEU, *C* = JPT, and *D* = GIH.

### Effect of sample size on mean squared error

To probe how sample size within each population affects the difference in estimator error rate, we theoretically computed the MSE for both 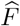 and 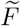 estimators when different numbers of individuals are sampled, with the constraint that every sampled individual is related to exactly one other individual in the sample from that population. Specifically, we evaluate the impact on these estimators when sampling from one pair to 50 pairs of related individuals, with relationships of inbred full sibling pairs (Φ_*xy*_ = 3/8), outbred full sibling pairs (Φ_*xy*_ = 1/4), parent-offspring pairs (Φ_*xy*_ = 1/4), and avuncular pairs (Φ_*xy*_ = 1/8). We compute the MSE as in *Effect of population F -statistic value on mean squared error* section above.

In almost all cases, the biased 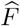 estimators always displayed elevated MSE compared to their corresponding unbiased 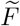 estimators (Figures 4 and S1-S3). For both estimators, we see a clear decrease in the MSE as the number of sampled individuals increases, with the greatest error observed when two individuals are sampled. As expected, a greater sample size allows one to better estimate allele frequencies, and ultimately reduces the mean pairwise kinship coefficient within the sample, as the number of pairs in the sample grows quadratically but the number of relative pairs grows linearly. We also find that the difference in the MSE at larger sample sizes is not as pronounced for normalized *F*_4_ as it is for *F*_2_, *F*_3_, and normalized *F*_3_, as the difference in bias between the biased and unbiased estimators is much smaller for normalized *F*_4_ (Figure S3).

### Effect of sample composition on mean squared error

Different types of relatives have different proportions of their alleles shared identical by descent, and thus have different pairwise kinship coefficients. Because we have demonstrated that bias and variance (and hence MSE) of estimators are influenced by within-population mean pairwise kinship coefficient across sampled individuals, the distribution of relative types within a sample will impact overall *F*-statistic estimation error. For this reason, it is important to examine how our *F*-statistics are affected by samples containing diverse mixtures of relative types. Specifically, to accurately assess the impact of relative composition, we hold sample sizes, number of relative pairs, and true population *F*-statistic values constant.

We computed the theoretical MSE when samples of 50 pairs of relatives (100 diploid individuals sampled) contain relative pairs of three different types as in Harris and DeGiorgio (2017a). In addition, each individual is related to exactly one other individual in the sample from the same population. For each statistic we vary the number of pairs related by each of three types of relationships between zero and 50, with 1326 combinations for each. We repeat this process for three configurations of relationships to probe estimator error as a function of the mixture of relative types. We also provide comparisons among inclusion of male-male full siblings (Φ_*xy*_ = 1/2), male-female full siblings (Φ_*xy*_ = 3/8) and female-female full siblings (Φ_*xy*_ = 1/4) at mixed-ploidy loci such as on the X chromosome (DeGiorgio et al., 2010), with results showing elevated MSE for both estimators for higher male-male sibling proportions, when compared to male-female or female-female full siblings. To investigate the effects of inbred individuals, we also provide a comparison between inbred full siblings (Φ_*xy*_ = 3/8) with inbreeding coefficient *f*_*x*_ = *f_y_* = 1/4, and outbred full-siblings (Φ_*xy*_ = 1/4) at a autosomal diploid loci. We see that MSE is higher for inbred full-siblings than for outbred full-siblings in all cases examined (Figures 3 and S4-S6).

**Figure 3:**
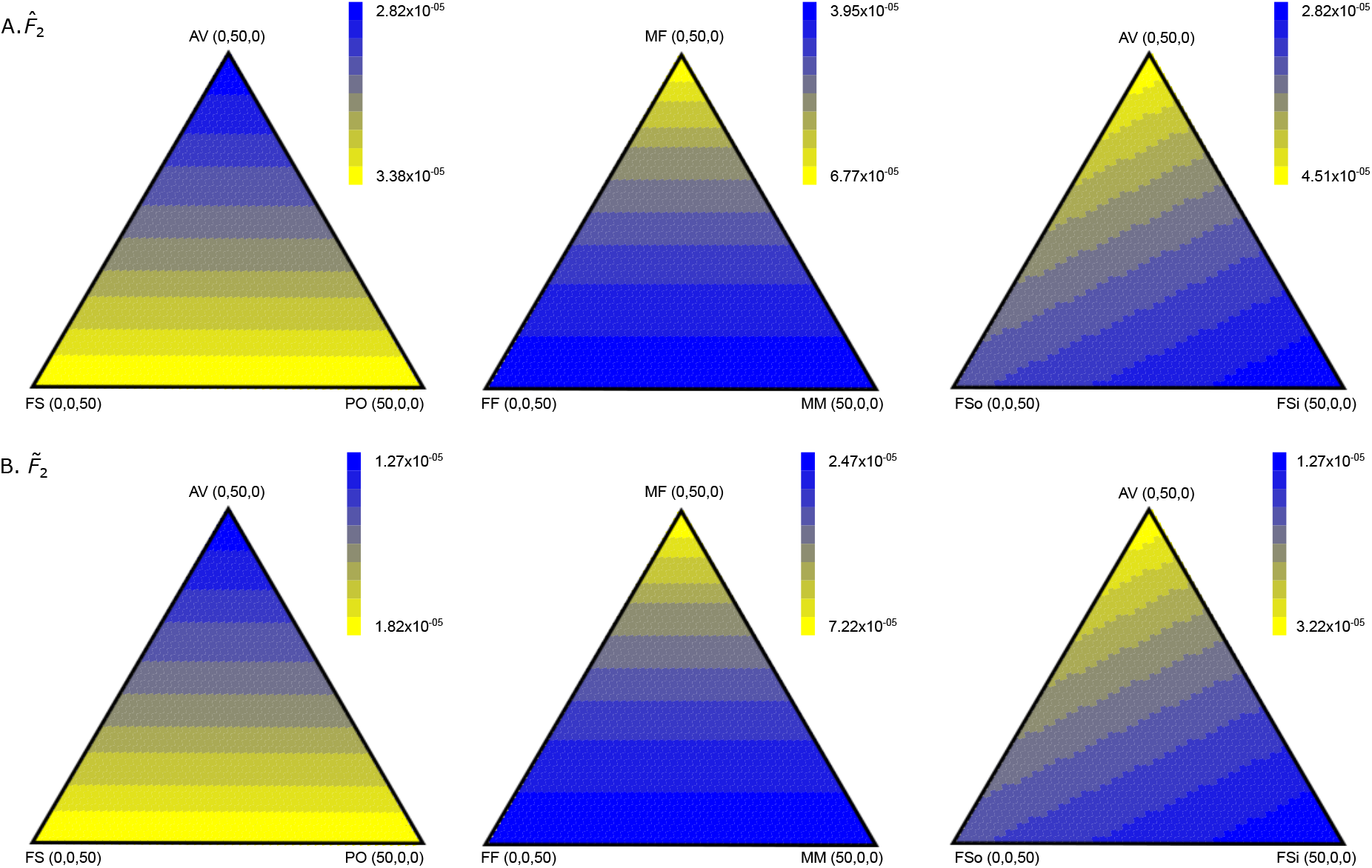
Theoretically calculated MSE of 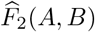 and 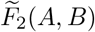 when including relatives or inbred individuals for *J* = 20 loci. The MSE is estimated for instances when samples of 100 individuals include individuals related to exactly one other in the sample. The first column shows MSE for samples with different combinations of parent-offsring (PO), full sibling (FS), and avuncular (AV) relationships, the second includes full siblings that are male-male (MM), male-female (MF) and female-female (MF). The last column includes AV relationships as well as inbred (FSi) and outbred (FSo) full siblings. The true value of *F*_2_(*A, B*) is 0.071.

**Figure 4:**
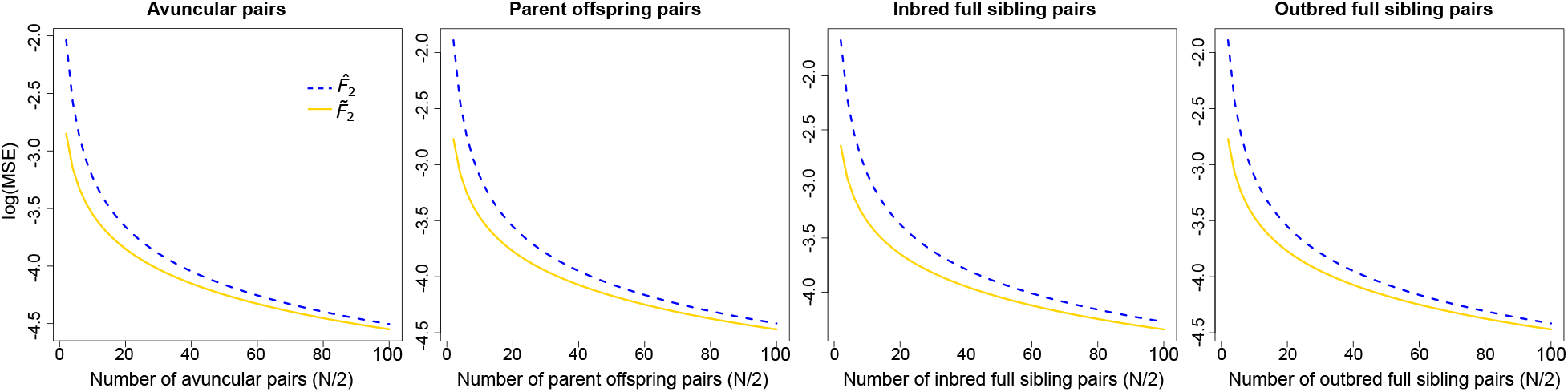
Mean squared error theoretically calculated for 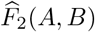 and 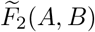 across different sample sizes or related pairs of individuals, including avuncular relationships, parent-offspring relationships, inbred full siblings, and outbred full siblings. The number of sampled individuals ranges from 2 to 100 with the number of relative pairs equaling half the total sampled, all computed using *J* = 20 loci. The true value of *F*_2_(*A, B*) is 0.071.

We also note that the value of MSE for the biased 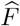 estimators is always greater than the value for their corresponding unbiased 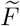 statistics, which is true in part due to the values of the true *F*-statistics for the loci we chose to use. Though the MSE is higher for the biased estimators, the variation in MSE values is similar for both estimators. For example, the data point with the highest proportion of avuncular relatives has the lowest MSE when compared to parent-offspring relationships and outbred full siblings. In all tested settings (Figures 3 and S4-S6), we notice similar patterns of MSE variation when comparing 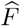 estimators with 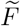 estimators. This pattern is again shared when comparing MSE variation among the estimators for *F*_2_, *F*_3_, normalized *F*_3_, and normalized *F*_4_. We can conclude that in all of these cases, the value of the mean kinship coefficient is most important in determining MSE when sample size and true *F*-statistic value are fixed.

### Simulations to evaluate theoretical MSE approximations

To verify that our theoretical approximations for MSE are reasonable, we simulate samples containing related individuals and use them to compute the biased 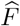- and unbiased 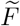-statistics as well as calculate their biases, variances, and MSEs. For each population (CEU, YRI, GIH, and JPT), we simulate 10 non-inbred parent-offspring pairs with each individual related to exactly one other individual in the sample. Genotypes for each individual are simulated by first sampling two alleles with replacement according to their respective population allele frequencies from each of the populations (CEU, YRI, GIH, and JPT) to create a set of 20 unrelated individuals per population. Individuals *x* and *y* that form one of the 10 relative pairs have the genotype of individual *y* modified according to their relationship type. Specifically, for each relative type, there are probabilities Δ_0_, Δ_1_, and Δ_2_ that the two individuals will share zero, one, or two alleles identical by descent, respectively. The first allele of individual *y* is copied from the first allele from individual *x* with probability Δ_1_ and the entire genotype of individual *x* is copied over to individual *y* with probability Δ_2_. This process is repeated across 20 independent loci to generate a sample of 20 individuals with 10 relative pairs in each population with genotypes taken at *J* = 20 independent loci. To generate 20 independent loci from the four 1000 Genomes Project populations, we used loci either on separate chromosomes, or at least one megabase away from each other.

For each of our new unbiased estimators we compute the bias, variance, and MSE along with the same values for the original estimators (Figures S7-S10). Comparing the bias measurements in these figures, we observe a clear reduction in bias when applying the 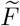 estimators as opposed to the 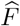 estimators. However, the variance is highly similar for 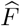 and 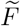 in all cases. As the value of variance is much larger than the magnitude of the bias (by an order of magnitude) and hence the squared bias, the resulting MSE is consequently similar as well. Because *F*_4_ is quantifying the relationship among four populations, more simulations may be required to converge to the pattern seen by theoretical simulations. For this reason, we increased the number of simulations used to compute the bias, variance, and MSE to 10^4^ for each data point in Figure S10, whereas 10^3^ simulation replicates were used for *F*_2_, and both versions of *F*_3_.

To compare the accuracy of our theoretical approximations to simulation results across a spectrum of relatedness between individuals in a sample, we simulate combinations of parent offspring, outbred full sibling, and avuncular relationships. In a manner similar to described above (first paragraph of *Simulations to evaluate theoretical MSE approximations*, we simulate a total of 10 relative pairs made up of a combination of each of the three relative types, with the number of each relative type ranging from zero to 10. We simulate each of these 66 distinct settings of relative type combinations with genotypes sampled at *J* = 20 independent loci, and completed 1000 independent replicates of each setting to obtain accurate measurements of bias, variance, and MSE for each simulation setting, with each simulation using true *F*-statistic values specified in Figures S7 and S8-S10. We compute the bias, variance, and MSE for simulations, and compare these values to theoretically calculated computations for each relative combination (Figures S12-S15). We find that although noisier, the bias variance, and MSE patterns in our simulation results match theoretical calculations, suggesting that our theoretical computations are accurate. For all cases the simulated bias measurements for the 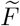 estimators are close to zero, whereas the 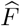 estimators display bias measurements matching the theoretically calculated 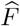 bias values.

### Utility and applications of unbiased estimators

In previous sections, we have shown through simulations that our theoretical results are producing expected patterns and evaluated the performance of our unbiased estimators under varying combinations of relatives, true *F*-statistic values, and sample sizes. In this section we show some potential applications of these estimators, using both simulated and empirical data. As discussed previously in the *Introduction*, the value of *F*_3_(*A*; *B, C*) can be used to identify whether population *A* is the result of admixture between populations related to *B* and *C* (Figure 1). A negative value of *F*_3_ indicates the presence of this process, whereas a non-negative value is inconclusive and means that further tests may be required to verify a history of admixture. However, because 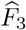 is upwardly biased and because 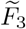 corrects for this bias, 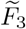 might allow us to detect admixture in cases where 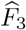 would be inconclusive, even without the presence of related or inbred individuals.

To explore this hypothesis, we first examine an admixture scenario in which *F*_3_(*A*; *B, C*) might provide marginally negative values. We simulate two populations (*B* and *C*) with effective population size of 10^4^ diploid individuals (Takahata, 1993) that diverged 2000 generations prior to sampling using SLiM (Haller and Messer, 2019). This simple divergence model has parameters inspired by the history relating African and non-African human populations (Gravel et al., 2011). These populations then merge with admixture proportions 0.4 and 0.6 for *B* and *C*, respectively, to form population *A* 400 generations prior to sampling. Using these parameters, the expected value is *F*_3_(*A*; *B, C*) = −0.0568. To generate genetic data from this model, we evolved sequences with a per-site per-generation mutation rate of *μ* = 1.25 × 10^−8^ (Scally and Durbin, 2012) and a uniform per-site per-generation recombination rate of *r* = 10^−8^ (Payseur and Nachman, 2000). We output 20 two megabase chromosomal regions containing allele frequency information for all three populations. Using these simulated population allele frequencies for each of these three populations, we then simulate 50 instances of 50 unrelated individuals each. We then compute 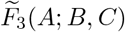 and 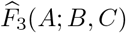 across *J* = 20 loci, either on separate chromosomes or at least one megabase away from each other to ensure independence.

Figure 5 illustrates that 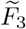 values are lower than 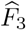, and are always negative when 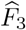 values are almost always positive. Because this statistic is used to test for admixture and a negative result indicates the presence of admixture, the use of the biased estimator leads to a different conclusion than when using the unbiased estimator. We also explore a setting in which populations contain related individuals with the same parameters as described above. Using allele frequency information from the three populations simulated previously (*A, B,* and *C*) we generate 50 individuals for each population, in which there are 25 parent-offspring pairs. Similarly to when relatives are not included in the population, 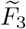 values are lower than 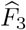, with most 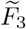 values giving negative values, and all 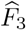 providing positive values, again lending different conclusions about the underlying demographic history of these populations (Figure 5).

**Figure 5:**
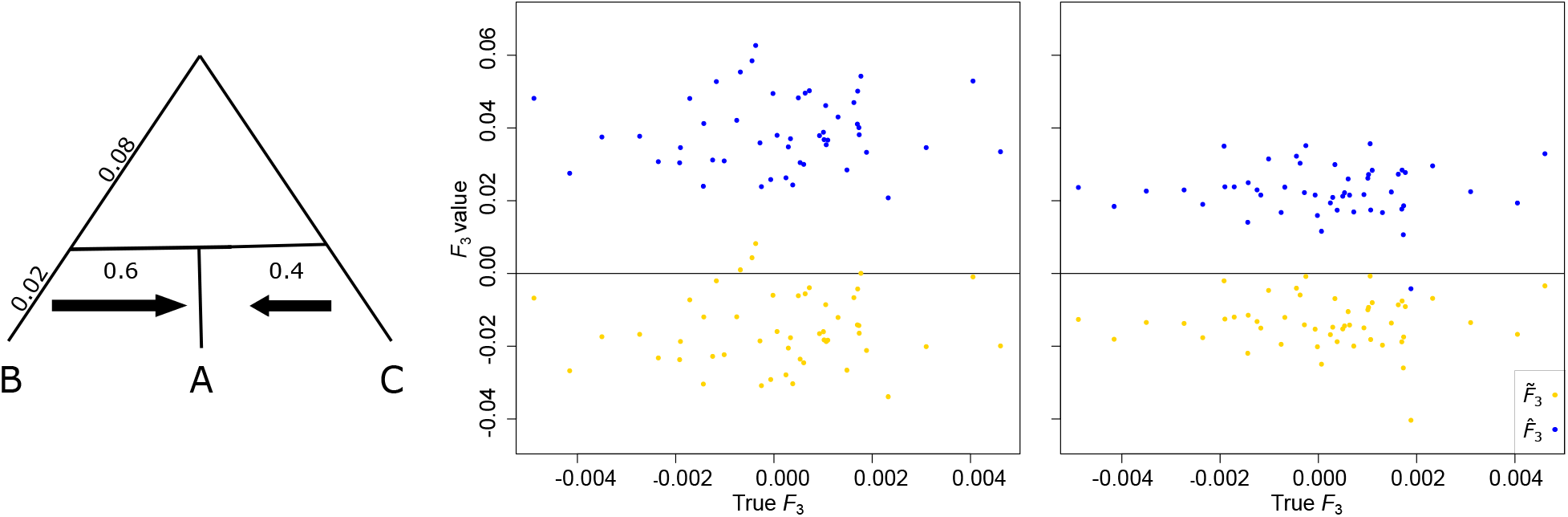
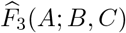 and 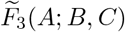 calculated for simulations where populations *B* and *C* merge with admixture proportions of 0.4 and 0.6, respectively, 400 generations ago (0.02 coalescent units) to form population *A*. Comparison of simulations with (middle) and without (right) relatives. The panel in the middle shows results for a sample containing 25 parent-offspring pairs, while the panel on the right shows results for 50 unrelated individuals.

Finally, we test the performance of our statistics on empirical data. We use populations from the HGDP SNP dataset (Li et al., 2008) that include related individuals (Rosenberg, 2006). Specifically, we use genotype information from Colombian, Lahu, Melanesian, Mandenka, San, and Druze populations, and we sample 20 independent loci that are at least one megabase apart from all populations for 1000 independent replicates of *J* = 20 loci, yielding 1000 independent draws. Each of these populations contains between two and 14 pairs of inferred related individuals, according to Rosenberg (2006). Using distinct pairs for *F*_2_, triples for *F*_3_, and quadruples for *F*_4_ of these populations and the relationships from Rosenberg (2006), we estimate 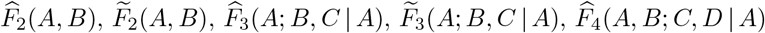, and 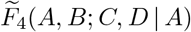 and compare the mean and standard deviation of the biased and unbiased estimators (Figure 6). In all cases shown, the biased estimator has higher mean than the unbiased estimator, although the standard deviations are similar for both. This indicates that correcting the bias generated by related individuals yields more accurate *F*-statistic estimates with minimal cost in precision of the estimates.

**Figure 6:**
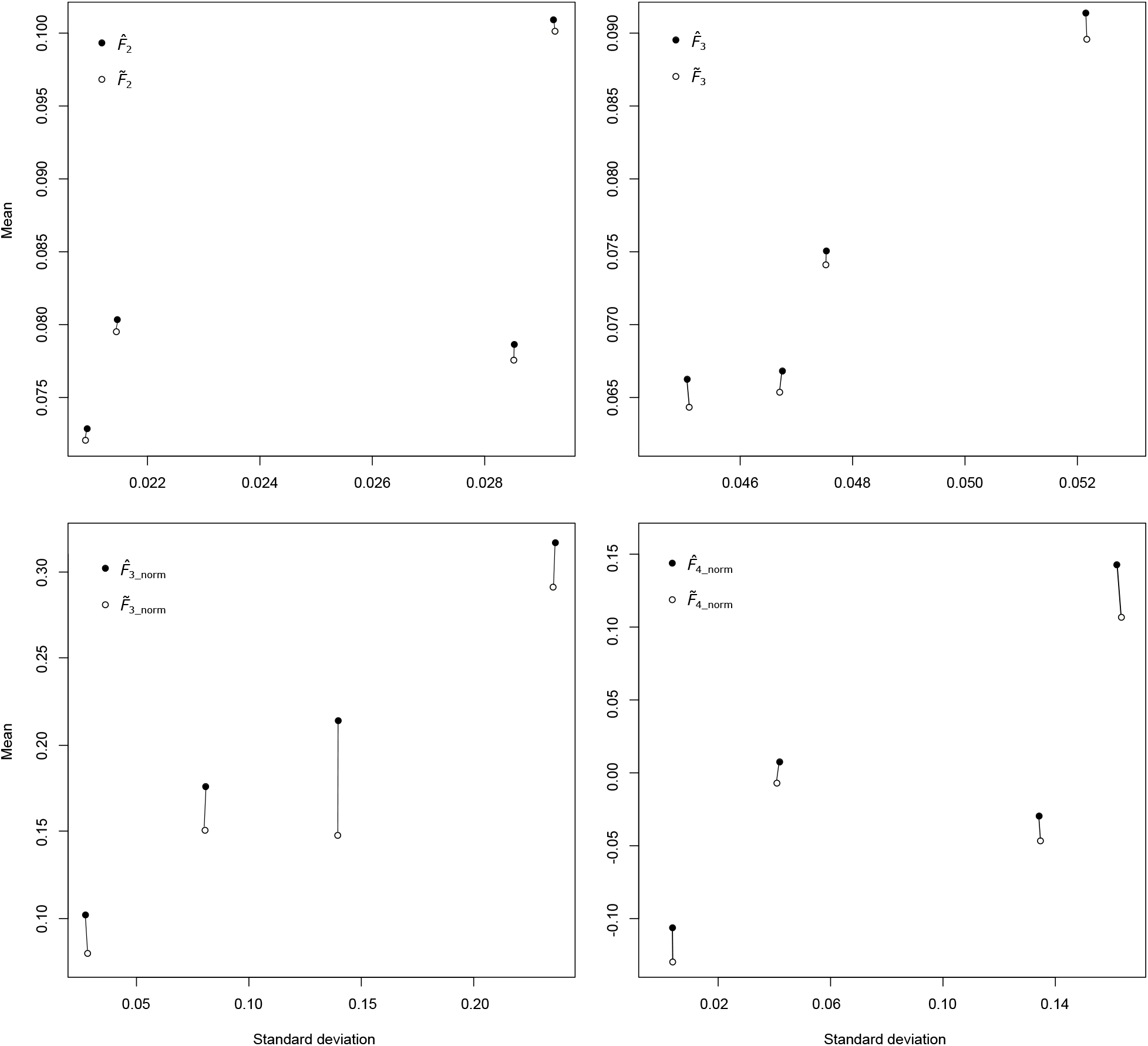
The difference between the means and standard deviations of biased and unbiased estimators of *F*_2_, normalized and un-normalized *F*_3_ and normalized *F*_4_ when estimated with genotype information from four different combinations of Colombian, Lahu, Melanesian, Mandenka, San, and Druze populations. All of these populations include between two and 14 relative pairs. The black dots represent the values for the biased estimators, while the white dots show the value for the unbiased estimators. Each mean and standard deviation was calculated for a combination of two, three, or four populations, for *F*_2_, *F*_3_, and *F*_4_, respectively, and consists of 1000 estimates of the statistic, each calculated from *J* = 20 randomly samples single nucleotide polymorphisms from the genome. The top row has values for the *F*_2_(*A, B*) (left) and *F*_3_(*A*; *B, C*) (right), while the bottom row shows results for normalized *F*_3_(*A*; *B, C* | *A*) (left) and normalized *F*_4_(*A, B*; *C, D* | *A*) (right).

## Discussion

We have introduced the unbiased estimators 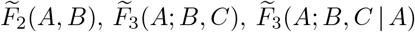, and 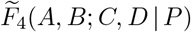 as well as shown that the estimators 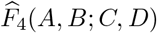 and 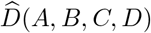 are unbiased with the inclusion of related and inbred individuals. In addition, we have demonstrated that the variance of 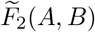 is similar to that of 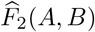, as are the variances of 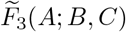 and 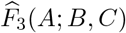. We have also provided variance calculations for all other *F*- and *D*-statistic estimators included in this study. Using these calculations we have compared the performance of the biased and newly derived unbiased estimators, and shown that in most cases the unbiased estimators have lower MSE values than the biased estimators of the same statistic.

Interestingly, the two statistics that sample from each analyzed population only once per locus—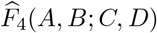 and 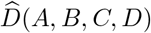—are unbiased with the inclusion of related or inbred individuals, whereas 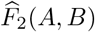 which samples from each population *A* and *B* twice, and 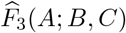, which samples from population *A* twice, are biased. This process of sampling more than once from a single population per locus is responsible for creating bias due simply to finite sample size, which is exacerbated by the inclusion of related or inbred individuals within the twice-sampled population.

The development of these unbiased statistics, and the proofs showing other statistics are unbiased is beneficial for anthropologists interested in populations such as hunter-gatherers, some of which are often small and widely dispersed yet retain high genetic diversity (Kim et al., 2014). Small population sizes may necessitate the sampling of close relatives, such as parents and offspring, or siblings. Along with small human populations, these statistics are often applied to non-human species. Some, such as elephants, rhinoceros, and cheetahs are close to extinction or have extremely small and inbred populations due to human activity. The *F*- and *D*-statistics may prove important in conservation efforts to test how (and whether) different populations of these animals are interacting. For these reasons, having estimators that are unbiased under such conditions is imperative in making accurate inferences about the relationships of such small populations with others. Although it may not be possible to identify relatives through the sampling process, especially in the case of wild animals, there are methods available to identify related individuals and estimate their likely degree of relatedness once the samples have been sequenced (Epstein et al., 2000). The inferences from these methods will allow users to identify pairwise kinship coefficients necessary to apply the unbiased statistics of this study. Moreover, even if relatedness is difficult to assess, many of the original statistics (*i.e.*, 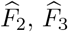, normalized 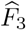, and normalized 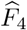) have bias due to finite sample size, and simply accounting for the bias may be important to accurately assess population history and diversity across populations (*e.g.*, Figure 5).

A key consideration when evaluating the importance unbiased estimators of *F*- and *D*-statistics is their potential use. Specifically, a number of applications of these statistics do not employ the raw estimates, but instead standardized estimates (Soraggi et al., 2018; Zheng and Janke, 2018), where a particular *F*- or *D*-statistic has its genomewide mean subtracted, and is normalized by the standard error using a genomic block jackknife procedure (Reich et al., 2009). Indeed, subtracting out this genomewide mean may circumvent bias issues. However, this assumes that all genomic blocks have similar sample properties, yet blocks with reduced sample size (*e.g.*, in regions with difficult to call genotypes) may still deviate from the genomewide expectation. In contrast, accounting for this bias due to finite sample size would provide estimates closer to the genomewide mean. Because the variance for these biased and unbiased estimators is approximately the same (compare Propositions 11 vs. 12 and 13 vs. 15), the standard errors used for normalizing these statistics are expected to be comparable, and thus, the unbiased estimators of the *F*-statistics derived here represent a more robust alternative to the original biased estimators, regardless of whether the raw or standardized values of the statistics are used. Furthermore, the raw value of some statistics, such as using the *F*_3_ statistic to detect population admixture, is important, and without correcting the bias of such statistics (Figure 5), key historical events relating populations could be missed.

The *F*- and *D*-statistics evaluated here are the most commonly used. However, since their development by Reich et al. (2009) and Patterson et al. (2012), other *D*-statistic type tests have been formulated to not only detect admixture, but also to identify the direction of gene flow—namely the partitioned *D*-statistics of Eaton and Ree (2013) and the *D*_FOIL_ statistics of Pease and Hahn (2015). Specifically, the *D*_FOIL_ statistics as originally formulated by Pease and Hahn (2015) sampled a single lineage (or allele) from each of a set of five populations *A*, *B*, *C*, *D*, and *O*, with a symmetric rooted topology ((*AB*)(*CD*)) relating populations *A*, *B*, *C*, and *D*, and with *O* an outgroup to these populations used to polarize the ancestral allelic state. Subsequently, Harris and DeGiorgio (2017b) derived allele frequency formulas for the *D*_FOIL_ statistics, and showed that allele frequency information for the outgroup population *O* is not needed for computation. The *D*_FOIL_ statistics are a set of four quantities (Harris and DeGiorgio, 2017b)

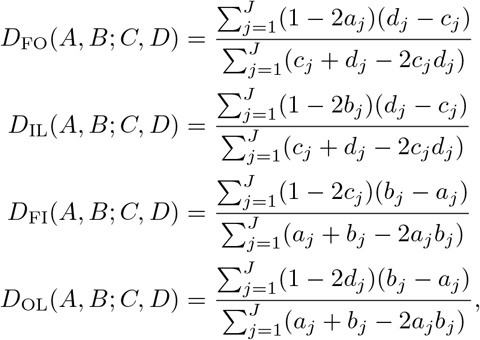

each of which does not have the frequencies for two alleles sampled from a single population multiplying each other. Hence, using sample allele frequencies in place of the population quantities would still yield approximately unbiased estimators of the *D*_FOIL_ statistics, regardless of whether related or inbred individuals were included in the sample. Though we chose to focus on the more classic *F*- and *D*-statistics, variance quantities for these partitioned *D* and *D*_FOIL_ statistics can be readily computed as we have done for other ratio estimators in this study.

Though we have only shown results when all populations contain samples with the same relative pair composition, it is trivial to include different relative types in different populations within these statistics. In addition, it is also possible to apply our new unbiased estimators when only some or none of the populations contain related or inbred individuals. Moreover, though we have demonstrated results for allele frequencies estimated as the sample proportion, we could have instead used the best linear unbiased estimator (BLUE) of McPeek et al. (2004), as all derivations in this article are based on a general form of a linear unbiased estimator. The BLUE allele frequency estimator would have superior properties to the sample proportion discussed here, as it has smallest variance (McPeek et al., 2004), and this reduction in variance translates to functions of the allele frequency as highlighted by improvements in both expected heterozygosity and *F*_ST_ by Harris and DeGiorgio (2017a). To apply the BLUE estimator, we would simply alter the weight *ϕ_x_*(*P*) of an individual *x* in population *P* at a particular locus with the equation

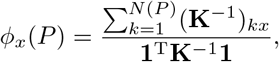

where **K** ∈ ℝ^*N*(*P*)×*N*(*P*)^ is the matrix of pairwise kinship coefficients, with element in row *j* and column *k* given by **K**_*jk*_ = Φ_*jk*_, **1** ∈ ℝ^*N*(*P*)^ is a column vector of ones, and superscript T indicates transpose. To facilitate easy application of these statistics, we have developed open-source software `funbiased` for use by the scientific community, which is available at https://github.com/MehreenRuhi/funbiased.

## Acknowledgments

We thank George (PJ) Perry for the helpful comments on an early draft of this manuscript and Alexandre Harris for fruitful discussions about our simulation protocol. This research was funded by National Institutes of Health grant R35GM128590, National Science Foundation grants DEB-1949268 and BCS-2001063, a NIGMS funded training grant on Computation, Bioinformatics, and Statistics (Predoctoral Training Program T32GM102057), and the NASA Pennsylvania Space Grant Graduate Fellowship. Computations for this research were performed on the Pennsylvania State University’s Institute for Computational and Data Sciences Advanced CyberInfrastructure (ICDS-ACI).

## Appendix

In this section, we provide proofs of key lemmas and propositions from the *Theory* section, and also develop and prove other important results.

### Proof of Lemma 1.

We first calculate

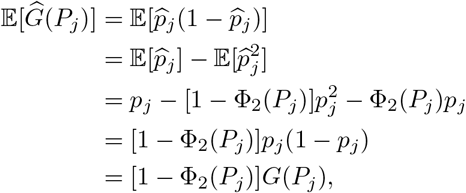

which gives

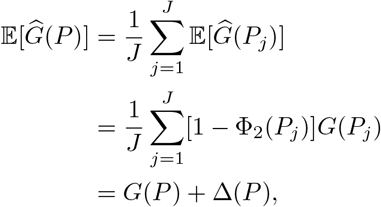

where we define the downward bias of 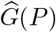 as

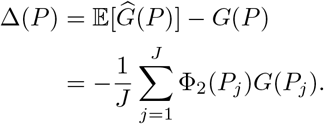

It follows that 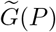 is an unbiased estimator of *G*(*P*) because

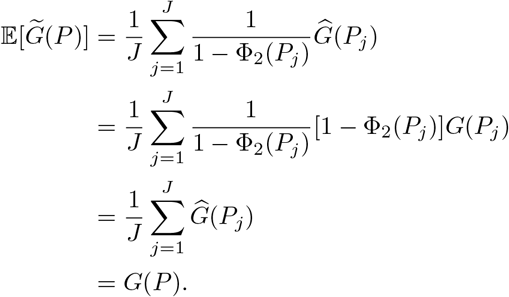

### Proof of Proposition 2.

We first calculate

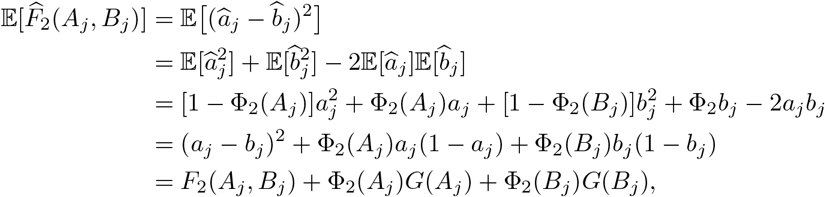

which gives

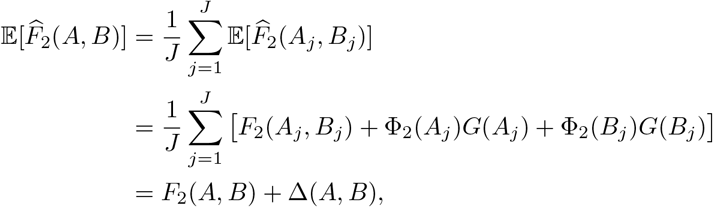

where we define the upward bias of 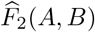 as

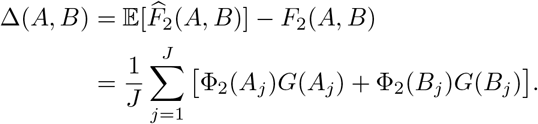

It follows that 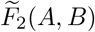 is an unbiased estimator of *F*_2_(*A, B*) because

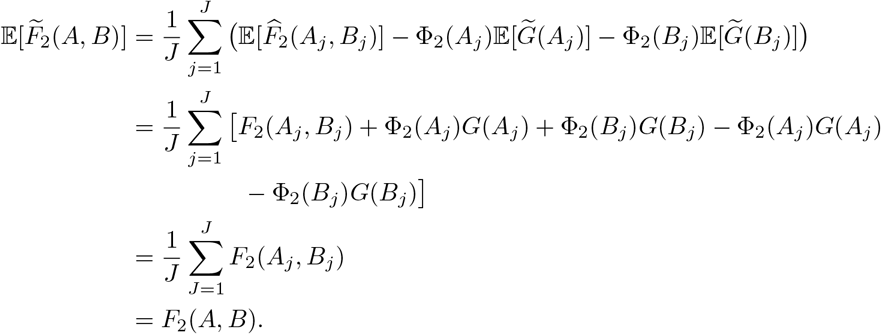

### Proof of Proposition 3.

We first calculate

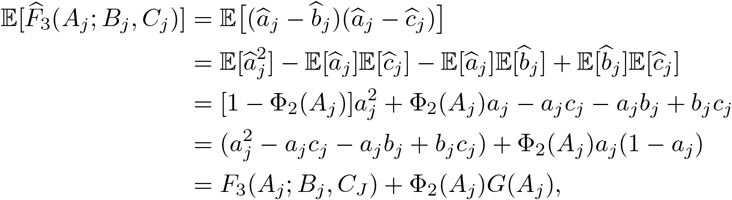

which gives

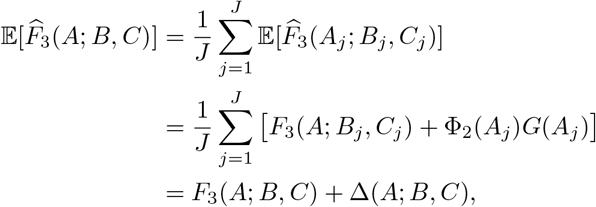

where we define the upward bias of 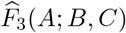 as

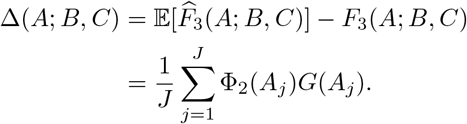

It follows that 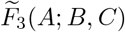 is an unbiased estimator of *F*_3_(*A*; *B, C*) because

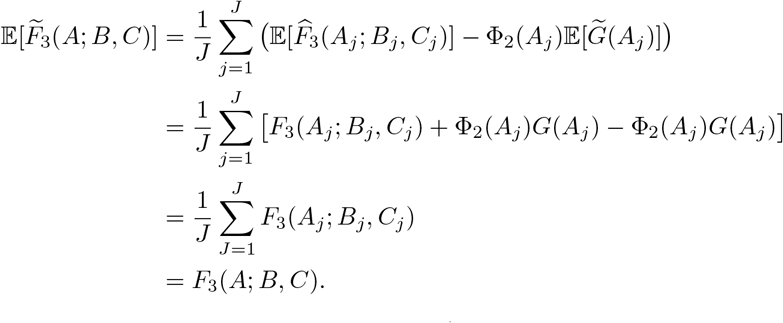

### Proof of Proposition 4.

Assuming that the expectation of 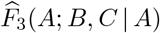 is approximately equal to the ratio of expectations of 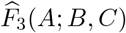 and 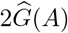, we find that

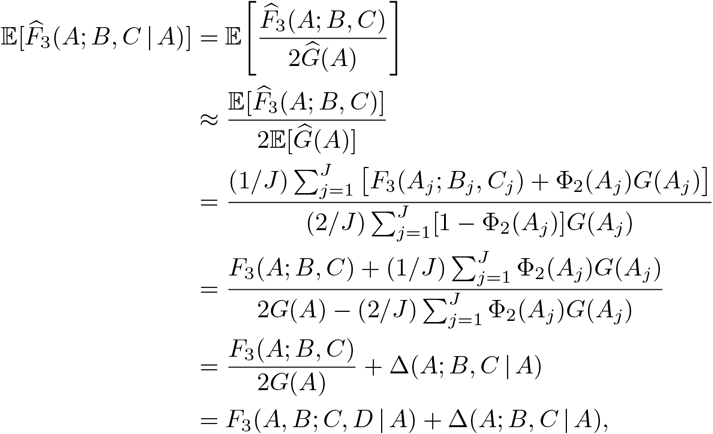

where we define the upward bias of 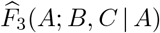 as

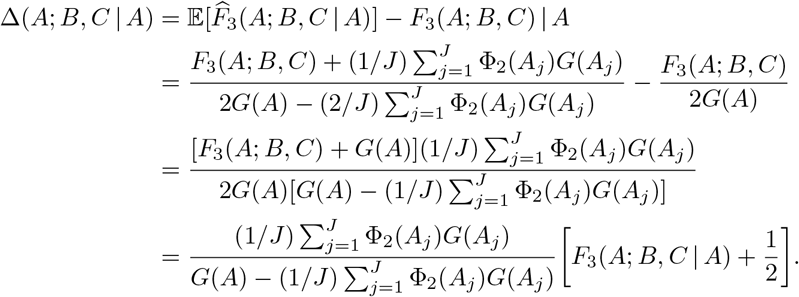

We can also see that 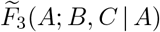 an approximately unbiased estimator of *F*_3_(*A*; *B, C* | *A*) because

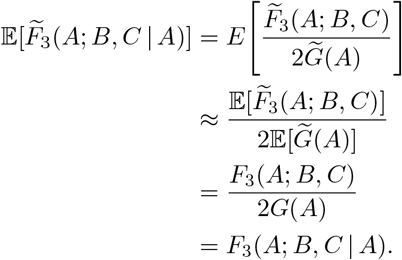

### Proof of Proposition 5.

We first calculate

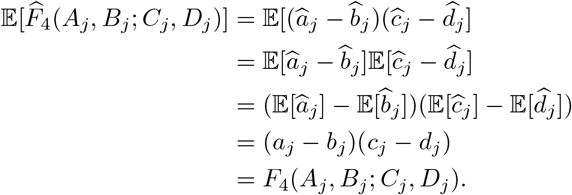

We show that 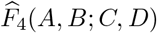 is unbiased estimator of *F*_4_(*A, B*; *C, D*) because

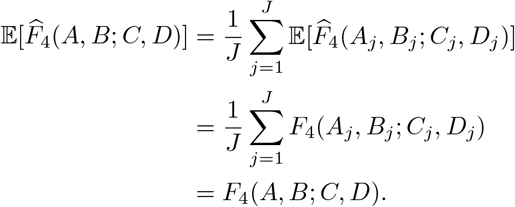

### Proof of Proposition 6.

Assuming that the expectation of 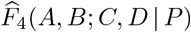 is approximately equal to the ratio of expectations of 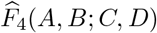 and 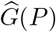 for some *P* ∈ {*A, B, C, D*}, we find that

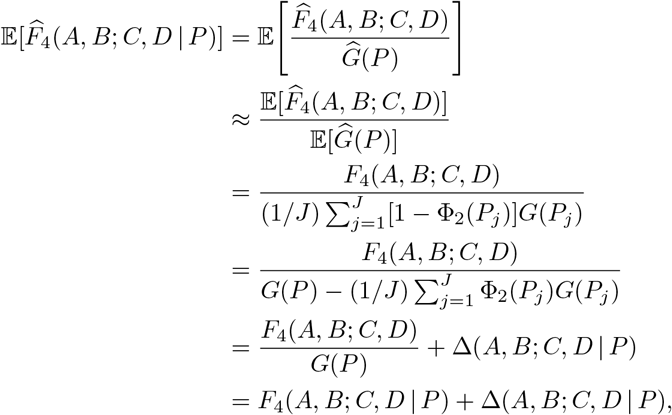

where we define the approximate upward bias of 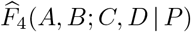 as

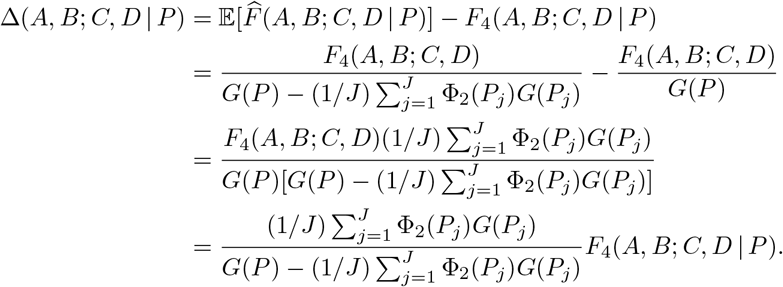

We can also see that 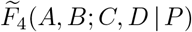 is an approximately unbiased estimator of *F*_4_(*A, B*; *C, D* | *P*) because

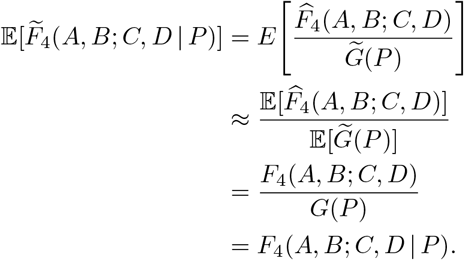

### Lemma 8.

Consider *J* polymorphic loci in populations *A*, *B*, *C*, and *D* with respective parametric reference allele frequencies *a*_*j*_, *b*_*j*_, *c*_*j*_, *d*_*j*_ ∈ (0, 1), and suppose we take a random sample of *N* (*P*_*j*_) individuals at locus *j* in population *P* ∈ {*A, B, C, D*}, some of which may be related or inbred. The estimator 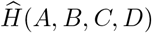 is unbiased.

*Proof.* We first calculate

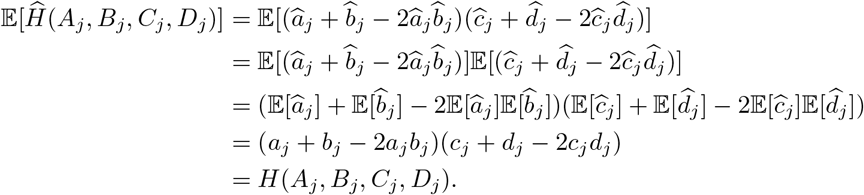

We show that 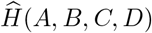 is unbiased estimator of *H*(*A, B, C, D*) because

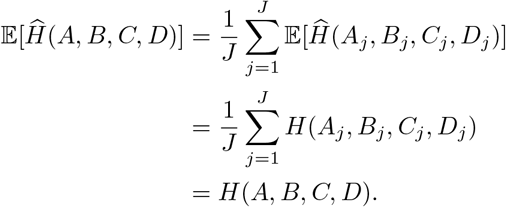

### Proof of Proposition 7.

Assuming that the expectation of 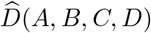 is approximately equal to the ratio of expectations of – 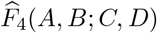 and 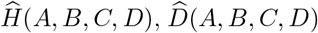 is an approximately unbiased estimator of *D*(*A, B, C, D*) because

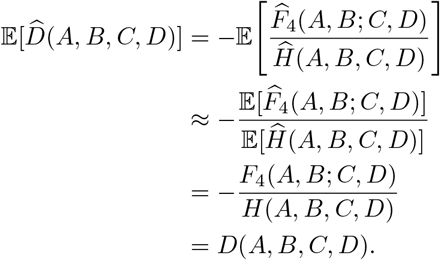

### Lemma 9.

Consider *J* independent polymorphic loci in a population *P* with parametric reference allele frequencies *p*_*j*_ ∈ (0, 1), and suppose we take a random sample of *N* (*P*_*j*_) individuals at locus *j*, some of which may be related or inbred. Moreover, assume that no individual is related to more than one other individual, which makes the terms Φ_3_(*P*_*j*_), Φ_4_(*P*_*j*_), Φ_2,2_(*P*_*j*_), and Φ_2_(*P*_*j*_)^2^ negligible to Φ_2_(*P*_*j*_). Based on this simplifying assumption, the estimator 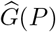 has an approximate variance

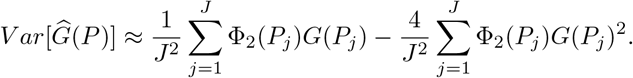

Moreover, the respective approximate variance for the unbiased estimator 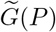 is

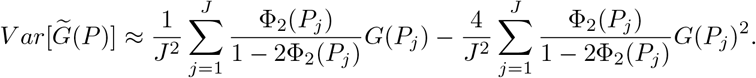

*Proof.* From the proo of Lemma 1, we have

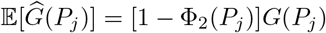

and we calculate

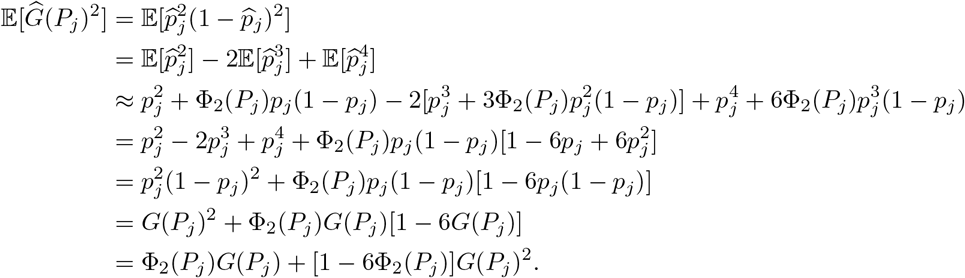

Therefore, we have that

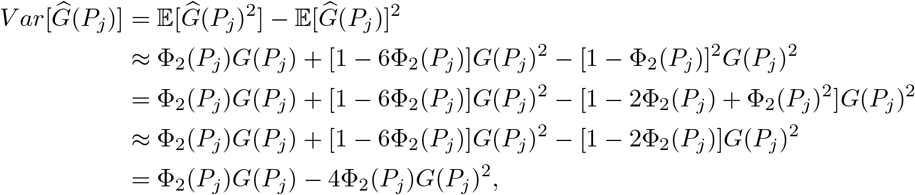

which gives

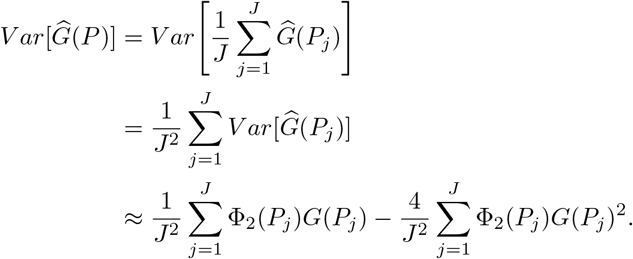

Recall that

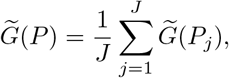

where

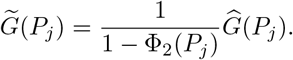

It follows that

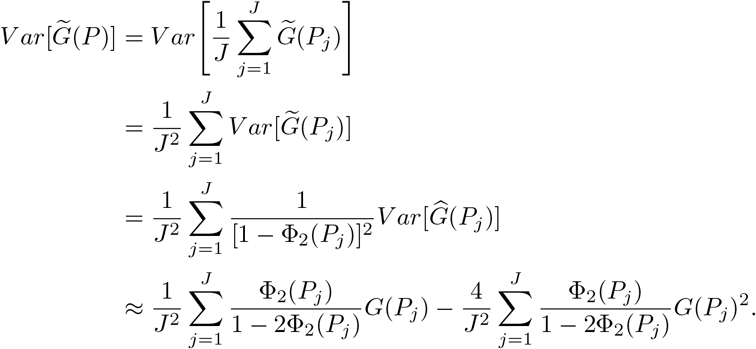

### Lemma 10.

Consider *J* independent polymorphic loci in populations *A* and *B* with respective parametric reference allele frequencies *a*_*j*_, *b*_*j*_ ∈ (0, 1), and suppose we take a random sample of *N* (*P*_*j*_) individuals at locus *j* in population *P* ∈ {*A, B*}, some of which may be related or inbred. Moreover, assume that no individual is related to more than one other individual, which makes the terms Φ_3_(*P*_*j*_), Φ_4_(*P*_*j*_), Φ_2,2_(*P*_*j*_), and Φ_2_(*P*_*j*_)^2^ negligible to Φ_2_(*P*_*j*_). Based on this simplifying assumption, the estimators 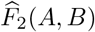 and 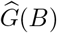 have an approximate covariance

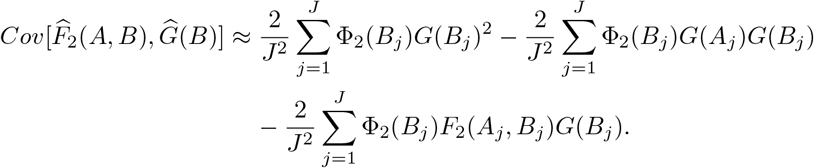

*Proof.* From the proofs of Lemma 1 and Proposition 2, we have

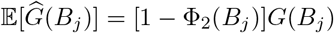

and

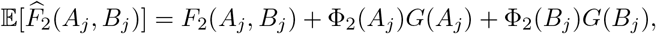

yielding

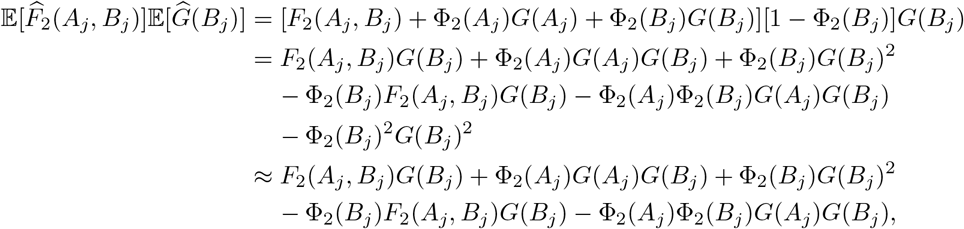

where we used the fact that Φ_2_(*B*_*j*_)^2^ is negligible compared to Φ_2_(*B*_*j*_) as an approximation. We also calculate

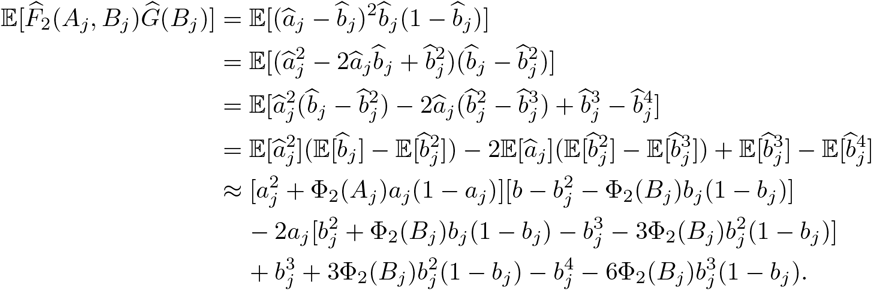

Recognizing that *G*(*a*_*j*_) = *a*_*j*_(1 − *a*_*j*_), *G*(*b*_*j*_) = *g_j_*(1 − *g_j_*), and 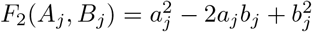, we have

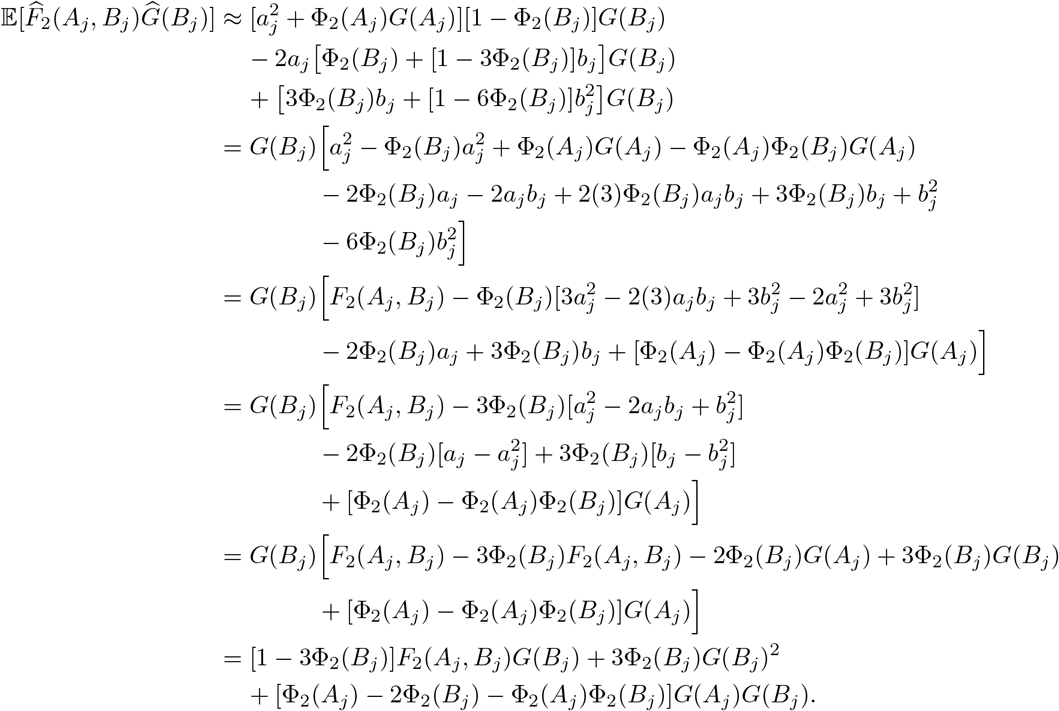

Therefore, we have that

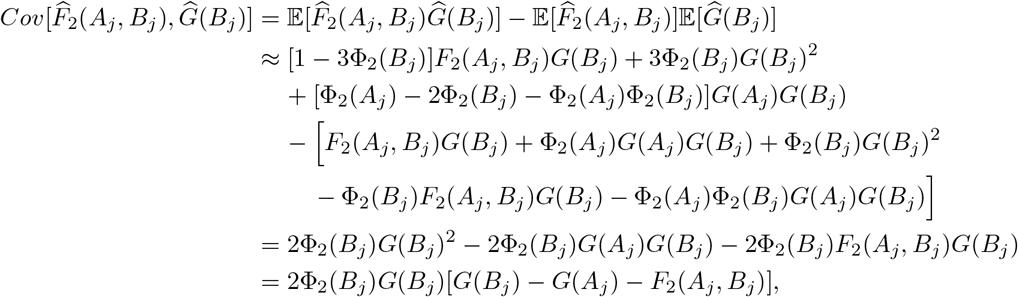

which gives

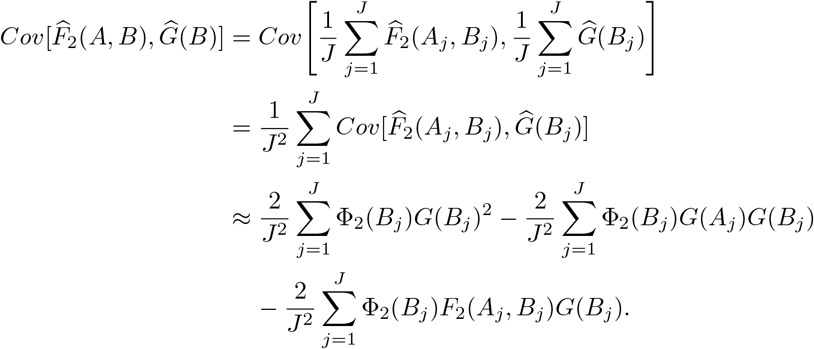

### Proposition 11.

Consider *J* independent polymorphic loci in a populations *A* and *B* with respective parametric reference allele frequencies *a*_*j*_, *b*_*j*_ ∈ (0, 1), and suppose we take a random sample of *N* (*P*_*j*_) individuals at locus *j* in population *P* ∈ {*A, B*}, some of which may be related or inbred. Moreover, assume that no individual is related to more than one other individual, which makes the terms Φ_3_(*P*_*j*_),

Φ_4_(*P*_*j*_), Φ_2,2_(*P*_*j*_), Φ_2_(*P*_*j*_)^2^, and Φ_2_(*A*_*j*_)Φ_2_(*B*_*j*_) negligible to Φ_2_(*P*_*j*_). Based on this simplifying assumption, the estimator 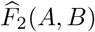 has approximate variance

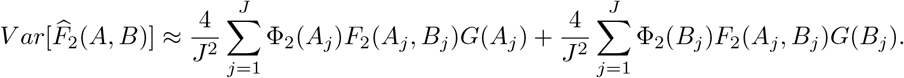

*Proof.* From the proof of Proposition 2, we have

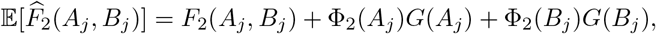

which gives

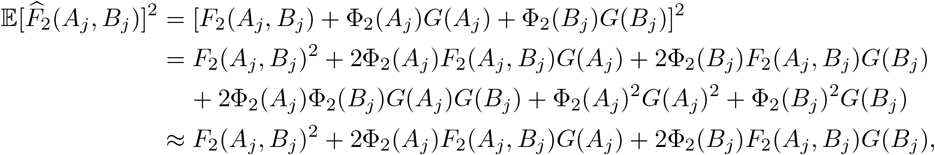

where we used the fact that Φ_2_(*A*_*j*_)^2^, Φ_2_(*B*_*j*_), and Φ_2_(*A*_*j*_)Φ_2_(*B*_*j*_) are negligible compared to Φ_2_(*A*_*j*_) and Φ_2_(*B*_*j*_) as an approximation. We also calculate

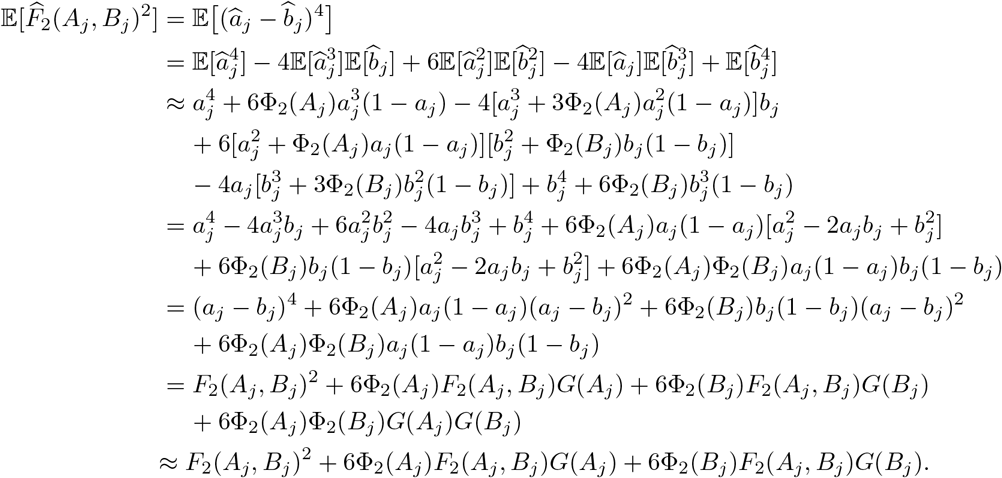

Therefore, we have that

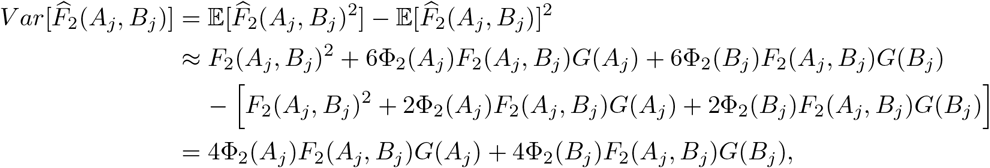

which gives

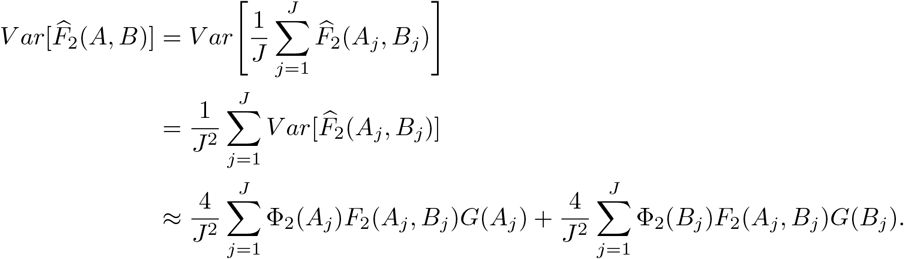

### Proposition 12.

Consider *J* independent polymorphic loci in a populations *A* and *B* with respective parametric reference allele frequencies *a*_*j*_, *b*_*j*_ ∈ (0, 1), and suppose we take a random sample of *N* (*P*_*j*_) individuals at locus *j* in population *P* ∈ {*A, B*}, some of which may be related or inbred. Moreover, assume that no individual is related to more than one other individual, which makes the terms Φ_3_(*P*_*j*_),

Φ_4_(*P*_*j*_), Φ_2,2_(*P*_*j*_), Φ_2_(*P*_*j*_)^2^, and Φ_2_(*A*_*j*_)Φ_2_(*B*_*j*_) negligible to Φ_2_(*P*_*j*_). Based on this simplifying assumption, the unbiased estimator 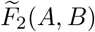 has approximate variance

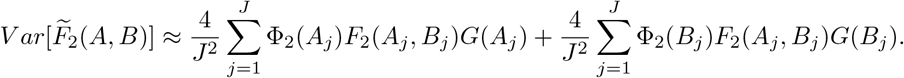

*Proof.* Recall that

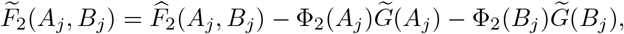

where 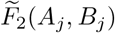 is an unbiased estimator for *F*_2_(*a*_*j*_, *b*_*j*_) and 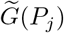 is an unbiased estimator of *G*(*P*_*j*_) for *P* ∈ {*A, B*} at locus *j* ∈ {1, 2, …, *J*}. Also, from the proof of Proposition 2, we have

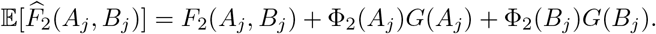

Therefore, we have that

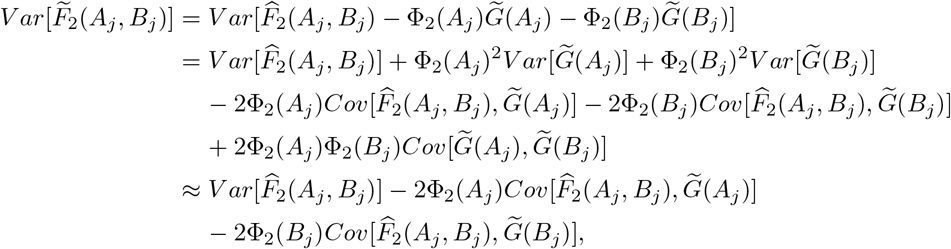

where we used the fact that Φ_2_(*A*_*j*_)^2^ and Φ_2_(*B*_*j*_)^2^ are negligible compared to Φ_2_(*A*_*j*_) and Φ_2_(*B*_*j*_) as an approximation, and where 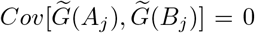 because drawing alleles in population *A* is independent of population *B*. Moreover, because 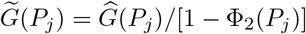, we have

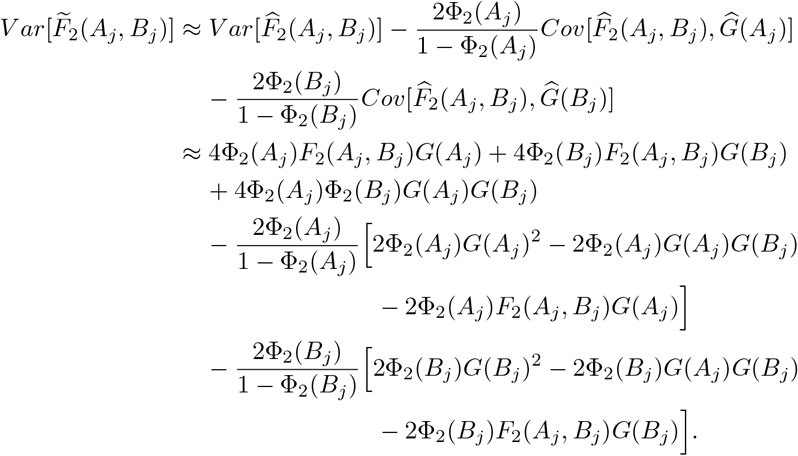

Recalling the assumption that Φ_2_(*A*_*j*_)^2^, Φ_2_(*B*_*j*_)^2^, and Φ_2_(*A*_*j*_)Φ_2_(*B*_*j*_) are negligible compared to Φ_2_(*A*_*j*_) and Φ_2_(*B*_*j*_), we have

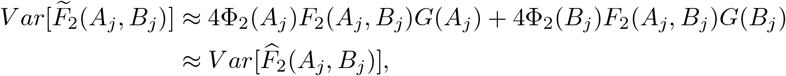

and it follows that

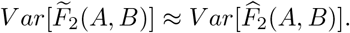

### Proposition 13.

Consider *J* independent polymorphic loci in populations *A*, *B*, and *C* with respective parametric reference allele frequencies *a*_*j*_, *b*_*j*_, *c*_*j*_ ∈ (0, 1), and suppose we take a random sample of *N* (*P*_*j*_) individuals at locus *j* in population *P* ∈ {*A, B, C*}, some of which may be related or inbred. Moreover, assume that no individual is related to more than one other individual, which makes the terms Φ_3_(*P*_*j*_), Φ_4_(*P*_*j*_), Φ_2,2_(*P*_*j*_), Φ_2_(*P*_*j*_)^2^, Φ_2_(*A*_*j*_)Φ_2_(*B*_*j*_), Φ_2_(*A*_*j*_)Φ_2_(*CB*_*j*_), and Φ_2_(*B*_*j*_)Φ_2_(*C*_*j*_) negligible to Φ_2_(*P*_*j*_). Based on this simplifying assumption, the estimator 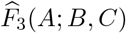 has approximate variance

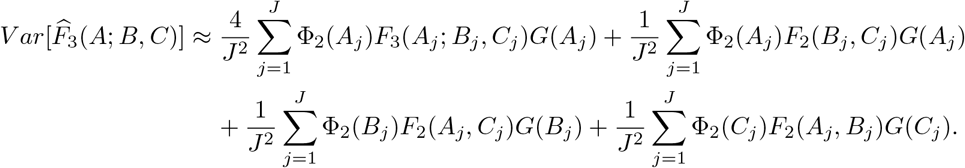

*Proof.* From the proof of Proposition 3, we have

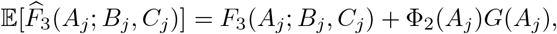

which gives

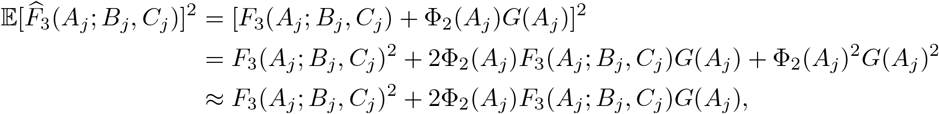

where we used the fact that Φ_2_(*A*_*j*_)^2^ is negligible compared to Φ_2_(*A*_*j*_) as an approximation. We also calculate

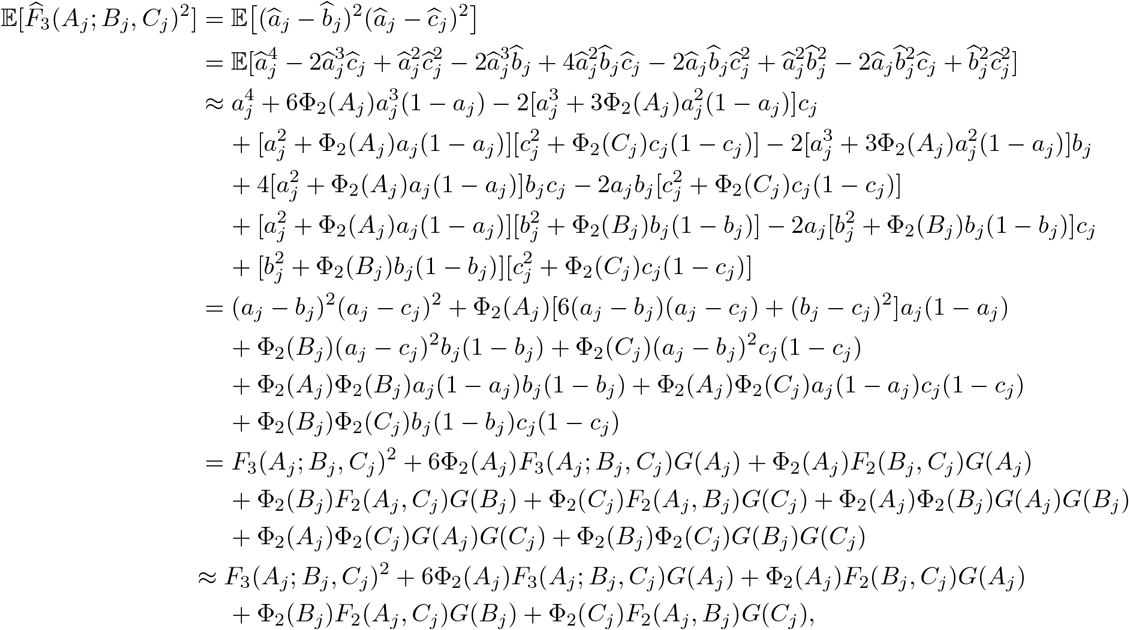

where we used the fact that Φ_2_(*A*_*j*_)Φ_2_(*B*_*j*_), Φ_2_(*A*_*j*_)Φ_2_(*C*_*j*_), and Φ_2_(*B*_*j*_)Φ_2_(*C*_*j*_) are negligible compared to Φ_2_(*A*_*j*_), Φ_2_(*B*_*j*_), and Φ_2_(*C*_*j*_) as an approximation. Therefore, we have that

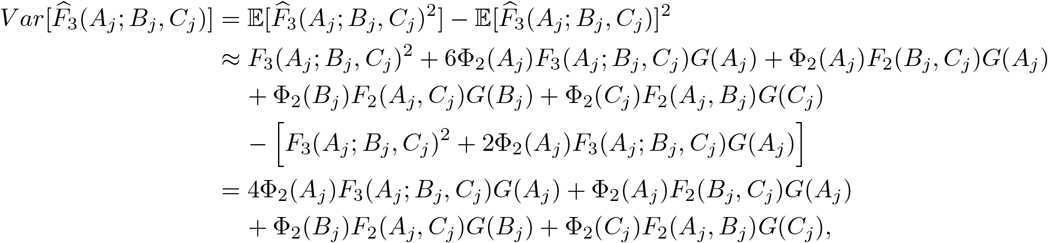

which gives

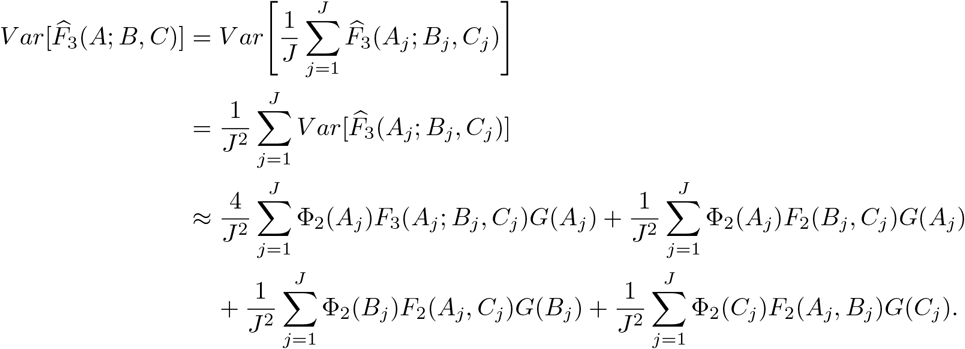

### Lemma 14.

Consider *J* independent polymorphic loci in populations *A*, *B*, and *c* with respective parametric reference allele frequencies *a*_*j*_, *b*_*j*_, *c*_*j*_ ∈ (0, 1), and suppose we take a random sample of *N* (*P*_*j*_) individuals at locus *j* in population *P* ∈ {*A, B, C*}, some of which may be related or inbred. Moreover, assume that no individual is related to more than one other individual, which makes the terms Φ_3_(*P*_*j*_), Φ_4_(*P*_*j*_), Φ_2,2_(*P*_*j*_),and Φ_2_(*P*_*j*_)^2^ negligible to Φ_2_(*P*_*j*_). Based on this simplifying assumption, the estimators 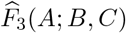 and 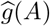 have an approximate covariance

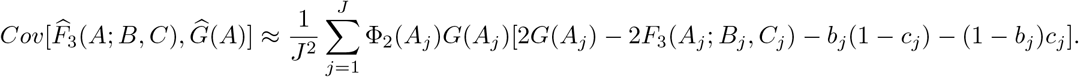

*Proof.* From the proofs of Lemma 1 and Proposition 3, we have

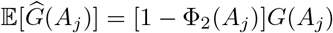

and

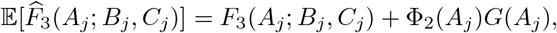

yielding

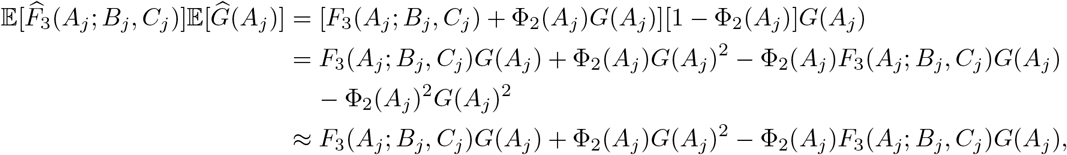

where we used the fact that Φ_2_(*A*_*j*_)^2^ is negligible compared to Φ_2_(*A*_*j*_) as an approximation. We also calculate

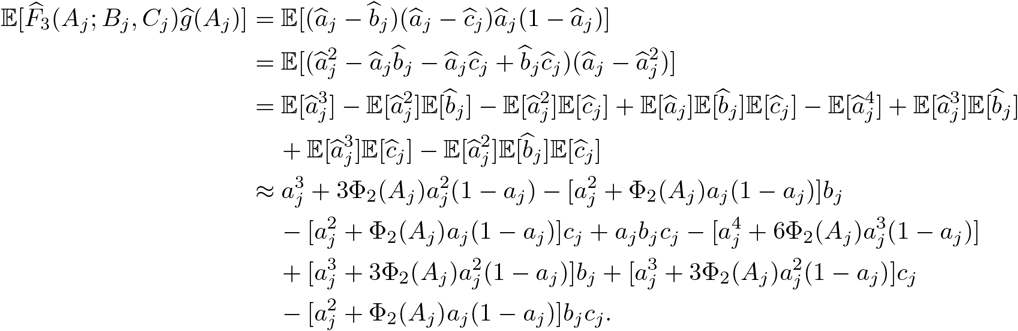

Recognizing that *G*(*a*_*j*_) = *a*_*j*_(1 − *a*_*j*_) and 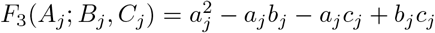, we have

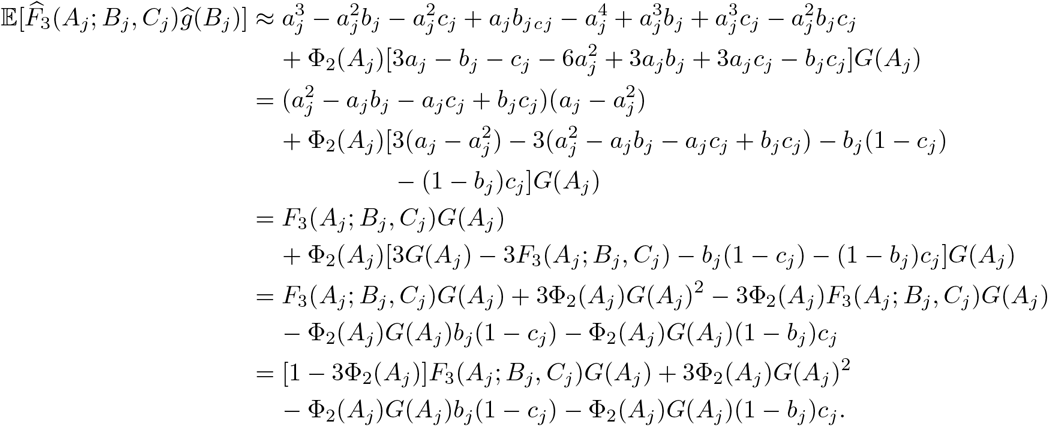

Therefore, we have that

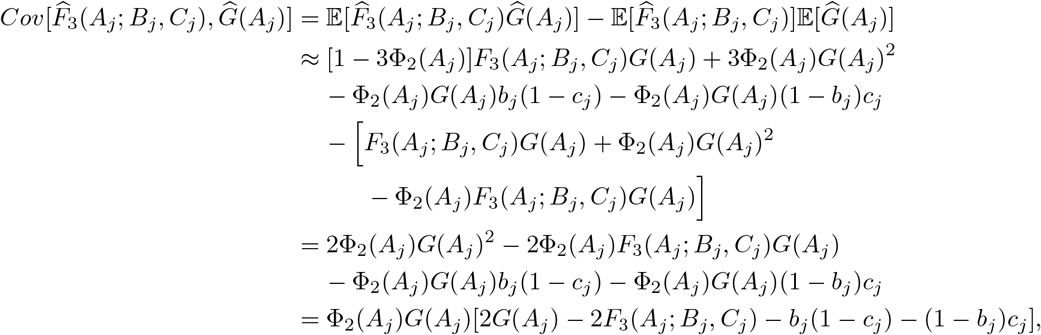

which gives

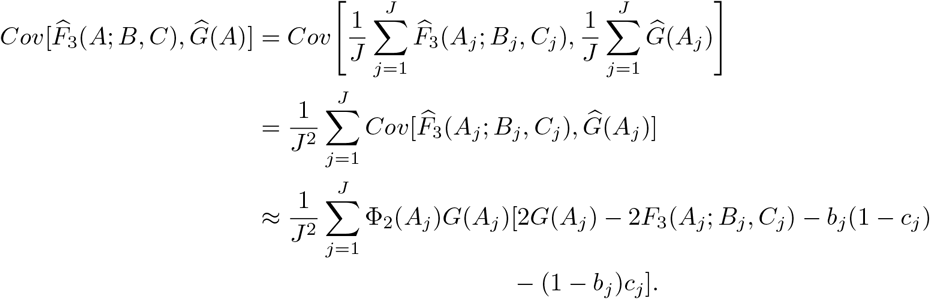

### Proposition 15.

Consider *J* independent polymorphic loci in populations *A*, *B*, and *C* with respective parametric reference allele frequencies *a*_*j*_, *b*_*j*_, *c*_*j*_ ∈ (0, 1), and suppose we take a random sample of *N* (*P*_*j*_) individuals at locus *j* in population *P* ∈ {*A, B, C*}, some of which may be related or inbred. Moreover, assume that no individual is related to more than one other individual, which makes the terms Φ_3_(*P*_*j*_), Φ_4_(*P*_*j*_), Φ_2,2_(*P*_*j*_), Φ_2_(*P*_*j*_)^2^, Φ_2_(*A*_*j*_)Φ_2_(*B*_*j*_), Φ_2_(*A*_*j*_)Φ_2_(*C*_*j*_), and Φ_2_(*B*_*j*_)Φ_2_(*C*_*j*_) negligible to Φ_2_(*P*_*j*_). Based on this simplifying assumption, the unbiased estimator 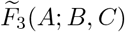 has approximate variance

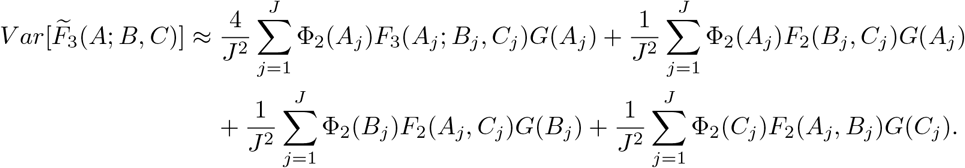

*Proof.* Recall that

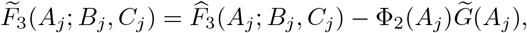

where 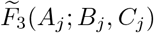 is an unbiased estimator for *F*_3_(*A*_*j*_; *B*_*j*_, *C*_*j*_) and 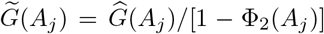 is an unbiased estimator of *G*(*A*_*j*_) at locus *j* ∈ {1, 2, …, *J*}. Also, from the proof of Proposition 3, we have

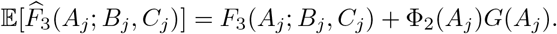

Therefore, we have that

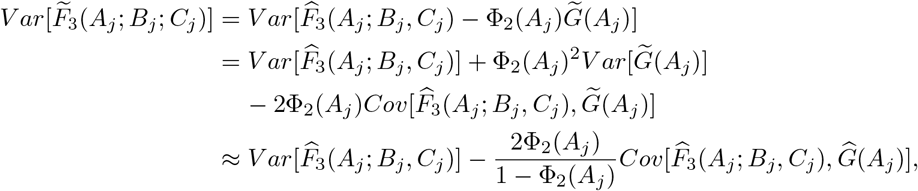

where we used the fact that Φ_2_(*A*_*j*_)^2^ is negligible compared to Φ_2_(*A*_*j*_) as an approximation. Moreover, because 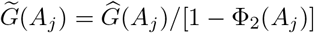, we have

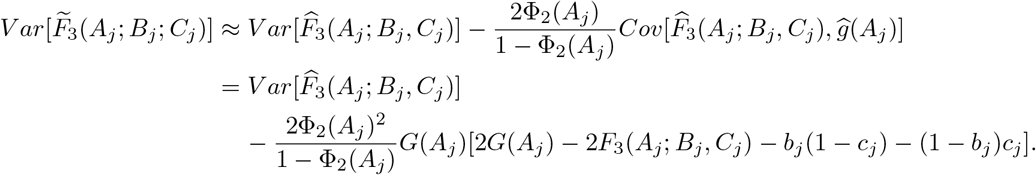

Recalling the assumption that Φ_2_(*A*_*j*_)^2^ is negligible compared to Φ_2_(*A*_*j*_), we have

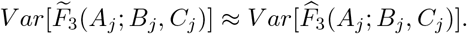

### Proposition 16.

Consider *J* independent polymorphic loci in populations *A*, *B*, *C*, and *D* with respective parametric reference allele frequencies *a*_*j*_, *b*_*j*_, *c*_*j*_, *d*_*j*_ ∈ (0, 1), and suppose we take a random sample of *N* (*P*_*j*_) individuals at locus *j* in population *P* ∈ {*A, B, C, D*}, some of which may be related or inbred. Moreover, assume that no individual is related to more than one other individual, which makes the terms Φ_3_(*P*_*j*_), Φ_4_(*P*_*j*_), Φ_2,2_(*P*_*j*_), Φ_2_(*P*_*j*_)^2^, Φ_2_(*A*_*j*_)Φ_2_(*C*_*j*_), Φ_2_(*A*_*j*_)Φ_2_(*D*_*j*_), Φ_2_(*B*_*j*_)Φ_2_(*C*_*j*_), and Φ_2_(*B*_*j*_)Φ_2_(*D*_*j*_) negligible to Φ_2_(*P*_*j*_). Based on this simplifying The unbiased estimator 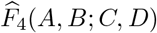 has approximate variance

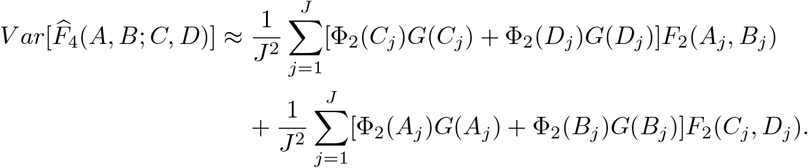

*Proof.* From the proofs of Propositions 2 and 5, we have

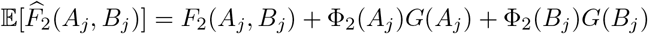

and

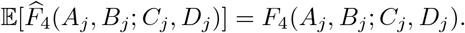

We calculate

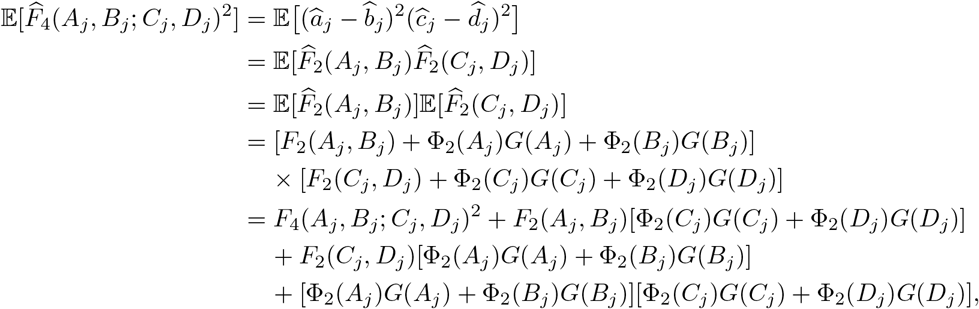

where we use the identity that

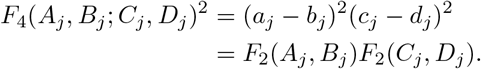

Therefore, we have that

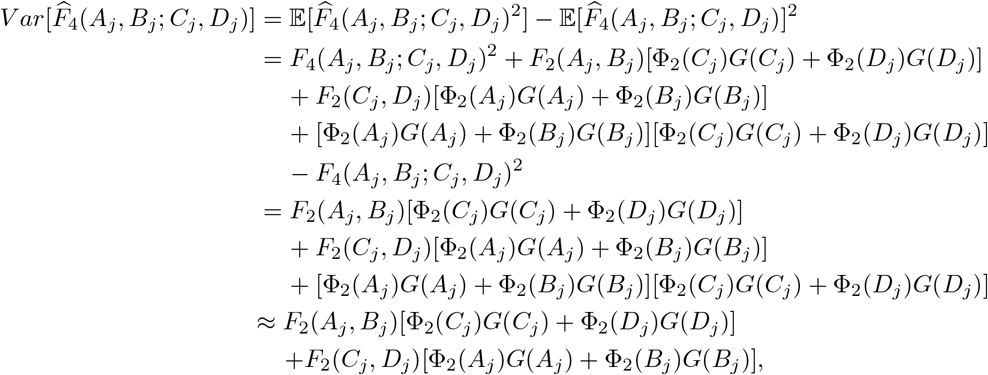

where we used the fact that Φ_2_(*A*_*j*_)Φ_2_(*C*_*j*_), Φ_2_(*A*_*j*_)Φ_2_(*D*_*j*_), Φ_2_(*B*_*j*_)Φ_2_(*C*_*j*_), and Φ_2_(*C*_*j*_)Φ_2_(*D*_*j*_) are negligible compared to Φ_2_(*A*_*j*_), Φ_2_(*B*_*j*_), Φ_2_(*C*_*j*_) and Φ_2_(*D*_*j*_) as an approximation. It follows that

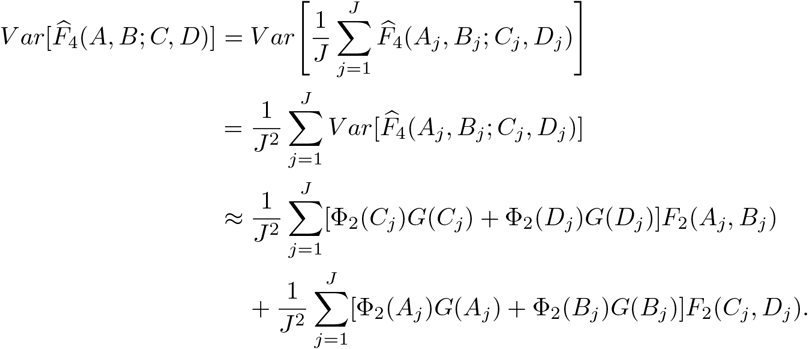

Following Wolter (2007), we have that an approximation to the variance of the ratio estimator *X/Y* is

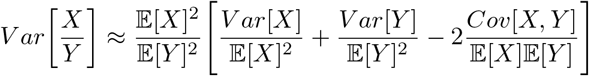

### Proposition 17.

Consider *J* polymorphic loci in populations *A*, *B*, and *C* with respective parametric reference allele frequencies *a*_*j*_, *b*_*j*_, *c*_*j*_ ∈ (0, 1), and suppose we take a random sample of *N* (*P*_*j*_) individuals at locus *j* in population *P* ∈ {*A, B, C*}, some of which may be related or inbred. Moreover, assume that no individual is related to more than one other individual, which makes the terms Φ_3_(*P*_*j*_), Φ_4_(*P*_*j*_), Φ_2,2_(*P*_*j*_), Φ_2_(*P*_*j*_)^2^, Φ_2_(*A*_*j*_)Φ_2_(*B*_*j*_), Φ_2_(*A*_*j*_)Φ_2_(*C*_*j*_), and Φ_2_(*B*_*j*_)Φ_2_(*C*_*j*_) negligible to Φ_2_(*P*_*j*_). Based on this simplifying assumption, the ratio estimator 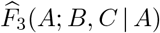 has approximate variance

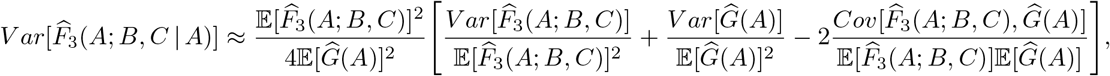

where the expectations are

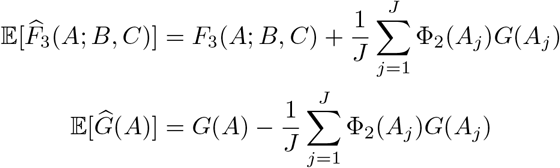

the variances are

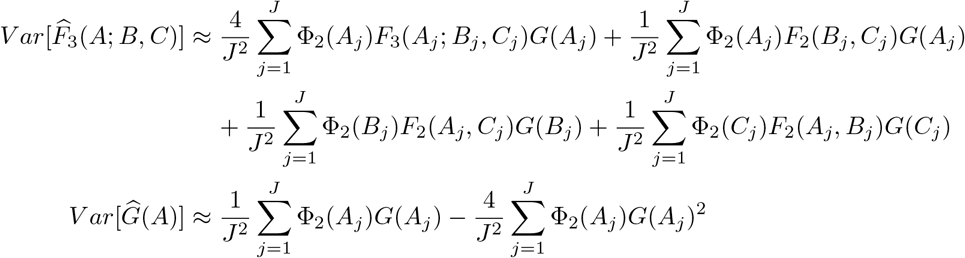

and the covariance is

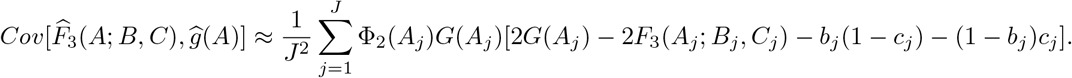

*Proof.* Recall that

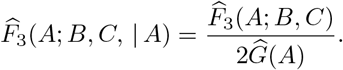

Assuming that 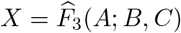 and 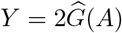, following the approximation in Wolter (2007) we have

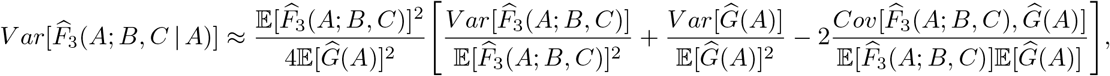

where 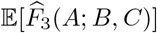 is given in Proposition 3, 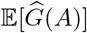 in Lemma 1, 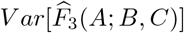 in Proposition 13, 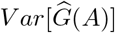 in Lemma 9, and 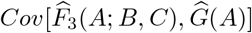 in Lemma 14.

### Lemma 18.

Consider *J* independent polymorphic loci in populations *A*, *B*, and *C* with respective parametric reference allele frequencies *a*_*j*_, *b*_*j*_, *c*_*j*_ ∈ (0, 1), and suppose we take a random sample of *N* (*P*_*j*_) individuals at locus *j* in population *P* ∈ {*A, B, C*}, some of which may be related or inbred. Moreover, assume that no individual is related to more than one other individual, which makes the terms Φ_3_(*P*_*j*_), Φ_4_(*P*_*j*_), Φ_2,2_(*P*_*j*_), and Φ_2_(*P*_*j*_)^2^ negligible to Φ_2_(*P*_*j*_). Based on this simplifying assumption, the unbiased estimators 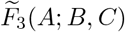 and 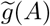 have an approximate covariance

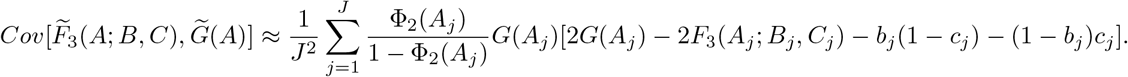

*Proof.* Recall that

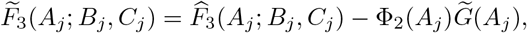

where 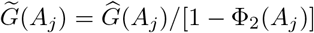. It follows that

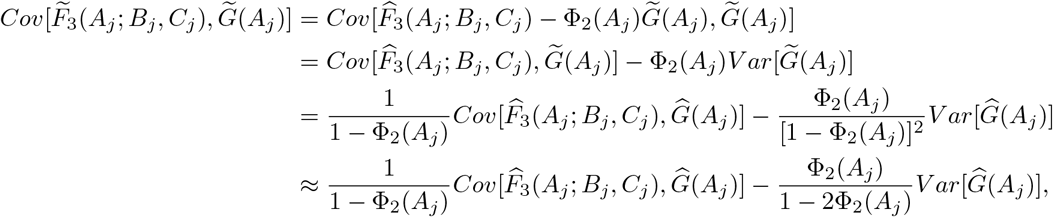

where we used the fact that Φ_2_(*A*_*j*_)^2^ is negligible compared to Φ_2_(*A*_*j*_) as an approximation. From the proofs of Lemmas 9 and 14, we have

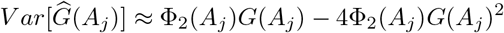

and

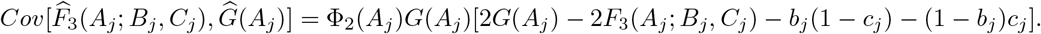

Assuming that Φ_2_(*A*_*j*_)^2^ is negligible compared to Φ_2_(*A*_*j*_), we have that

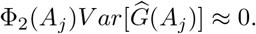

We therefore have that

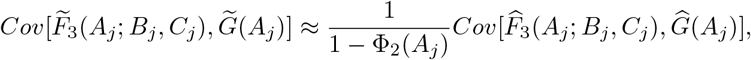

and thus by independence of loci we have

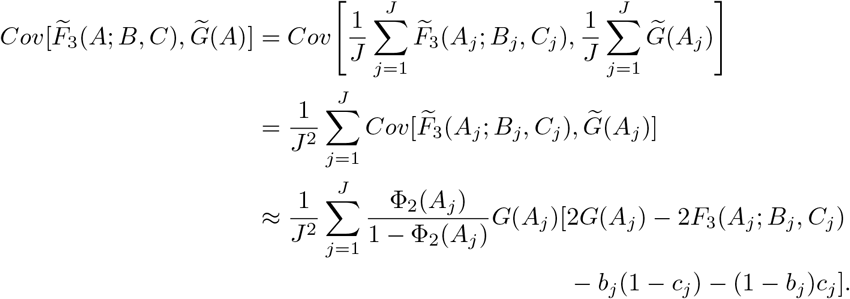

### Proposition 19.

Consider *J* polymorphic loci in populations *A*, *B*, and *C* with respective parametric reference allele frequencies *a*_*j*_, *b*_*j*_, *c*_*j*_ ∈ (0, 1), and suppose we take a random sample of *N* (*P*_*j*_) individuals at locus *j* in population *P* ∈ {*A, B, C*}, some of which may be related or inbred. Moreover, assume that no individual is related to more than one other individual, which makes the terms Φ_3_(*P*_*j*_), Φ_4_(*P*_*j*_), Φ_2,2_(*P*_*j*_), Φ_2_(*P*_*j*_)^2^, Φ_2_(*A*_*j*_)Φ_2_(*B*_*j*_), Φ_2_(*A*_*j*_)Φ_2_(*C*_*j*_), and Φ_2_(*B*_*j*_)Φ_2_(*C*_*j*_) negligible to Φ_2_(*P*_*j*_). Based on this simplifying assumption, the approximately unbiased ratio estimator 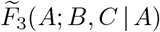 has approximate variance

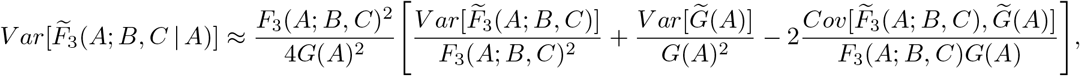

where the variances are

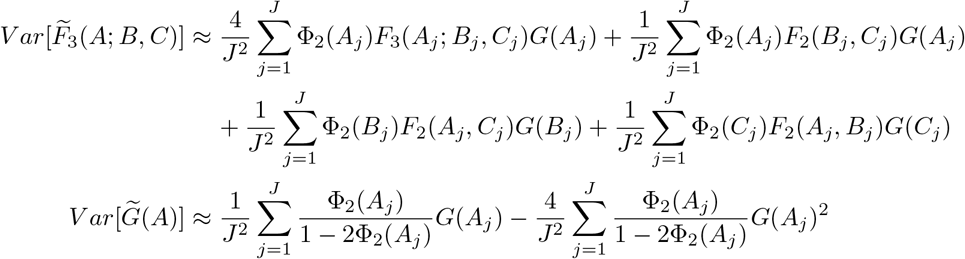

and the covariance is

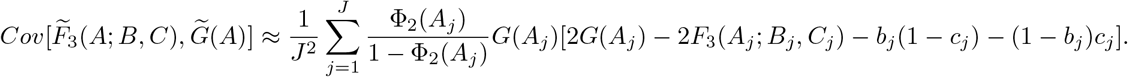

*Proof.* Recall that

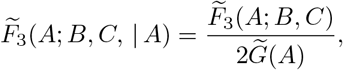

where 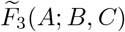 is an unbiased estimator for *F*_3_(*A*; *B, C*) and 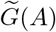 is an unbiased estimator of *G*(*A*). Assuming that 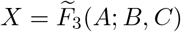 and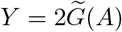, following the approximation in Wolter (2007) we have

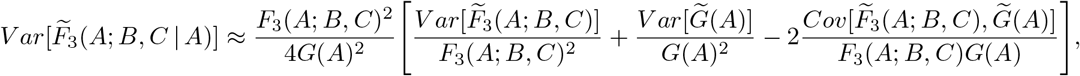

where 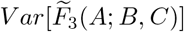 is given in Proposition 15, 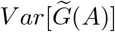 in Lemma 9, and 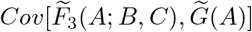 in Lemma 18.

### Lemma 20.

Consider *J* independent polymorphic loci in populations *A*, *B*, *C*, and *D* with respective parametric reference allele frequencies *a*_*j*_, *b*_*j*_, *c*_*j*_, *d*_*j*_ ∈ (0, 1), and suppose we take a random sample of *N* (*P*_*j*_) individuals at locus *j* in population *P* ∈ {*A, B, C, D*}, some of which may be related or inbred. Moreover, assume that no individual is related to more than one other individual, which makes the terms Φ_3_(*P*_*j*_), Φ_4_(*P*_*j*_), Φ_2,2_(*P*_*j*_), and Φ_2_(*P*_*j*_)^2^ negligible to Φ_2_(*P*_*j*_). Based on this simplifying assumption, the estimators 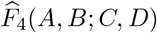 and 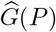, *P* ∈ {*A, B, C, D*}, have approximate covariances

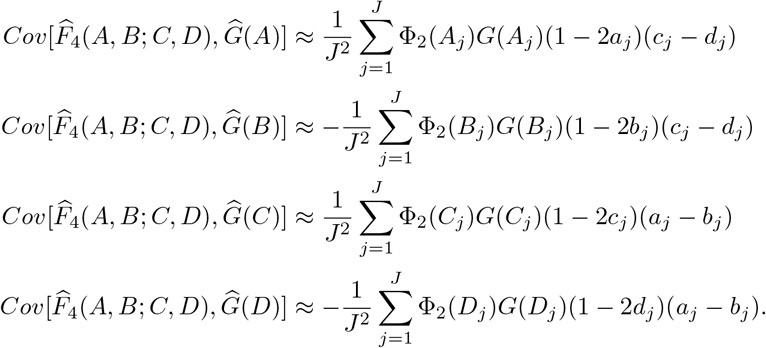

*Proof.* From the proofs of Lemma 1 and Proposition 5, we that have

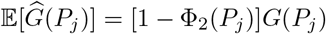

and

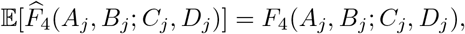

yielding

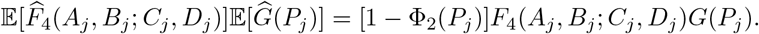

We first calculate

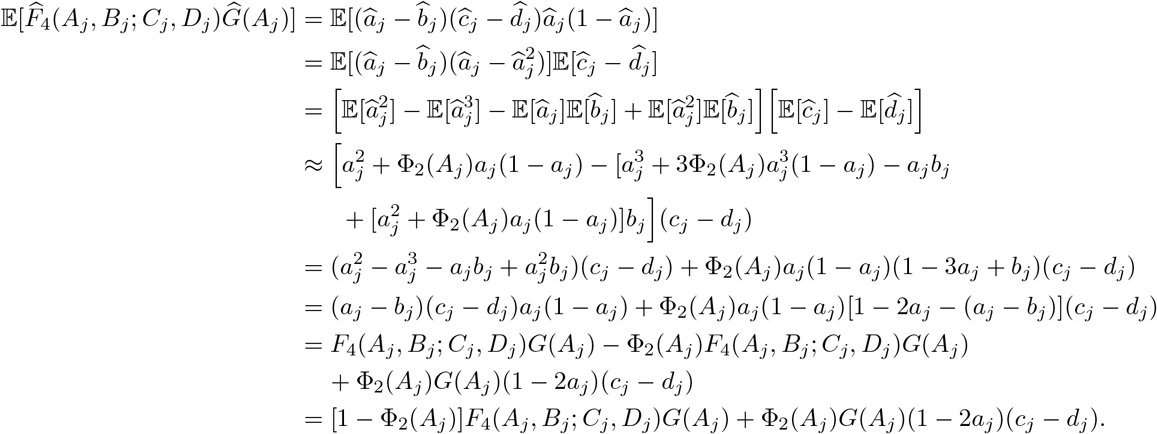

Hence, we have that

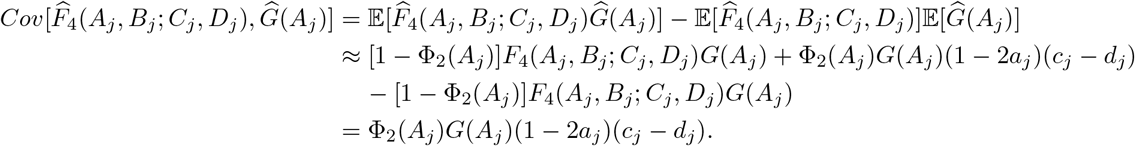

Similarly, we have that

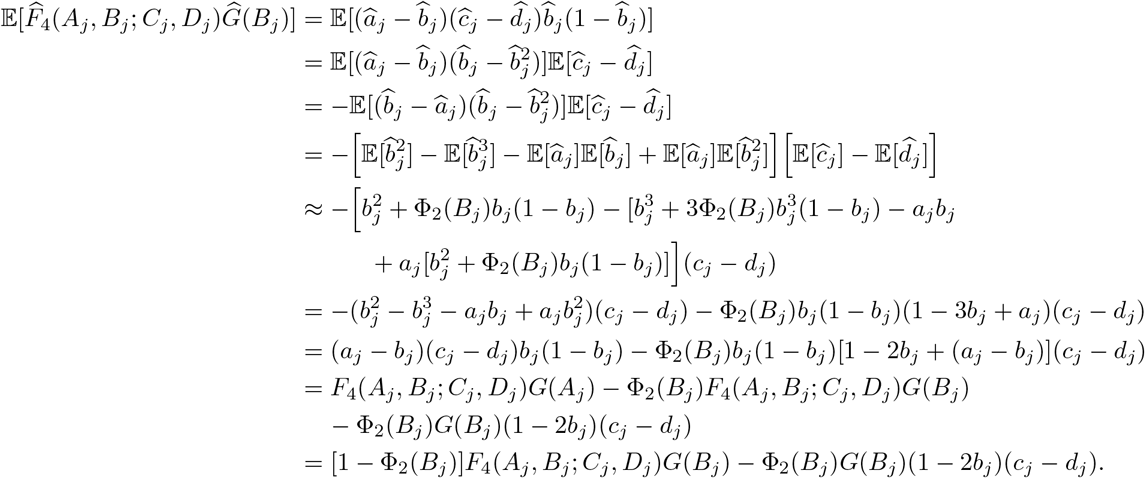

Hence, we have that

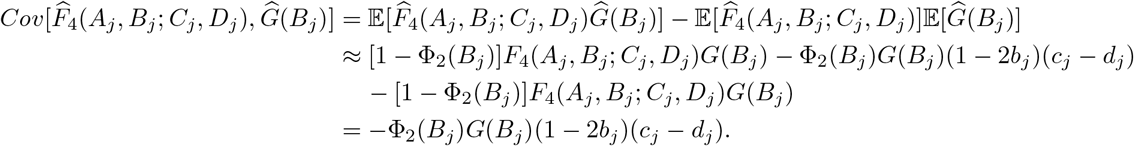

Parallel to the derivation for *P* = *A*, we have

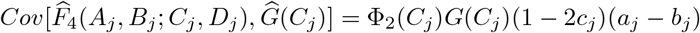

and parallel to the derivation for *P* = *B*, we have

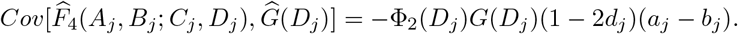

We know that by independence of loci we have

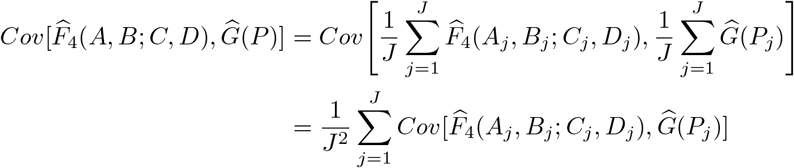

which gives

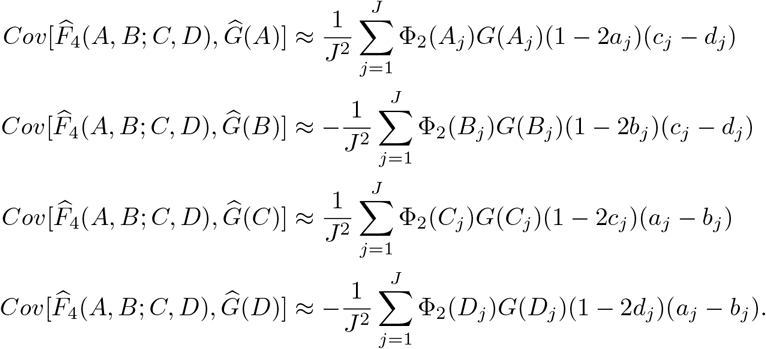

### Proposition 21.

Consider *J* polymorphic loci in populations *A*, *B*, *C*, and *D* with respective parametric reference allele frequencies *a*_*j*_, *b*_*j*_, *c*_*j*_, *d*_*j*_ ∈ (0, 1), and suppose we take a random sample of *N* (*P*_*j*_) individuals at locus *j* in population *P* ∈ {*A, B, C, D*}, some of which may be related or inbred. Moreover, assume that no individual is related to more than one other individual, which makes the terms Φ_3_(*P*_*j*_), Φ_4_(*P*_*j*_), Φ_2,2_(*P*_*j*_), and Φ_2_(*P*_*j*_)^2^ negligible to Φ_2_(*P*_*j*_). Based o this simplifying assumption, the ratio estimator 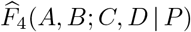 has approximate variance

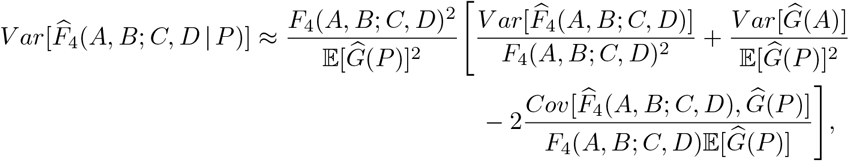

where the expectation is

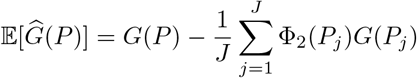

the variances are

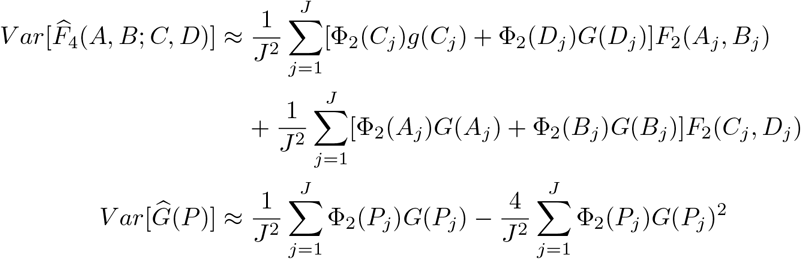

and the covariances are

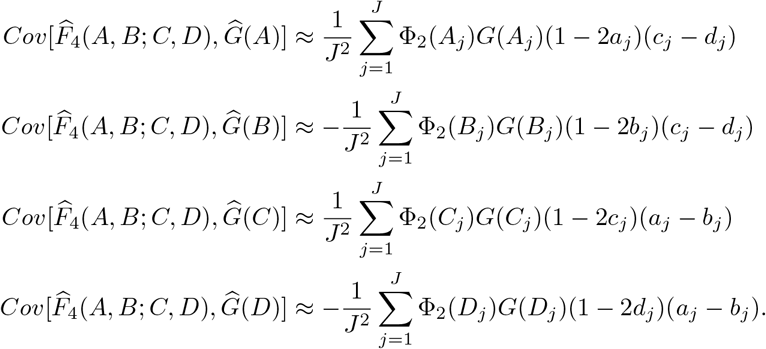

*Proof.* Recall that

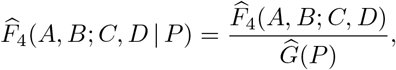

where 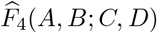 is an unbiased estimator for *F*_4_(*A, B*; *C, D*). Assuming that 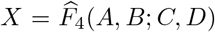 and 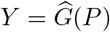, following the approximation in Wolter (2007) we have

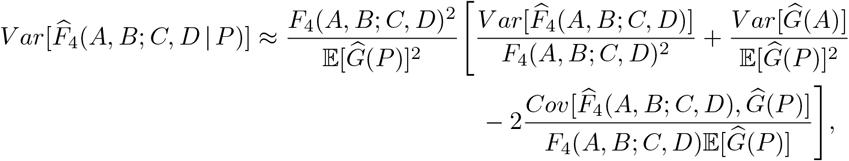

where 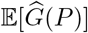 is given in Lemma 1, 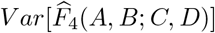 in Proposition 16, 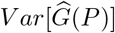 in Lemma 9, and 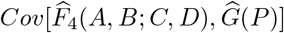 in Lemma 20 for each population *P* ∈ {*A, B, C, D*}.

### Lemma 22.

Consider *J* independent polymorphic loci in populations *A*, *B*, *C*, and *D* with respective parametric reference allele frequencies *a*_*j*_, *b*_*j*_, *c*_*j*_, *d*_*j*_ ∈ (0, 1), and suppose we take a random sample of *N* (*P*_*j*_) individuals at locus *j* in population *P* ∈ {*A, B, C, D*}, some of which may be related or inbred. Moreover, assume that no individual is related to more than one other individual, which makes the terms Φ_3_(*P*_*j*_), Φ_4_(*P*_*j*_), Φ_2,2_(*P*_*j*_), and Φ_2_(*P*_*j*_)^2^ negligible to Φ_2_(*P*_*j*_). Based on this simplifying assumption, the unbiased estimators 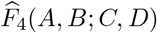 and 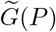, *P* ∈ {*A, B, C, D*}, have approximate covariances

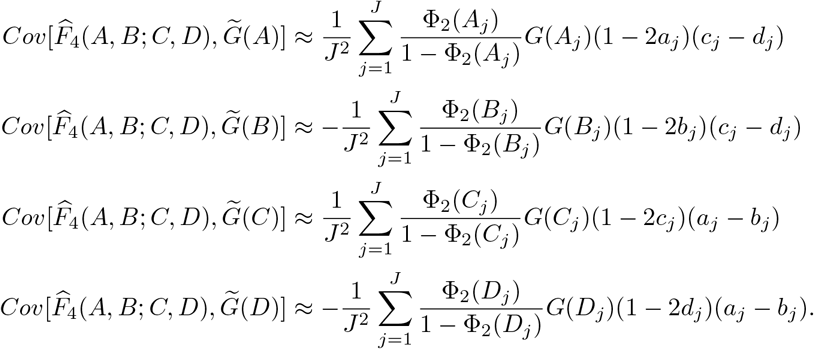

*Proof.* Recall that 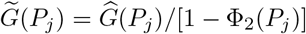. It follows that

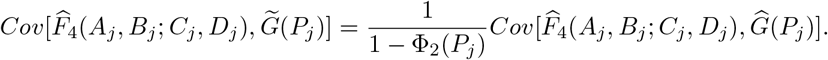

From the proof of Lemma 20, we have

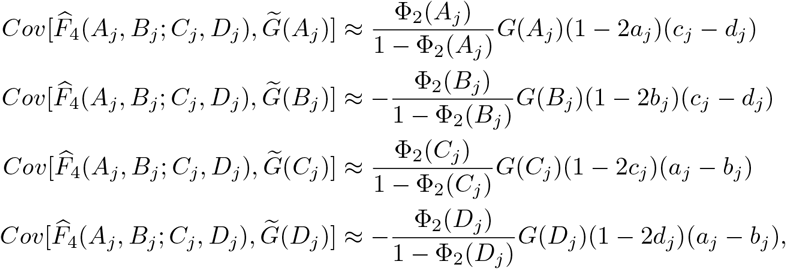

yielding

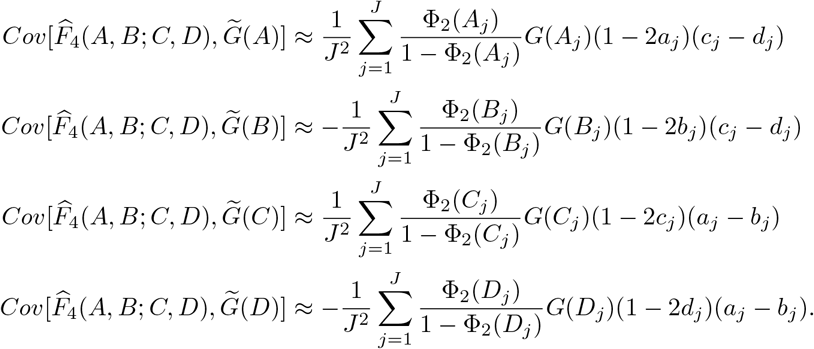

### Proposition 23.

Consider *J* polymorphic loci in populations *A*, *B*, *C*, and *D* with respective parametric reference allele frequencies *a*_*j*_, *b*_*j*_, *c*_*j*_, *d*_*j*_ ∈ (0, 1), and suppose we take a random sample of *N* (*P*_*j*_) individuals at locus *j* in population *P* ∈ {*A, B, C, D*}, some of which may be related or inbred. Moreover, assume that no individual is related to more than one other individual, which makes the terms Φ_3_(*P*_*j*_), Φ_4_(*P*_*j*_), Φ_2,2_(*P*_*j*_), and Φ_2_(*P*_*j*_)^2^ negligible to Φ_2_(*P*_*j*_). Based on this simplifying assumption, the approximately unbiased ratio estimator 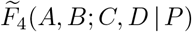 has approximate variance

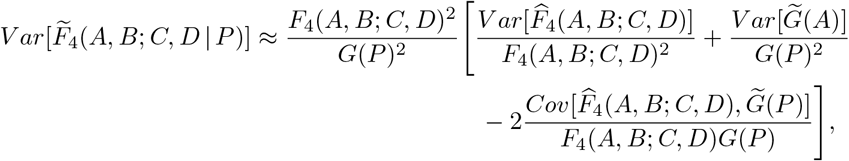

where the variances are

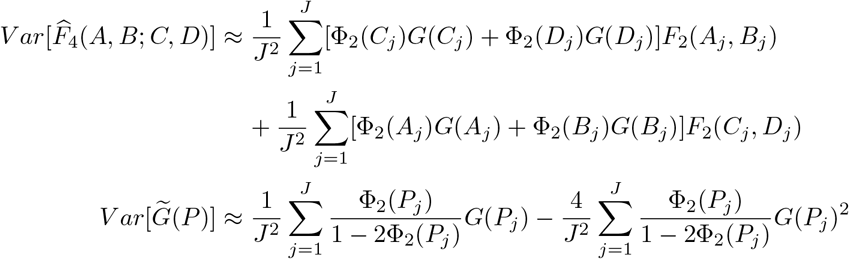

and the covariances are

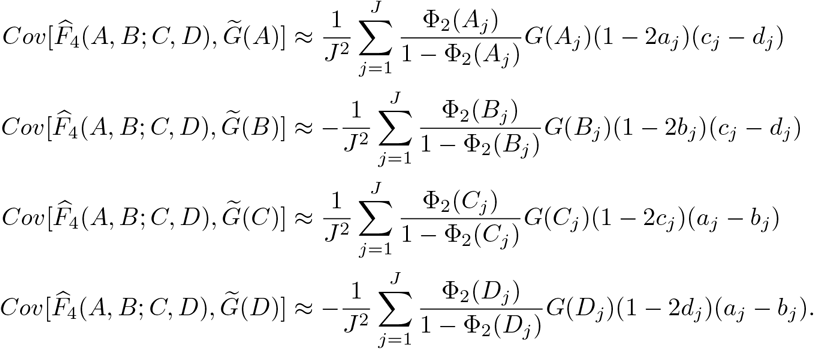

*Proof.* Recall that

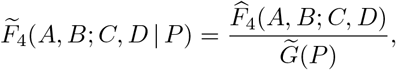

where 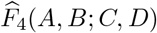 is an unbiased estimator for *F*_4_(*A, B*; *C, D*) and 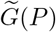 is an unbiased estimator of *G*(*P*). Assuming that 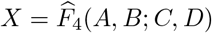 and 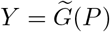, following the approximation in Wolter (2007) we have

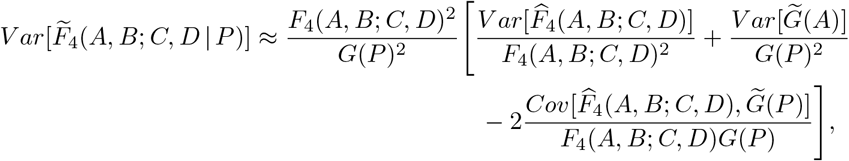

where 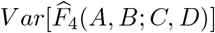 is given in Proposition 16, 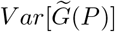 in Lemma 9, and 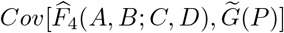 in Lemma 22 for each population *P* ∈ {*A, B, C, D*}.

### Lemma 24.

Consider *J* independent polymorphic loci in populations *A*, *B*, *C*, and *D* with respective parametric reference allele frequencies *a*_*j*_, *b*_*j*_, *c*_*j*_, *d*_*j*_ ∈ (0, 1), and suppose we take a random sample of *N* (*P*_*j*_) individuals at locus *j* in population *P* ∈ {*A, B, C, D*}, some of which may be related or inbred. Moreover, assume that no individual is related to more than one other individual, which makes the terms Φ_3_(*P*_*j*_), Φ_4_(*P*_*j*_), Φ_2,2_(*P*_*j*_), Φ_2_(*P*_*j*_)^2^, Φ_2_(*A*_*j*_)Φ_2_(*B*_*j*_), Φ_2_(*A*_*j*_)Φ_2_(*C*_*j*_), Φ_2_(*A*_*j*_)Φ_2_(*D*_*j*_), Φ_2_(*B*_*j*_)Φ_2_(*C*_*j*_), Φ_2_(*B*_*j*_)Φ_2_(*D*_*j*_), and Φ_2_(*C*_*j*_)Φ_2_(*D*_*j*_) negligible to Φ_2_(*P*_*j*_). Based on this simplifying assumption, the unbiased estimator 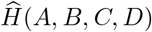 has approximate variance

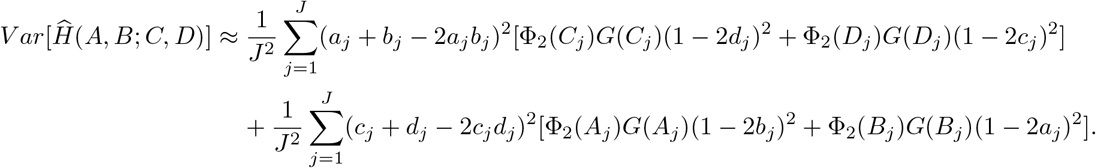

*Proof.* From the proof of Lemma 8, we have that

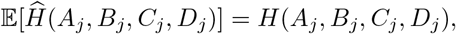

yielding

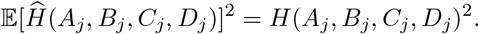

We first calculate

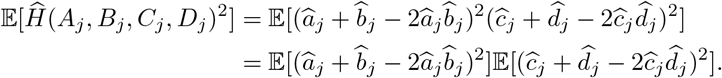

We compute the first term as

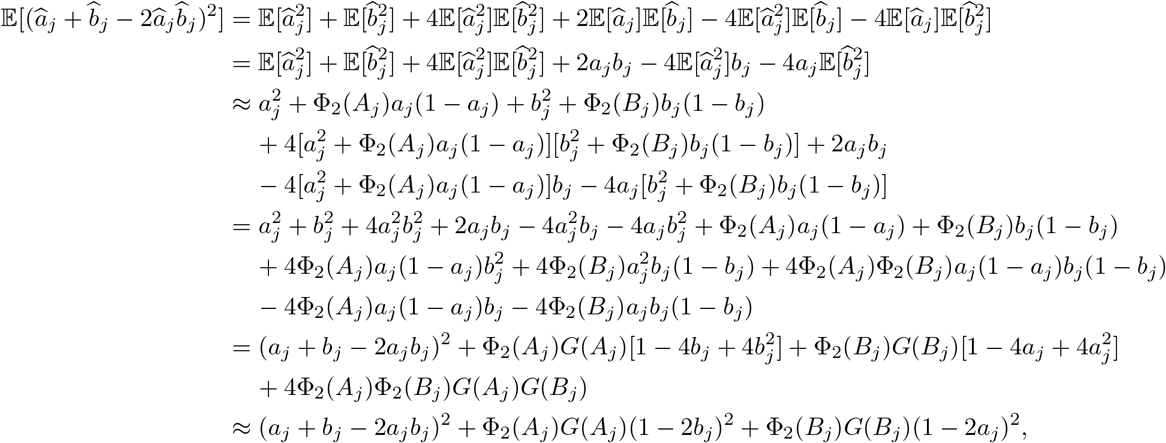

where we used the fact that Φ_2_(*A*_*j*_)Φ_2_(*B*_*j*_) is negligible compared to Φ_2_(*A*_*j*_) and Φ_2_(*B*_*j*_) as an approximation. Using a similar argument we have that

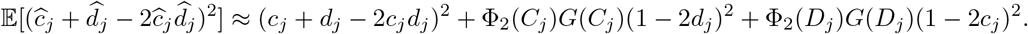

Hence, we have that

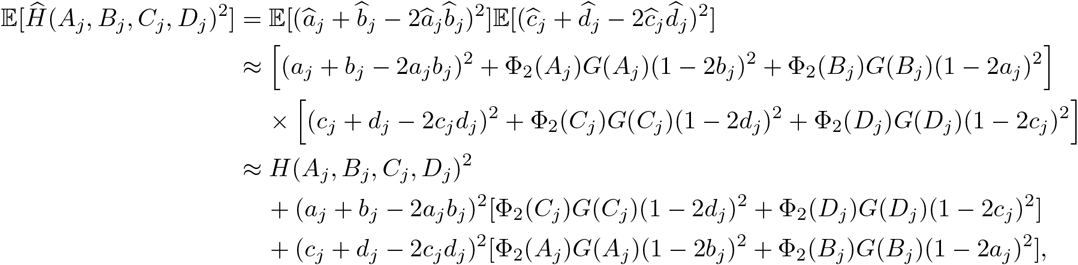

where we used the fact that Φ_2_(*A*_*j*_)Φ_2_(*C*_*j*_), Φ_2_(*A*_*j*_)Φ_2_(*D*_*j*_), Φ_2_(*B*_*j*_)Φ_2_(*C*_*j*_), and Φ_2_(*B*_*j*_)Φ_2_(*D*_*j*_) are negligible compared to Φ_2_(*A*_*j*_), Φ_2_(*B*_*j*_), Φ_2_(*C*_*j*_), and Φ_2_(*D*_*j*_) as an approximation. Putting it together, we have

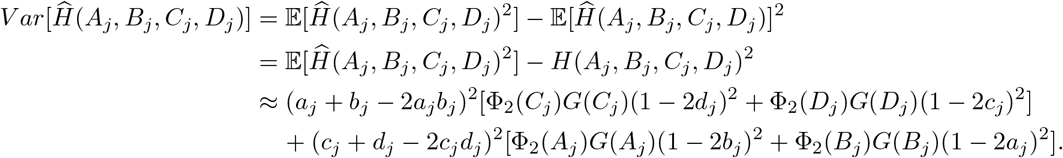

Given the assumption of independent loci, we have

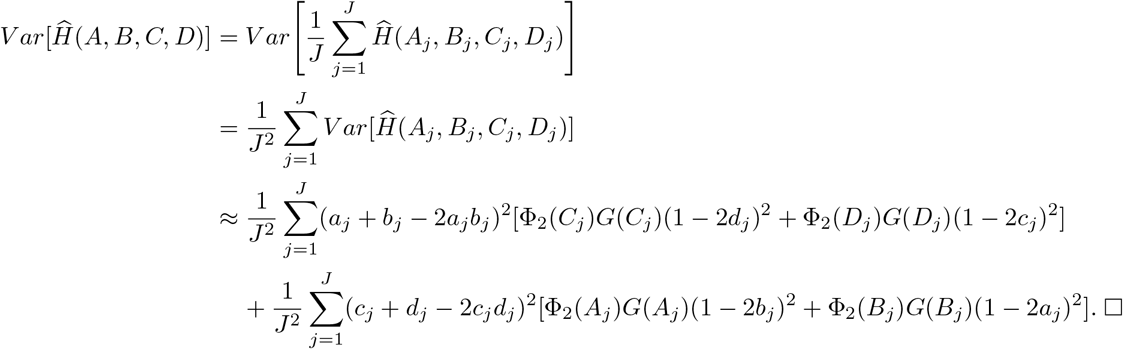

### Lemma 25.

Consider *J* independent polymorphic loci in populations *A*, *B*, *C*, and *D* with respective parametric reference allele frequencies *a*_*j*_, *b*_*j*_, *c*_*j*_, *d*_*j*_ ∈ (0, 1), and suppose we take a random sample of *N* (*P*_*j*_) individuals at locus *j* in population *P* ∈ {*A, B, C, D*}, some of which may be related or inbred. Moreover, assume that no individual is related to more than one other individual, which makes the terms Φ_3_(*P*_*j*_), Φ_4_(*P*_*j*_), Φ_2,2_(*P*_*j*_), Φ_2_(*P*_*j*_)^2^, Φ_2_(*A*_*j*_)Φ_2_(*C*_*j*_), Φ_2_(*A*_*j*_)Φ_2_(*D*_*j*_), Φ_2_(*B*_*j*_)Φ_2_(*C*_*j*_), and Φ_2_(*B*_*j*_)Φ_2_(*D*_*j*_) negligible to Φ_2_(*P*_*j*_). Based on this simplifying assumption, the unbiased estimators 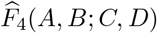 and 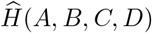 have approximate covariance

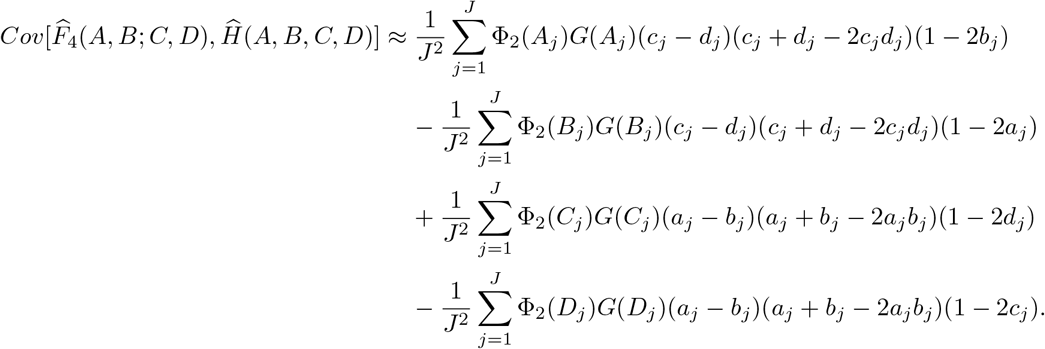

*Proof.* From the proofs of Proposition 5 and Lemma 8, we that have

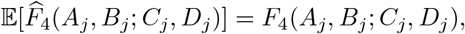

and

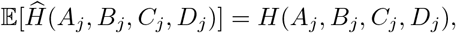

yielding

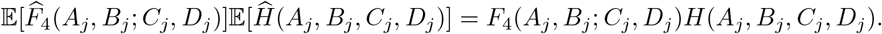

We first calculate

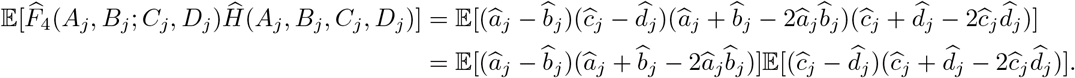

We compute the first term as

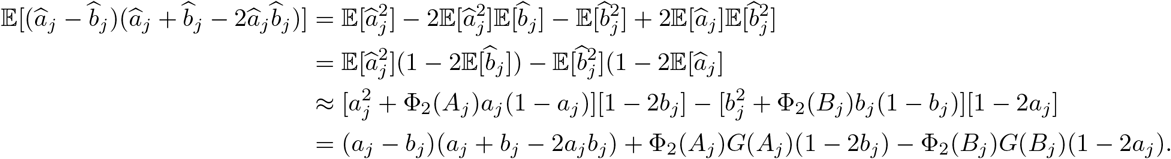

Using a similar argument, we have that

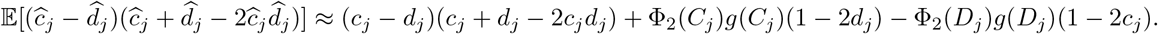

Hence, we have that

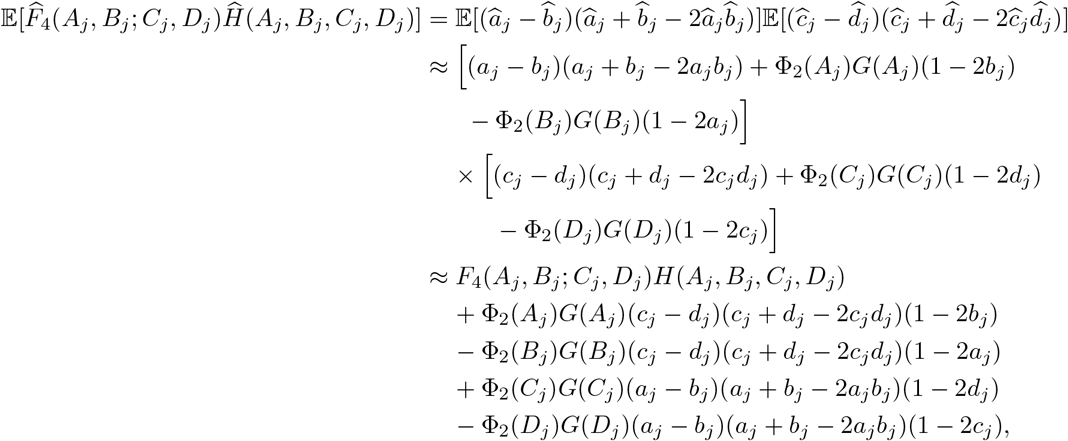

where we used the fact that Φ_2_(*A*_*j*_)Φ_2_(*C*_*j*_), Φ_2_(*A*_*j*_)Φ_2_(*D*_*j*_), Φ_2_(*B*_*j*_)Φ_2_(*C*_*j*_), and Φ_2_(*B*_*j*_)Φ_2_(*D*_*j*_) are negligible compared to Φ_2_(*A*_*j*_), Φ_2_(*B*_*j*_), Φ_2_(*C*_*j*_), and Φ_2_(*D*_*j*_) as an approximation. Putting it together, we have

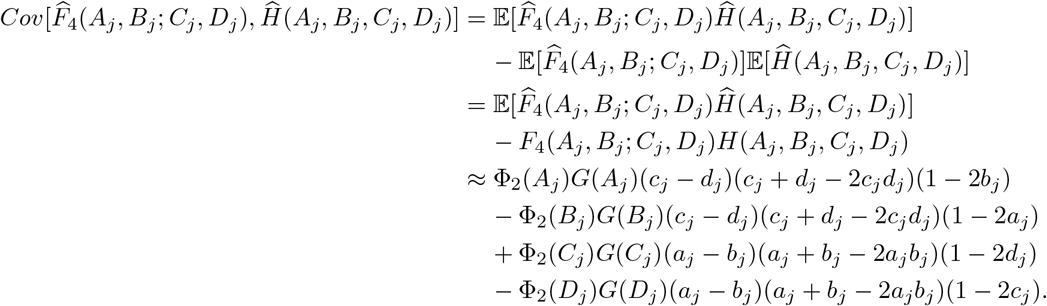

Given the assumption of independent loci, we have

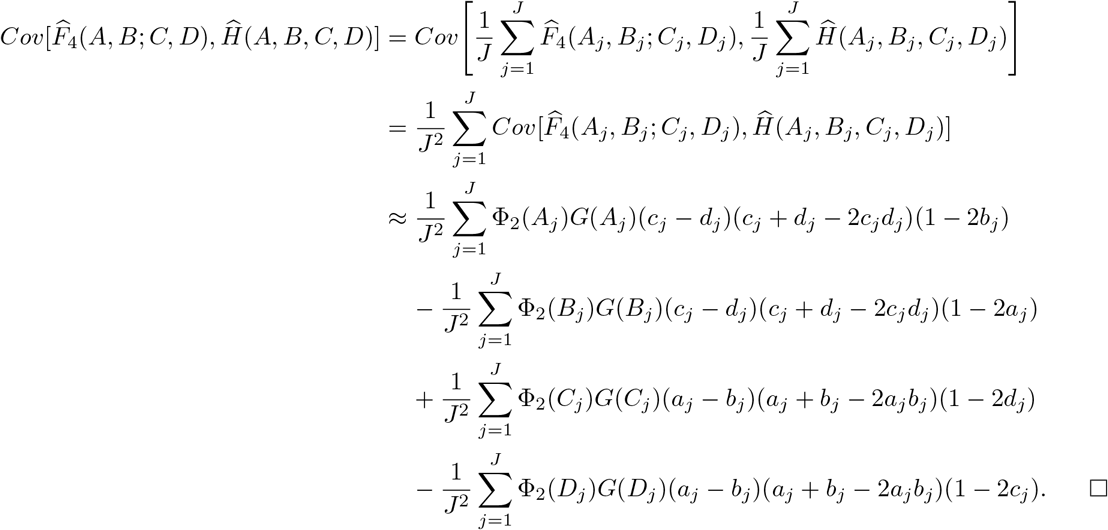

### Proposition 26.

Consider *J* polymorphic loci in populations *A*, *B*, *C*, and *D* with respective parametric reference allele frequencies *a*_*j*_, *b*_*j*_, *c*_*j*_, *d*_*j*_ ∈ (0, 1), and suppose we take a random sample of *N* (*P*_*j*_) individuals at locus *j* in population *P* ∈ {*A, B, C, D*}, some of which may be related or inbred. Moreover, assume that no individual is related to more than one other individual, which makes the terms Φ_3_(*P*_*j*_), Φ_4_(*P*_*j*_), Φ_2,2_(*P*_*j*_), Φ_2_(*P*_*j*_)^2^, Φ_2_(*A*_*j*_)Φ_2_(*B*_*j*_), Φ_2_(*A*_*j*_)Φ_2_(*C*_*j*_), Φ_2_(*A*_*j*_)Φ_2_(*D*_*j*_), Φ_2_(*B*_*j*_)Φ_2_(*C*_*j*_), Φ_2_(*B*_*j*_)Φ_2_(*D*_*j*_), and Φ_2_(*C*_*j*_)Φ_2_(*D*_*j*_) negligible to Φ_2_(*P*_*j*_). Based on this simplifying assumption, the approximately unbiased ratio estimator 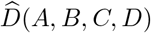 has approximate variance

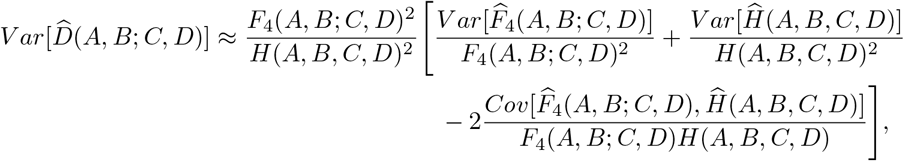

where the variances are

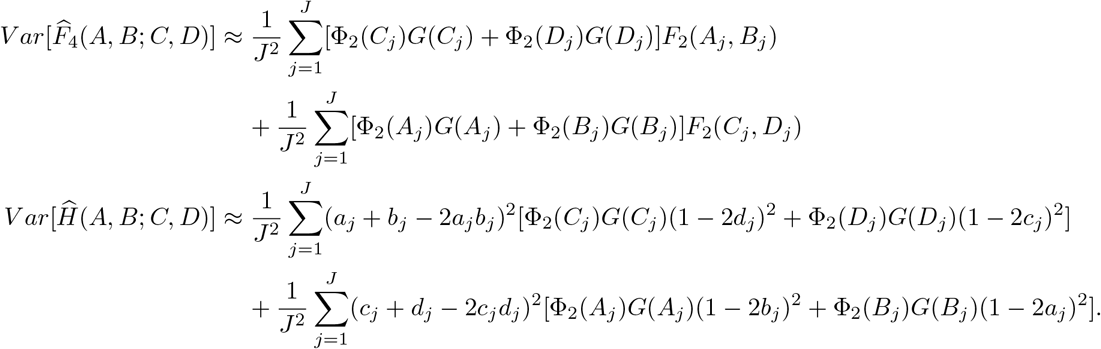

and the covariance is

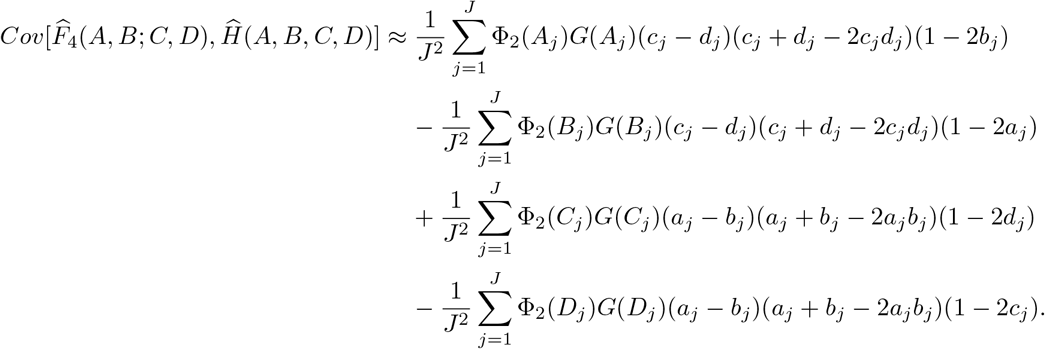

*Proof.* Recall that

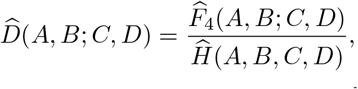

where 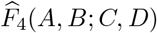 is an unbiased estimator for *F*_4_(*A, B*; *C, D*) and 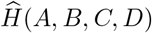 is an unbiased estimator of *H*(*A, B, C, D*). Assuming that 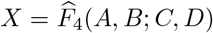 and 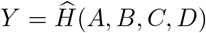, following the approximation in Wolter (2007) we have

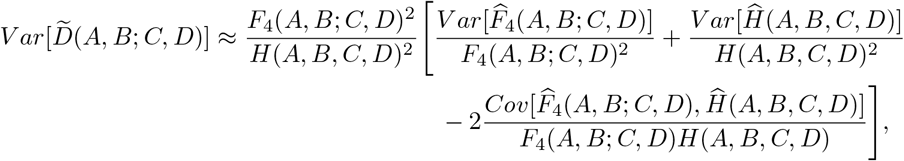

where 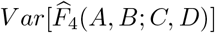 is given in Proposition 16, 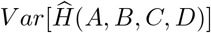 in Lemma 24, and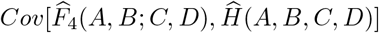 in Lemma 25.

## References

M. DeGiorgio and N. A. Rosenberg. An unbiased estimator of gene diversity in samples containing related individuals. Molecular Biology and Evolution, 26:501–512, 2009.

M. DeGiorgio, I. Jankovic, and N. A. Rosenberg. Unbiased estimation of gene diversity in samples containing related individuals: Exact variance and arbitrary ploidy. Genetics, 186:1367–1387, 2010.

D. A. R. Eaton and R. H. Ree. Inferring Phylogeny and Introgression using RADseq Data: An Example from Flowering Plants (Pedicularis: Orobanchaceae). Systematic Biology, 62:689–706, 2013.

M. P. Epstein, W. L. Duren, and M. Boehnke. Improved inference of relationship for pairs of individuals. American journal of human genetics, 67:1219–1231, 2000.

S. Gravel, B. M. Henn, R. N. Gutenkunst, A. R. Indap, G. T. Marth, A. G. Clark, F. Yu, R. A. Gibbs,, and C. D. Bustamante. Demographic history and rare allele sharing among human populations. Proceedings of the National Academy of Sciences, 108(29):11983–11988, 2011.

R. E. Green, J. Krause, A. W. Briggs, T. Maricic, U. Stenzel, M. Kircher, N. Patterson, H. Li, W. Zhai, M. H.-Y. Fritz, N. F. Hansen, E. Y. Durand, A.-S. Malaspinas, J. D. Jensen, T. Marques-Bonet, C. Alkan, K. Prüfer, M. Meyer, H. A. Burbano, J. M. Good, R. Schultz, A. Aximu-Petri, A. Butthof, B. Höber, B. Höffner, M. Siegemund, A. Weihmann, C. Nusbaum, E. S. Lander, C. Russ, N. Novod, J. Affourtit, M. Egholm, C. Verna, P. Rudan, D. Brajkovic, Ž. Kucan, I. Gušic, V. B. Doronichev, L. V. Golovanova, C. Lalueza-Fox, M. de la Rasilla, J. Fortea, A. Rosas, R. W. Schmitz, P. L. F. Johnson, E. E. Eichler, D. Falush, E. Birney, J. C. Mullikin, M. Slatkin, R. Nielsen, J. Kelso, M. Lachmann, D. Reich, and S. Pääbo. A draft sequence of the neandertal genome. Science, 328:710–722, 2010.

M. Hajdinjak, Q. Fu, A. Hübner, M. Petr, F. Mafessoni, S. Grote, P. Skoglund, V. Narasimham, H. Rougier, I. Crevecoeur, P. Semal, M. Soressi, S. Talamo, J.-J. Hublin, I. Gušić, Ž. Kućan, P. Rudan, L. V. Golo-vanova, V. B. Doronichev, C. Posth, J. Krause, P. Korlević, S. Nagel, B. Nickel, M. Slatkin, N. Patterson, D. Reich, K. Prüfer, M. Meyer, S. Pääbo, and J. Kelso. Reconstructing the genetic history of late nean-derthals. Nature, 555:652–656, 2018.

B. C. Haller and P. W. Messer. SLiM 3: Forward Genetic Simulations Beyond the Wright–Fisher Model. Molecular Biology and Evolution, 36:632–637, 2019.

A. M. Harris and M. DeGiorgio. An unbiased estimator of gene diversity with improved variance for samples containing related and inbred individuals of any ploidy. G3: Genes, Genomes, Genetics, 7:671–691, 2017a.

A. M. Harris and M. DeGiorgio. Admixture and ancestry inference from ancient and modern samples through measures of population genetic drift. Human Biology, 89:21–46, 2017b.

D. Huson, T. Klöpper, P. Lockhart, and M. Steel. Reconstruction of reticulate networks from gene trees. RECOMB 2005, 3500:233–249, 2005.

H. L. Kim, A. Ratan, G. H. Perry, A. Montenegro, W. Miller, and S. C. Schuster. Khoisan hunter-gatherers have been the largest population throughout most of modern-human demographic history. Nature communications, 5:5692–5692, 2014.

R. J. Kulathinal, L. S. Stevison, and M. A. F. Noor. The genomics of speciation in drosophila: Diversity, divergence, and introgression estimated using low-coverage genome sequencing. PLOS Genetics, 5: 5:e1000550, 2009.

K. Lange. Mathematical and Statistical Methods for Genetic Analysis. Springer, 2002.

J. Z. Li, D. M. Absher, H. Tang, A. M. Southwick, A. M. Casto, S. Ramachandran, H. M. Cann, G. S. Barsh, M. Feldman, L. L. Cavalli-Sforza, and R. M. Myers. Worldwide human relationships inferred from genome-wide patterns of variation. Science, 319:1100–1104, 2008.

S. H. Martin, J. W. Davey, and C. D. Jiggins. Evaluating the Use of ABBA–BABA Statistics to Locate Introgressed Loci. Molecular Biology and Evolution, 32:244–257, 2014.

S. McPeek, Mary, X. Wu, and C. Ober. Best linear unbiased allele-frequency estimation in complex pedigrees. Biometrics, 60:359–367, 2004.

L. Molinaro, F. Montinaro, B. Yelmen, D. Marnetto, D. M. Behar, T. Kivisild, and L. Pagani. West asian sources of the eurasian component in ethiopians: a reassessment. Scientific Reports, 9:18811, 2019.

P. Moorjani, K. Thangaraj, N. Patterson, M. Lipson, P.-R. Loh, P. Govindaraj, B. Berger, D. Reich, and Singh. Genetic evidence for recent population mixture in india. The American Journal of Human Genetics, 93:422–438, 2013.

L. Nei and A. K. Roychoudhury. Sampling variances of heterozygosity and genetic distance. 76:379–390, 1974.

M. Patterson, P. Moorjani, Y. Luo, S. Mallick, N. Rohland, Y. Zhan, T. Genschoreck, T. Webster, and D. Reich. Ancient admixture in human history. Genetics, 192:1065–1093, 2012.

B. A. Payseur and M. W. Nachman. Microsatellite variation and recombination rate in the human genome. Genetics, 156:1285–1298, 2000.

J. B. Pease and M. W. Hahn. Detection and Polarization of Introgression in a Five-Taxon Phylogeny. Systematic Biology, 64:651–662, 2015.

D. Reich, K. Thangaraj, N. Patterson, A. L. Price, and L. Singh. Reconstructing indian population history. Nature, 461:489 EP –, 2009.

D. Reich, N. Patterson, D. Campbell, A. Tandon, S. Mazieres, N. Ray, M. V. Parra, W. Rojas, C. Duque, N. Mesa, L. F. García, O. Triana, S. Blair, A. Maestre, J. C. Dib, C. M. Bravi, G. Bailliet, D. Corach, T. Hünemeier, M. C. Bortolini, F. M. Salzano, M. L. Petzl-Erler, V. Acuña-Alonzo, C. Aguilar-Salinas, S. Canizales-Quinteros, T. Tusié-Luna, L. Riba, M. Rodríguez-Cruz, M. Lopez-Alarcón, R. Coral-Vazquez, T. Canto-Cetina, I. Silva-Zolezzi, J. C. Fernandez-Lopez, A. V. Contreras, G. Jimenez-Sanchez, M. J. Gómez-Vázquez, J. Molina, Á. Carracedo, A. Salas, C. Gallo, G. Poletti, D. B. Witonsky, G. Alkorta-Aranburu, R. I. Sukernik, L. Osipova, S. A. Fedorova, R. Vasquez, M. Villena, C. Moreau, R. Barrantes, D. Pauls, L. Excoffier, G. Bedoya, F. Rothhammer, J.-M. Dugoujon, G. Larrouy, W. Klitz, D. Labuda, J. Kidd, K. Kidd, A. Di Rienzo, N. B. Freimer, A. L. Price, and A. Ruiz-Linares. Reconstructing native american population history. Nature, 488:370–374, 2012.

N. A. Rosenberg. Standardized subsets of the hgdp-ceph human genome diversity cell line panel, accounting for atypical and duplicated samples and pairs of close relatives. Annals of Human Genetics, 70:841–847, 2006.

A. Scally and R. Durbin. Revising the human mutation rate: implications for understanding human evolution. Nature Reviews Genetics, 13:745, 2012.

S. Soraggi, C. Wiuf, and A. Albrechtsen. Powerful inference with the *D*-statistic on low-coverage whole-genome data. G3: Genes, Genomes, Genetics, 8:551–566, 2018.

N. Takahata. Allelic genealogy and human evolution. Molecular Biology and Evolution, 10:2–22, 1993.

The 1000 Genomes Project Consortium. A global reference for human genetic variation. Nature, 526:68–74, 2015.

D. A. Turissini and D. R. Matute. Fine scale mapping of genomic introgressions within the drosophila yakuba clade. PLOS Genetics, 13:1–40, 2017.

R. S. Waples and E. C. Anderson. Purging putative siblings from population genetic data sets: a cautionary view. Molecular Ecology, 26:1211–1224, 2017.

B. S. Weir. Sampling properties of gene diversity. Plant population genetics, breeding and genetic resources, pages 23–42, 1989.

B. S. Weir and C. C. Cockerham. Estimating *F*-statistics for the analysis of population structure. Evolution, 38:1358–1370, 1984.

K. M. Wolter. Introduction to variance estimation. Springer, New York, NY, 2nd edition, 2007.

Y. Zheng and A. Janke. Gene flow analysis method, the *D*-statistic, is robust in a wide parameter space. BMC bioinformatics, 19:10–10, 2018.

