## Supplement for "Properties and unbiased estimation of *F*- and *D*-statistics in samples containing related and inbred individuals"

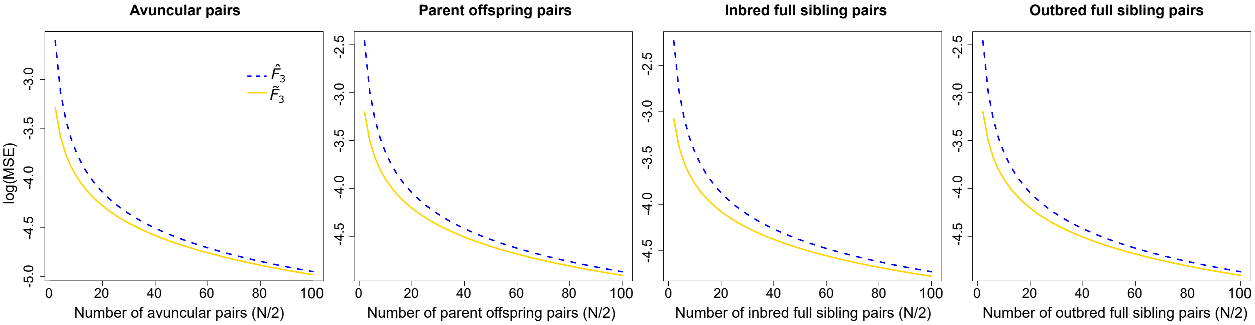


Figure S1: Mean squared error theoretically calculated for $\widehat{F}_3(A;B,C)$ and $\widetilde{F}_3(A;B,C)$ across different sample sizes or related pairs of individuals, including avuncular relationships, parent-offspring relationships, inbred full siblings, and outbred full siblings. The number of sampled individuals ranges from two to 100 with the number of relative pairs equaling half the total sampled, all computed using $J = 20$ loci. The true value of $F_3(A;B,C)$ is 0.033.

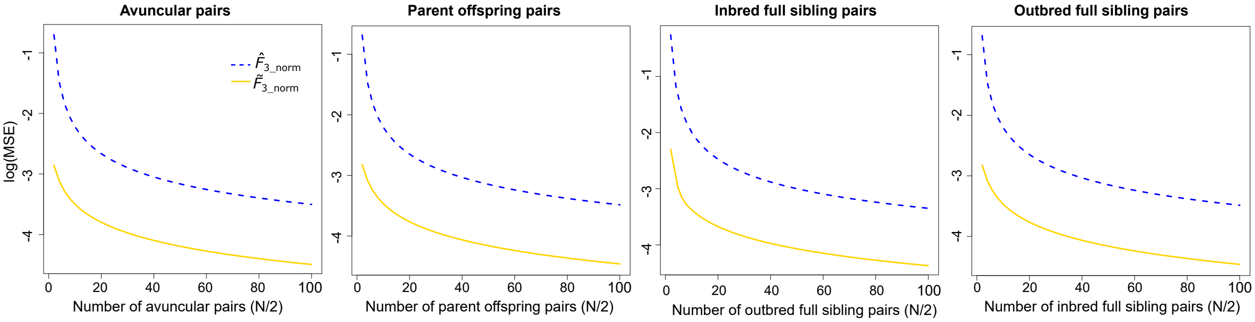


Figure S2: Mean squared error theoretically calculated for normalized $\widehat{F}_3(A;B,C\,|\,A)$ and $\widetilde{F}_3(A;B,C\,|\,A)$ across different sample sizes or related pairs of individuals, including avuncular relationships, parent-offspring relationships, inbred full siblings, and outbred full siblings. The number of sampled individuals ranges from two to 100 with the number of relative pairs equaling half the total sampled, all computed using $J = 20$ loci. The true value of normalized $F_3(A;B,C\,|\,A)$ is 0.116.

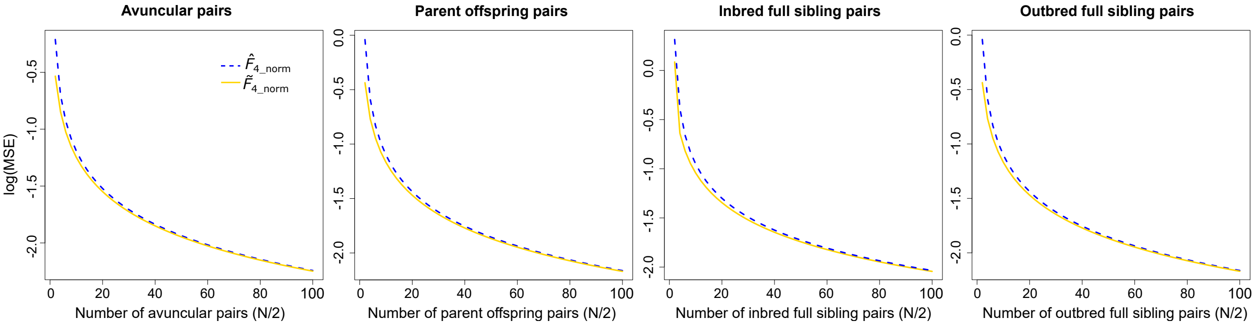


Figure S3: Mean squared error theoretically calculated for $\widehat{F}_4(A, B; C, D \mid A)$ and $\widetilde{F}_4(A, B; C, D \mid A)$ across different sample sizes or related pairs of individuals, including avuncular relationships, parent-offspring relationships, inbred full siblings, and outbred full siblings. The number of sampled individuals ranges from two to 100 with the number of relative pairs equaling half the total sampled, all computed using $J = 20$ loci. The true value of normalized $F_4(A, B; C, D \mid A)$ is 0.052.

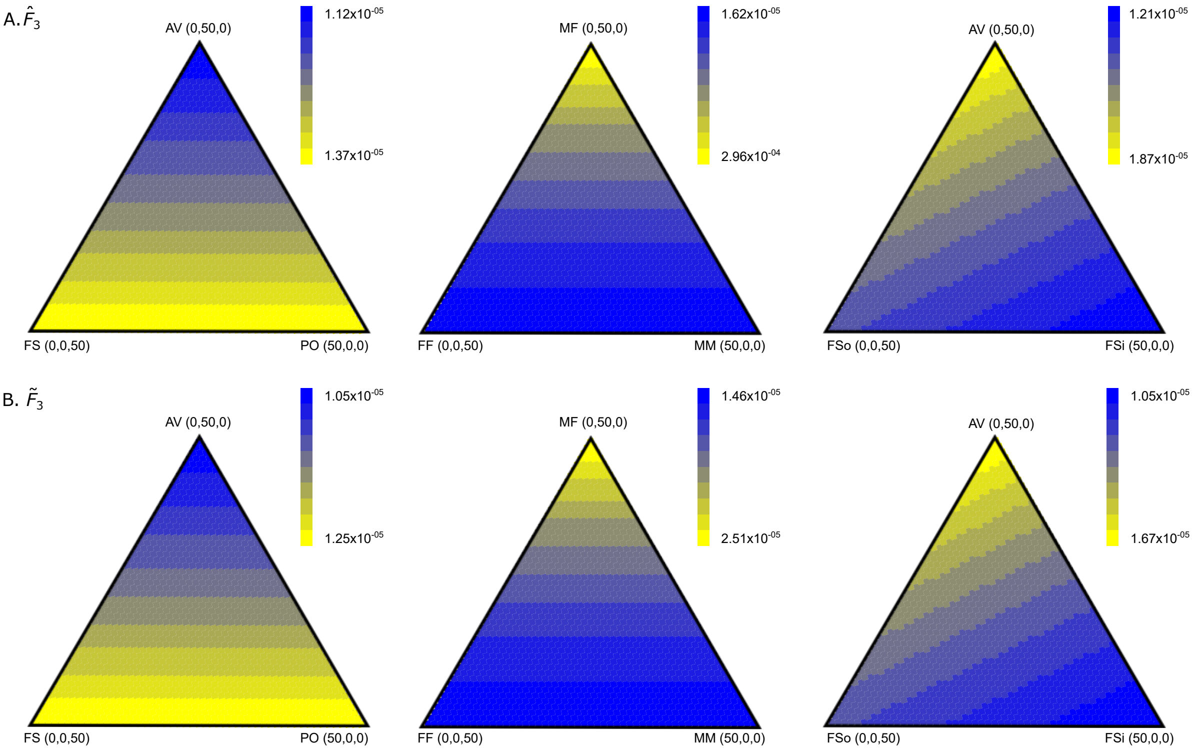


Figure S4: Theoretically calculated MSE of $\widehat{F_3}(A;B,C)$ and $\widetilde{F_3}(A;B,C)$ when including relatives or inbred individuals for $J = 20$ loci. The MSE is estimated for instances when samples of 100 individuals include individuals related to exactly one other in the sample. The first column shows MSE for samples with different combinations of parent-offspring (PO), full sibling (FS), and avuncular (AV) relationships, the second includes full siblings that are male-male (MM), male-female (MF) and female-female (MF). The last column includes AV relationships as well as inbred (FSi) and outbred (FSo) full siblings. The true value of $F_3(A;B,C)$ is 0.033.

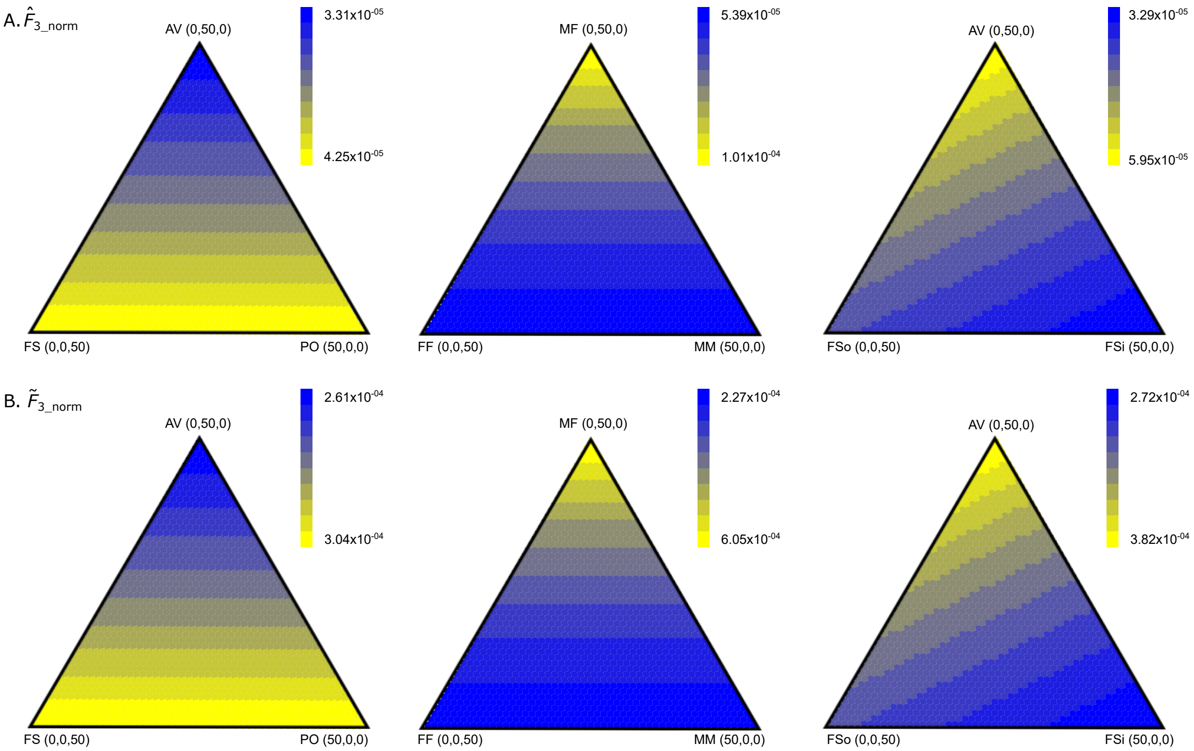


Figure S5: Theoretically calculated MSE of normalized $\widehat{F}_3(A; B, C \mid A)$ and normalized $\widetilde{F}_3(A; B, C \mid A)$ when including relatives or inbred individuals for $J = 20$ loci. The MSE is estimated for instances when samples of 100 individuals include individuals related to exactly one other in the sample. The first column shows MSE for samples with different combinations of parent-offspring (PO), full sibling (FS), and avuncular (AV) relationships, the second includes full siblings that are male-male (MM), male-female (MF) and female-female (MF). The last column includes AV relationships as well as inbred (FSi) and outbred (FSo) full siblings. The true value of normalized $F_3(A; B, C \mid A)$ is 0.116.

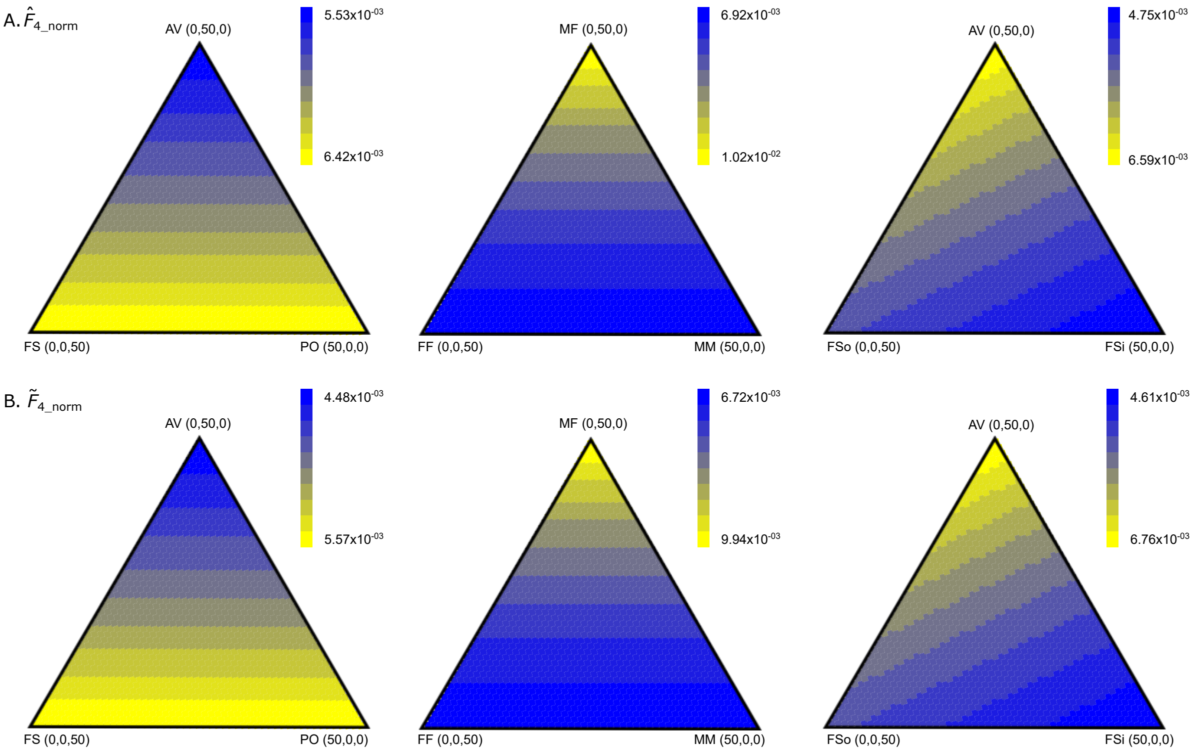


Figure S6: Theoretically calculated MSE of normalized $\widehat{F}_4(A,B;C,D\,|\,A)$ and normalized $\widetilde{F}_4(A,B;C,D\,|\,A)$ when including relatives or inbred individuals for $J = 20$ loci. The MSE is estimated for instances when samples of 100 individuals include individuals related to exactly one other in the sample. The first column shows MSE for samples with different combinations of parent-offspring (PO), full sibling (FS), and avuncular (AV) relationships, the second includes full siblings that are male-male (MM), male-female (MF) and female-female (MF). The last column includes AV relationships as well as inbred (FSi) and outbred (FSo) full siblings. The true value of normalized $F_4(A,B;C,D\,|\,A)$ is 0.052.

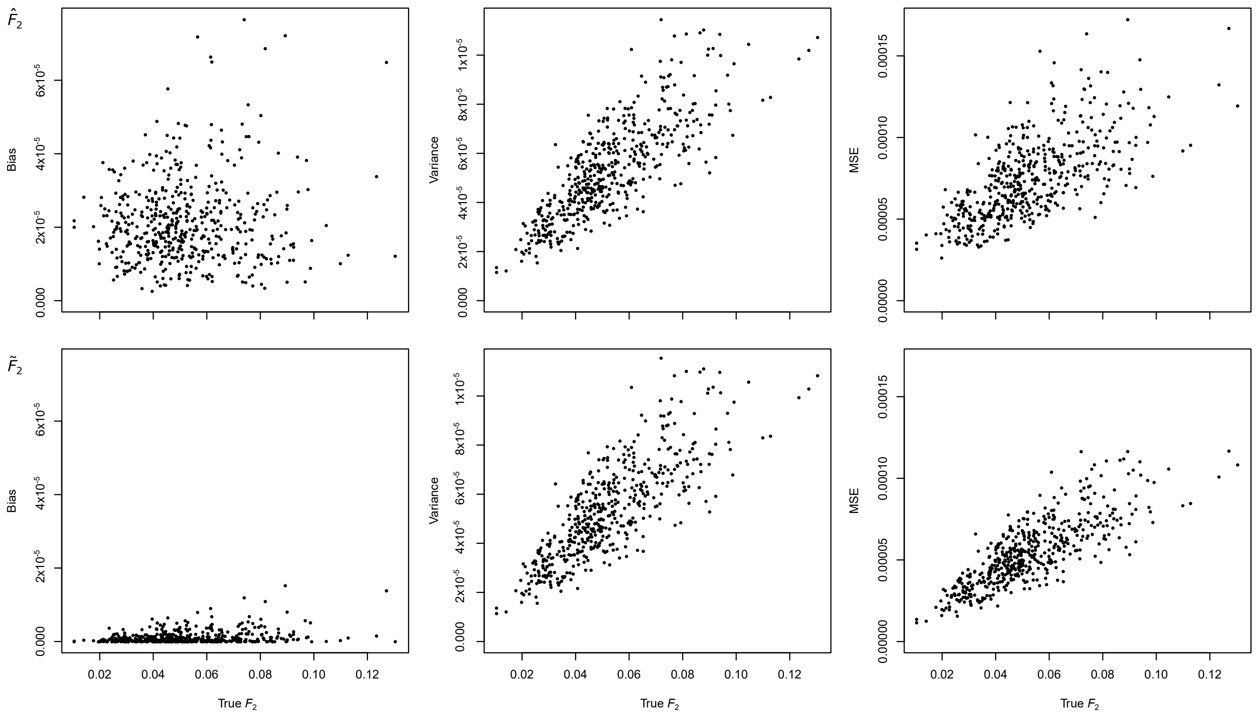


Figure S7: Comparison of squared bias, variance, and MSE for $\widehat{F}_2(A,B)$ and $\widetilde{F}_2(A,B)$ from simulated data including 60 parent offspring relative pairs. Each estimate was computed from $J = 20$ randomly sampled loci using $A =$ CEU and $B =$ YRI.

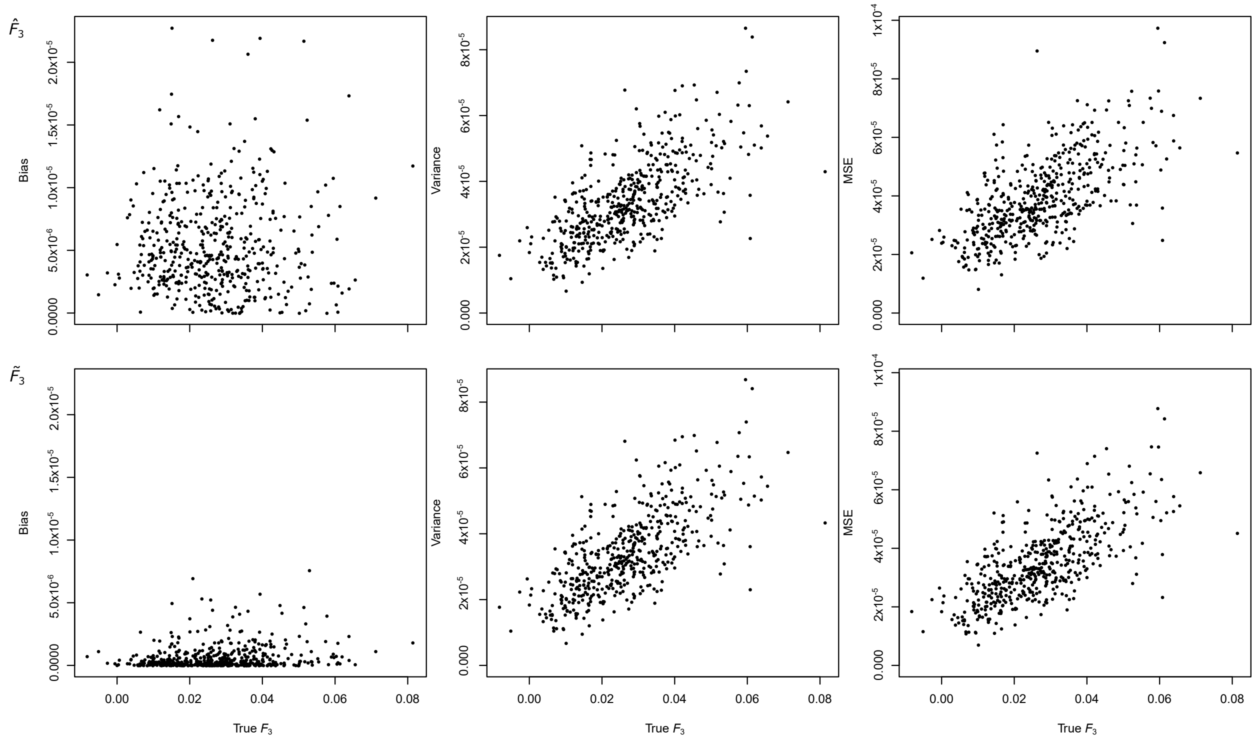


Figure S8: Comparison of squared bias, variance, and MSE for $\widehat{F}_3(A;B,C)$ and $\widetilde{F}_3(A;B,C)$ from simulated data including 60 parent offspring relative pairs. Each estimate was computed using $J = 20$ randomly sampled loci using $A =$ JPT, $B =$ CEU and $C =$ YRI.

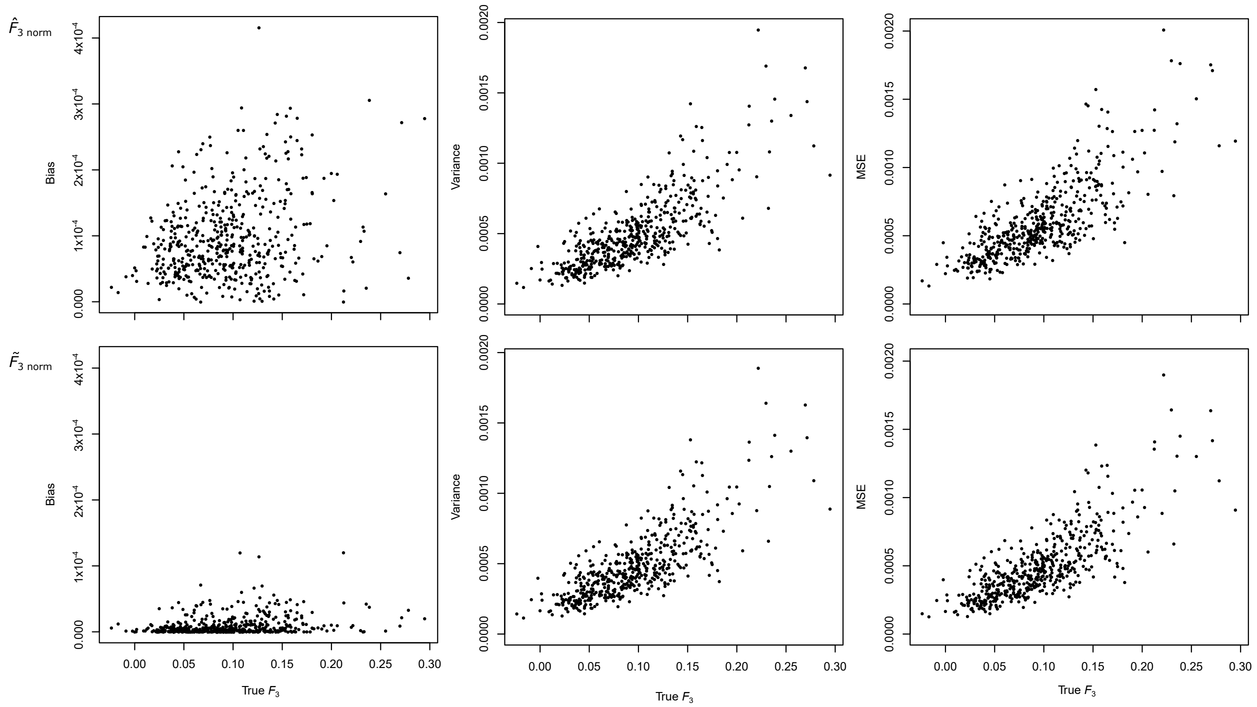


Figure S9: Comparison of squared bias, variance, and MSE for normalized $\widehat{F}_3(A;B,C\,|\,A)$ and $\widetilde{F}_3(A;B,C\,|\,A)$ from simulated data including 60 relative pairs. Each estimate was computed using $J = 20$ randomly sampled loci using $A =$ JPT, $B =$ CEU and $C =$ YRI.

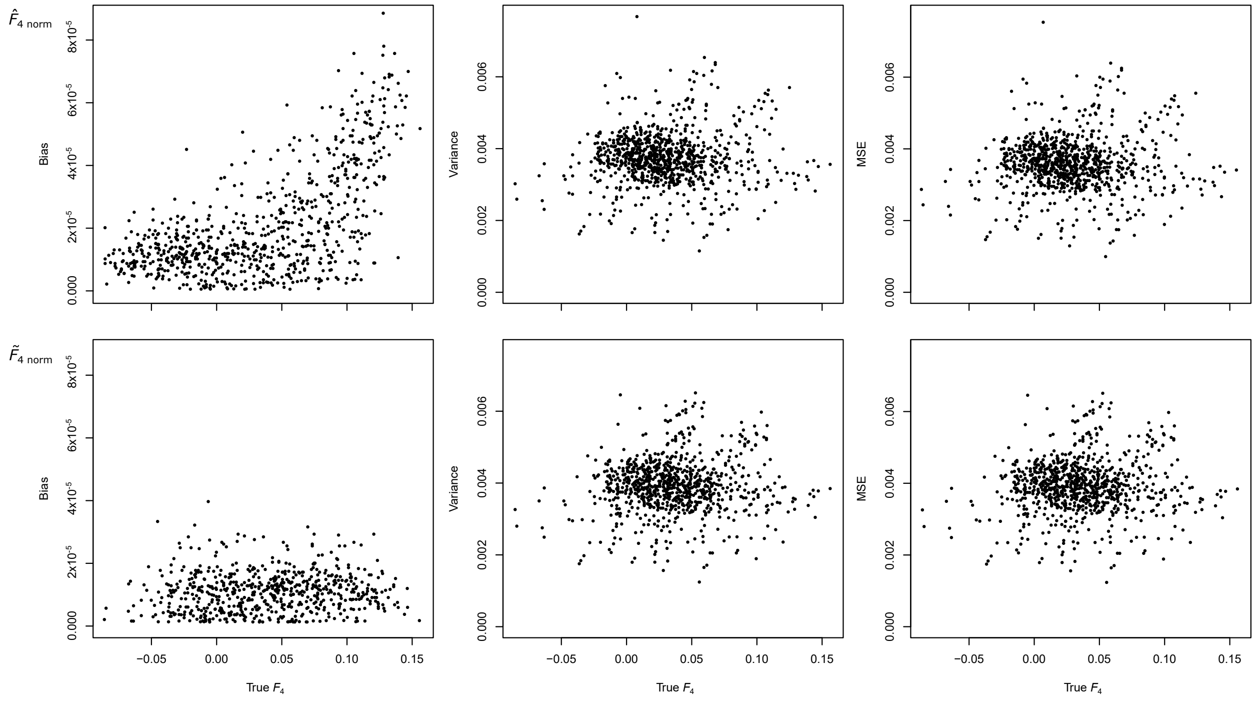


Figure S10: Comparison of squared bias, variance, and MSE for normalized $\widehat{F}_4(A, B; C, D \mid A)$ and $\widetilde{F}_4(A, B; C, D \mid A)$ from simulated data including 60 relative pairs. Each estimate was computed using $J = 20$ randomly sampled loci using $A =$ YRI, $B =$ CEU, $C =$ JPT, and $D =$ GIH.

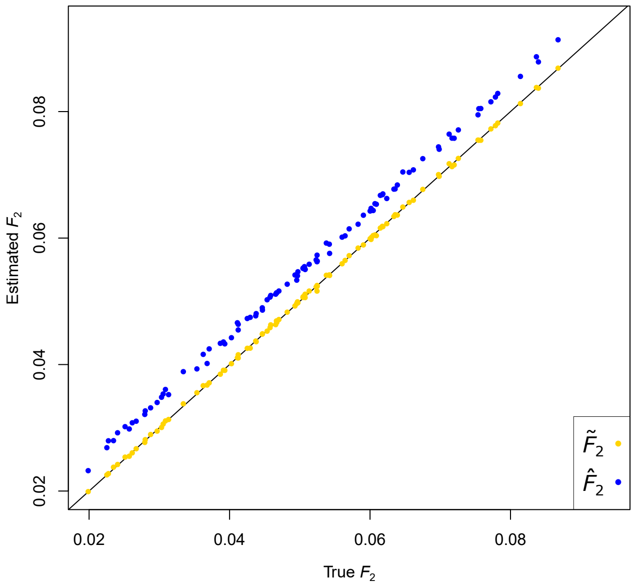


Figure S11: Comparison of true $F_2(A, B)$ to estimated $\widetilde{F}_2(A, B)$ and $\widehat{F}_2(A, B)$. Each dot represents the mean of 1000 simulations of parent offspring pairs used to compute $\widetilde{F}_2(A, B)$ and $\widehat{F}_2(A, B)$. Each simulation contains 50 parent offspring pairs.

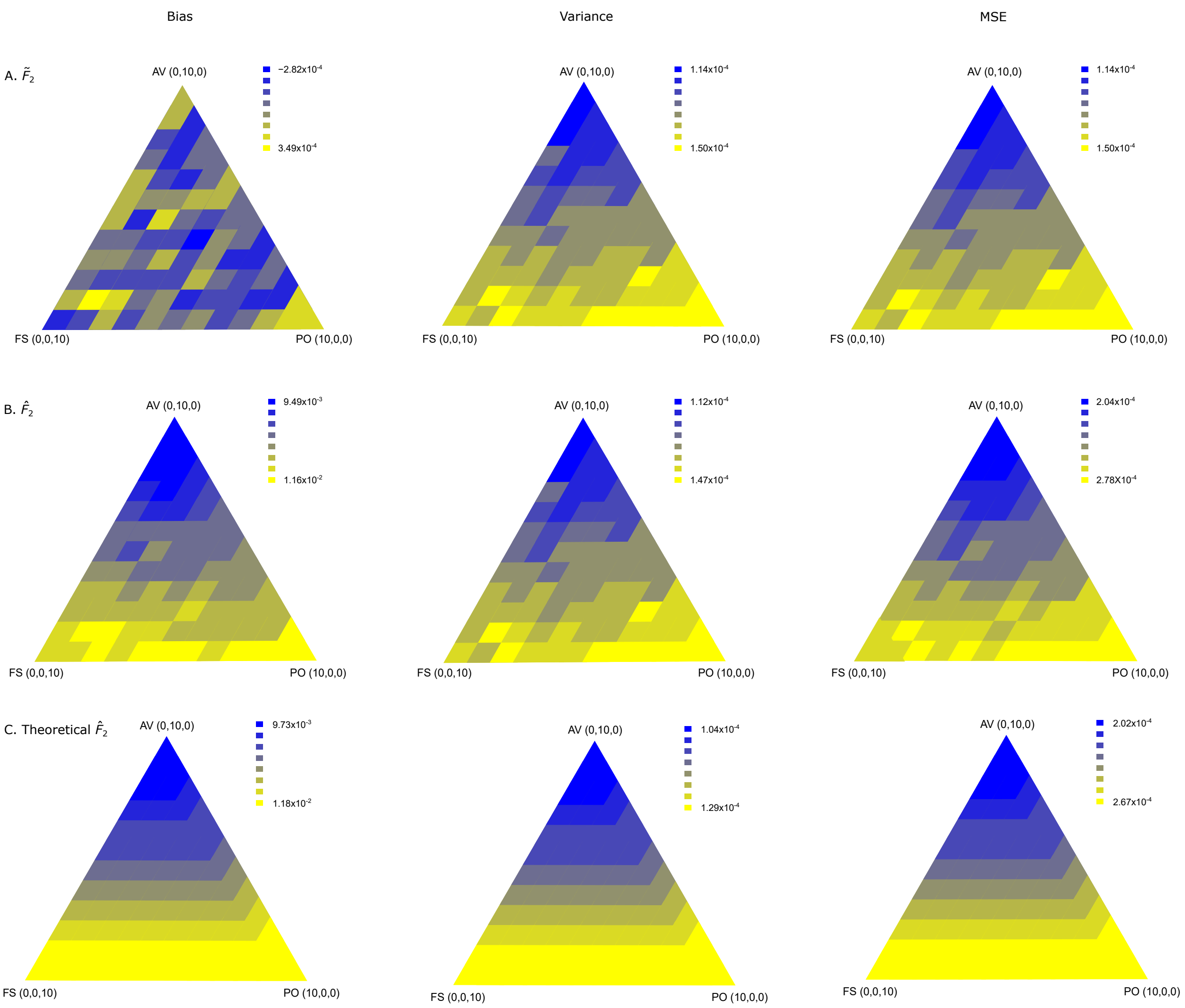


Figure S12: Theoretical vs. simulated $F_2(A, B)$ bias, variance, and MSE when including various combinations of parent-offspring, avuncular and outbred full-sibling pairs. (Top row) Simulations of relative pairs used to compute bias, variance, and MSE of $\widetilde{F}_2(A, B)$. (Middle row) Simulations of relative pairs used to compute bias, variance, and MSE of $\widehat{F}_2(A, B)$. (Bottom) Theoretically computed bias, variance, and MSE for $\widehat{F}_2(A, B)$.The true value of $F_2(A, B)$ is 0.071, computed for $J = 20$ loci.

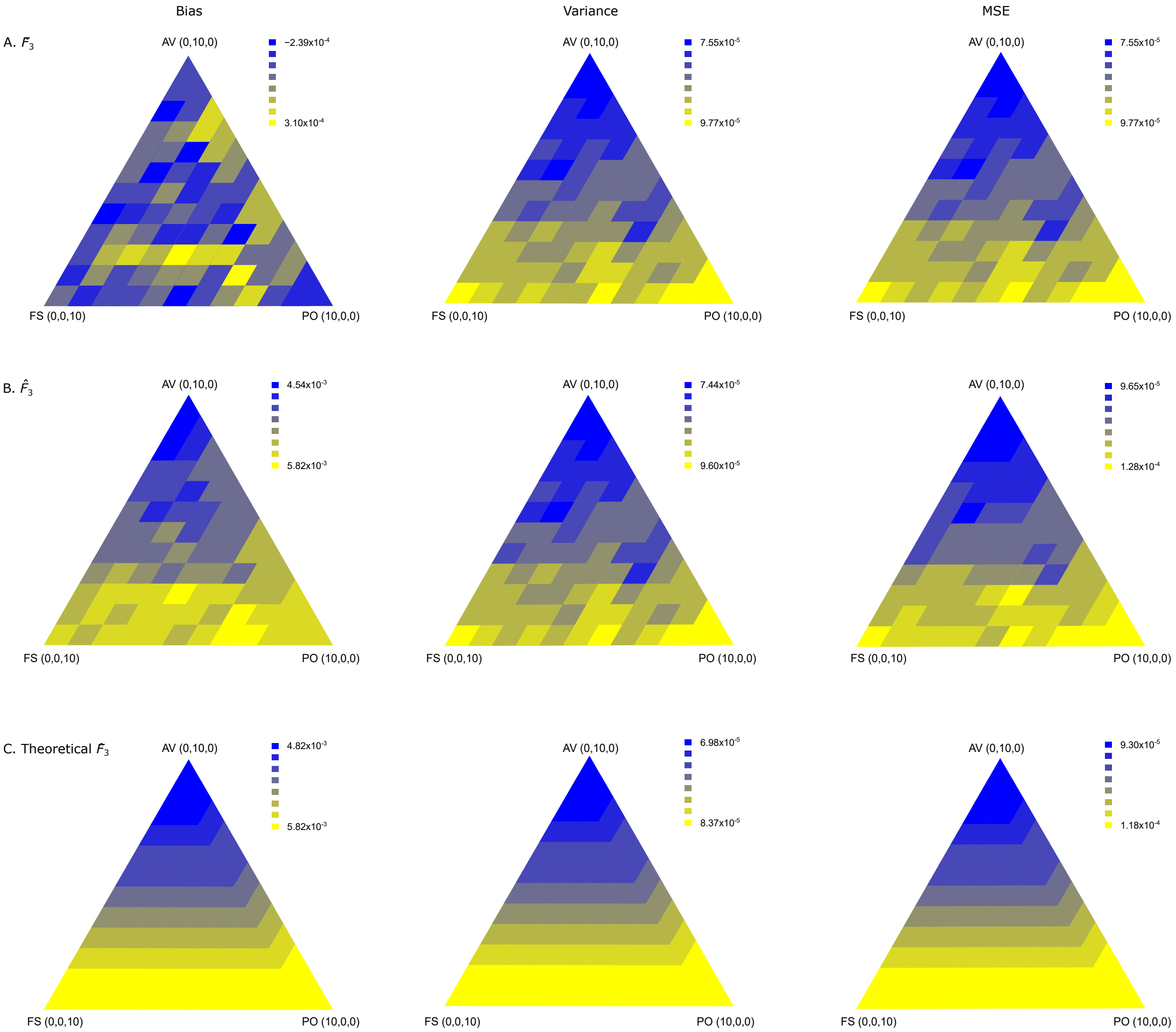


Figure S13: Theoretical vs. simulated $F_3(A;B,C)$ bias, variance, and MSE when including various combinations of parent-offspring, avuncular and outbred full-sibling pairs. (Top row) Simulations of relative pairs used to compute bias, variance, and MSE of $\widetilde{F_3}(A;B,C)$. (Middle row) Simulations of relative pairs used to compute bias, variance, and MSE of $\widehat{F_3}(A;B,C)$. (Bottom) Theoretically computed bias, variance, and MSE for $\widehat{F_3}(A;B,C)$. The true value of $F_3(A;B,C)$ is 0.033, computed for $J = 20$ loci.

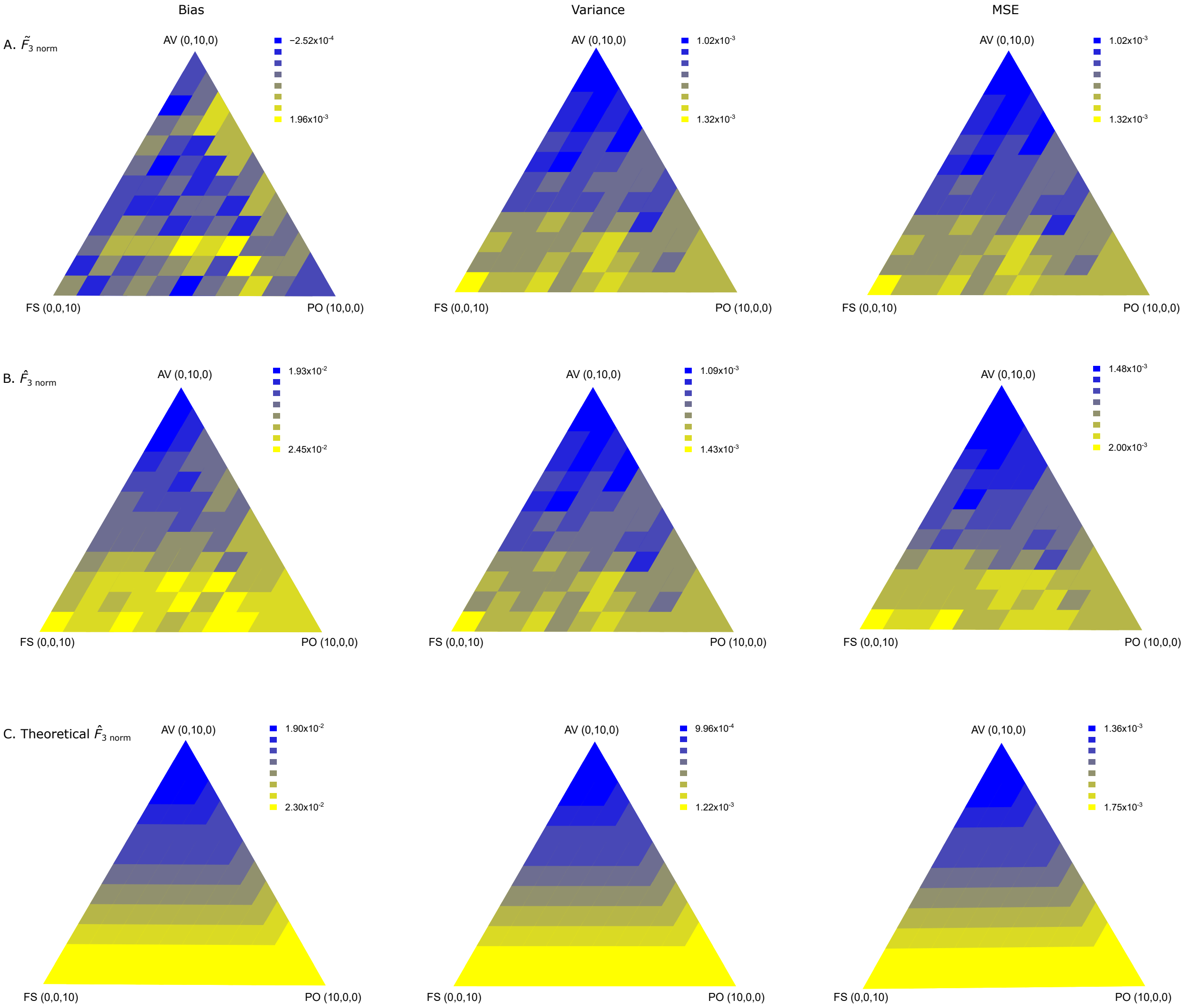


Figure S14: Theoretical vs. simulated normalized $F_3(A;B,C\,|\,A)$ bias, variance, and MSE when including various combinations of parent-offspring, avuncular and outbred full-sibling pairs. (Top row) Simulations of relative pairs used to compute bias, variance, and MSE of normalized $\widetilde{F}_3(A;B,C\,|\,A)$. (Middle row) Simulations of relative pairs used to compute bias, variance, and MSE of normalized $\widehat{F}_3(A;B,C\,|\,A)$. (Bottom) Theoretically computed bias, variance, and MSE for normalized $\widehat{F}_3(A;B,C\,|\,A)$. The true value of normalized $F_3(A;B,C\,|\,A)$ is 0.116, computed for $J = 20$ loci.

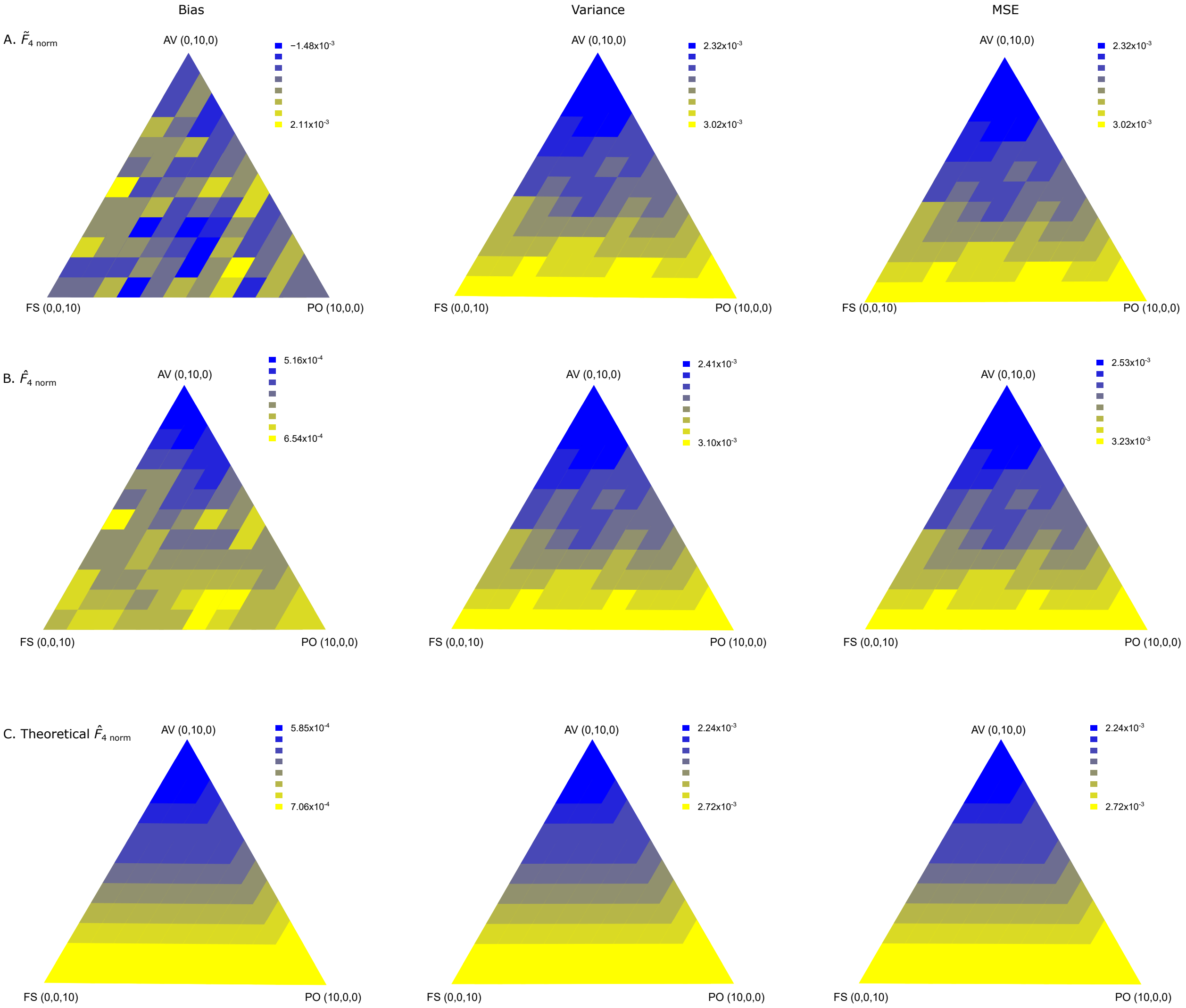


Figure S15: Theoretical vs. simulated normalized $F_4(A, B; C, D \mid A)$ bias, variance, and MSE when including various combinations of parent-offspring, avuncular and outbred full-sibling pairs. (Top row) Simulations of relative pairs used to compute bias, variance, and MSE of normalized $\widetilde{F}_4(A, B; C, D \mid A)$. (Middle row) Simulations of relative pairs used to compute bias, variance, and MSE of normalized $\widehat{F}_4(A, B; C, D \mid A)$. (Bottom) Theoretically computed bias, variance, and MSE for normalized $\widehat{F}_4(A, B; C, D \mid A)$. The true value of normalized $F_4(A, B; C, D \mid A)$ is 0.052, computed for $J = 20$ loci.
